# A suite of engineered mice for interrogating psychedelic drug actions

**DOI:** 10.1101/2023.09.25.559347

**Authors:** Yi-Ting Chiu, Ariel Y. Deutch, Wei Wang, Gavin P Schmitz, Karen Lu Huang, D. Dewran Kocak, Pierre Llorach, Kasey Bowyer, Bei Liu, Noah Sciaky, Kunjie Hua, Chongguang Chen, Sarah E. Mott, Jesse Niehaus, Jeffrey F. DiBerto, Justin English, Jessica J. Walsh, Grégory Scherrer, Melissa A Herman, Zhuhao Wu, William C Wetsel, Bryan L Roth

## Abstract

Psychedelic drugs like lysergic acid diethylamide (LSD) and psilocybin have emerged as potentially transformative therapeutics for many neuropsychiatric diseases, including depression, anxiety, post-traumatic stress disorder, migraine, and cluster headaches. LSD and psilocybin exert their psychedelic effects via activation of the 5-hydroxytryptamine 2A receptor (HTR2A). Here we provide a suite of engineered mice useful for clarifying the role of HTR2A and HTR2A-expressing neurons in psychedelic drug actions. We first generated *Htr2a*-EGFP-CT-IRES-CreERT2 mice (CT:C-terminus) to independently identify both HTR2A-EGFP-CT receptors and HTR2A-containing cells thereby providing a detailed anatomical map of HTR2A and identifying cell types that express HTR2A. We also generated a humanized *Htr2a* mouse line and an additional constitutive *Htr2A*-Cre mouse line. Psychedelics induced a variety of known behavioral changes in our mice validating their utility for behavioral studies. Finally, electrophysiology studies revealed that extracellular 5-HT elicited a HTR2A-mediated robust increase in firing of genetically-identified pyramidal neurons--consistent with a plasma membrane localization and mode of action. These mouse lines represent invaluable tools for elucidating the molecular, cellular, pharmacological, physiological, behavioral, and other actions of psychedelic drugs *in vivo*.

## INTRODUCTION

Serotonin (5-hydroxytryptamine; 5-HT) is a biogenic amine neurotransmitter essential for modulating mood, perception, cognition, pain, feeding, and a variety of other functions in the central nervous system and periphery (Berger et al., 2009; Barnes et al., 2021). To mediate these actions a large family of 14 distinct receptors that are heterogeneously distributed throughout the body have evolved (Berger et al., 2009; Barnes et al., 2021; Gumpper et al., 2023). Except for the 5-HT3 receptor, all mammalian 5-HT receptors are G-protein-coupled receptors. Although the cell bodies of 5-HT-producing neurons are exclusively localized to brainstem sites, they project widely throughout the brain and spinal cord to regulate broad neural targets through 5-HT receptors (Hillarp et al., 1966; Steinbusch et al., 2021; Jacobs et al., 1992).

Of the various 5-HT receptors, the 5-hydroxytryptamine 2A receptor (HTR2A) has recently received the most attention, in part due to the apparent involvement of the HTR2A in the pathogenesis of several psychiatric disorders, including schizophrenia, depression, Parkinson’s Disease psychosis, and anxiety disorders (Meltzer et al., 2013; Barnes et al., 2021; McClure-Begley et al., 2022). Moreover, the HTR2A has been the focus of intense interest because this receptor is the primary molecular target for psychedelic drugs (Glennon et al., 1984), including lysergic acid diethylamide (LSD), psilocybin, *N,N’*-dimethyltryptamine, and mescaline (Peroutka et al., 1981; Gonzalez-Maeso et al., 2008; Keiser et al., 2009b; Kometer et al., 2013; Preller et al., 2018; Vollenweider et al., 1998). The HTR2A also has high affinity for most second generation (atypical) antipsychotic drugs (Meltzer et al., 1989), as well as many typical and atypical antidepressants (Palvimaki et al., 1996; Roth et al., 2004). At the cellular level, HTR2A agonists promote Gαq protein coupling (Roth et al., 1984; Kim et al., 2020) and β-arrestin recruitment (Gray et al., 2003; Wacker et al., 2013), leading to receptor internalization (Berry et al., 1996) and the activation of many downstream signaling pathways (Gumpper et al., 2023).

The HTR2A was first identified by radioligand binding in 1978 (Leysen et al., 1978), was characterized as a membrane protein in 1985 (Wouters et al., 1985), and its encoding gene, *Htr2a*, was cloned in 1988 (Pritchett et al., 1988). The structure of *Htr2a* was characterized further by the Chen and Shih groups for human *HTR2A* and Toth group for murine *Htr2a*, who identified various introns, exons, and promoters (Chen et al., 1992; Shih et al., 1996; Toth, 1996). The sequence of the *HTR2A* was generally conserved between species with the rodent *Htr2a* and human *HTR2A* genes sharing 91.5% deduced amino acid identity with a single amino acid difference in the binding pocket that alters the binding properties of many psychedelic (Kim et al., 2020) and non-psychedelic 5-HT2A agonists (Johnson et al., 1994).

Initial receptor autoradiographic (Pazos et al., 1985; Pazos et al., 1987; Roth et al., 1987) and *in situ* hybridization studies (Mengod et al., 1990; Pompeiano et al., 1994; (Wright et al., 1995) identified a heterogeneous distribution of HTR2A in the rat and human brains with high expression in the deep layers of the human and rodent cerebral cortex. Subsequent studies with HTR2A antibodies (Willins et al., 1997a; Jakab et al., 1998) suggested the HTR2A was localized primarily to pyramidal neurons with expression in somas, apical dendrites (Willins et al., 1997a), and post-synaptic densities (Miner et al., 2003; Xia et al., 2003b) as well as occasional interneurons (Willins et al., 1997a; Jakab and Goldman-Rakic, 1998). Moreover, the HTR2A was expressed in subcortical regions, particularly in telencephalic sites (Bubser et al., 2001; Cornea-Hebert et al., 1999). However, immunohistochemical studies of HTR2A location were conflicting and inconsistent (for example, see (McDonald and Mascagni, 2007) presumably because almost none of the HTR2A antibodies used have been rigorously characterized. Only one antibody--which was suboptimal for routine studies--has been validated in wild-type (WT) and *Htr2a* knock-out mice (Magalhaes et al., 2010b; Yadav et al., 2011b). Autoradiographic studies yielded somewhat more consistent findings, but most suffered from relatively low resolution as well as specificity issues related to radioligands (Pazos et al., 1985;). Similarly, *in-situ* hybridization revealed only mRNA levels, which do not necessarily correspond with protein expression. Thus, there is a pressing need for tools to precisely study HTR2A protein distribution and function.

Here we provide as a resource to the scientific community, a suite of *Htr2a* reporter mice generated by CRISPR-mediated recombination at the endogenous *Htr2a* locus. We show this resource facilitates the identification and targeting of the HTR2A *in vivo* and provides potential opportunities for modulating the activity of HTR2A-expressing neurons with a variety of chemogenetic (Armbruster et al., 2007) and optogenetic (Boyden et al., 2005) actuators. We further demonstrate the utility of these mice for electrophysiological, biochemical, and genetic studies of HTR2A-identified neurons. Given the essential role of the HTR2A in mediating the action of all known psychedelic and many non-psychedelic therapeutics, this resource will be of significant value to the scientific research community.

## RESULTS

### Generation and characterization of the Htr2a-EGFP-CT reporter mouse line

Prior to initiating our studies, we compared available *Htr2a* reporter mice that were developed as part of the GENSAT initiative (Gong et al., 2003; Gerfen et al., 2013). Two of these mouse strains, Tg(*Htr2a*-Cre)KM207Gsat/Mmucd and Tg(*Htr2a*-Cre)KM208Gsat/Mmucd, have reported distributions of GFP following Cre-mediated recombination with reporter mice (http://www.gensat.org/cre.jsp). As shown at similar coronal sections in Supplementary Fig S1A and Fig S1B, these mice display distinct patterns of GFP expression. However, neither corresponds to the reported patterns of endogenous receptor binding (Pazos et al., 1985; Roth et al., 1987; Lopez-Gimenez et al., 1997; Lopez-Gimenez et al., 2013), or for *Htr2a* mRNA (Mengood et al., 1990; Pompeiano et al., 1994) (Wright et al., 1995) or protein expression in rodents (Bubser et al., 2001; Willins et al., 1997a) (see Suppl. Fig S1C to S1E). We note an absence of expression in the striatum in Tg(*Htr2a*-Cre)KM207Gsat/Mmucd and Tg(*Htr2a*-Cre)KM208Gsat/Mmucd mice (Suppl. Fig S1A and S1B) even though many prior studies have demonstrated significant HTR2A expression in this brain region (for example, see Bubser et al., 2001 and (Roth et al., 1987)). In addition, recent mapping studies by the Allen Institute for Brain Science disclosed abundant *Htr2a* mRNA expression in murine cortex and in a patch-like distribution in striatum (https://mouse.brain-map.org/experiment/show/81671344; Suppl. Fig S1C and S1G), consistent with the observation that HTR2A is localized to striosomes in rodent (Lopez-Gimenez et al., 1997) brains (FigS1D) and human (Lopez-Gimenez et al., 1999).

*Htr2a*-EGFP-CT-IRES-CreERT2 (*Htr2a*^EGFP-CreERT2^) mice were generated using CRISPR/Cas9 genome editing of exon 3 of *Htr2a* locus on chromosome 14 and a construct spanning nucleotides 74640840-74709494 to insert EGFP-CT followed by an IRES (internal ribosome entry site) and an estrogen-responsive Cre-recombinase (CreERT2) (Fig 1A). The mouse line has two modifications. *First*, a cDNA encoding EGFP was inserted into the C-terminus (CT) of the receptor, flanked on both ends by a serine- and glycine-rich linker. The insertion site of EGFP was after residue 452, yielding a construct we term *Htr2a*-EGFP-CT. Parenthetically, this insertion site was identified based on our previous studies (Abbas et al., 2009; Xia et al., 2003a; Xia et al., 2003b) and it was positioned between the 8^th^ helix and the C-terminal PDZ binding motif of the HTR2A. This location has been demonstrated not to affect expression, function, targeting, or trafficking of the receptor in neurons or transfected HEK293 cells (Abbas et al., 2009; Xia et al., 2003a; Xia et al., 2003b). *Second*, the IRES-CreERT2 sequence directly followed the coding sequence, and it preceded the 3’ UTR of *Htr2a*. The expression of CreERT2 recombinase under control of the endogenous *Htr2a* promoter provides a genetic manipulation platform to leverage Cre-dependent reporter mice or AAV viruses for specifically expressing reporter or actuator proteins in HTR2A containing cells.

**Fig 1.**
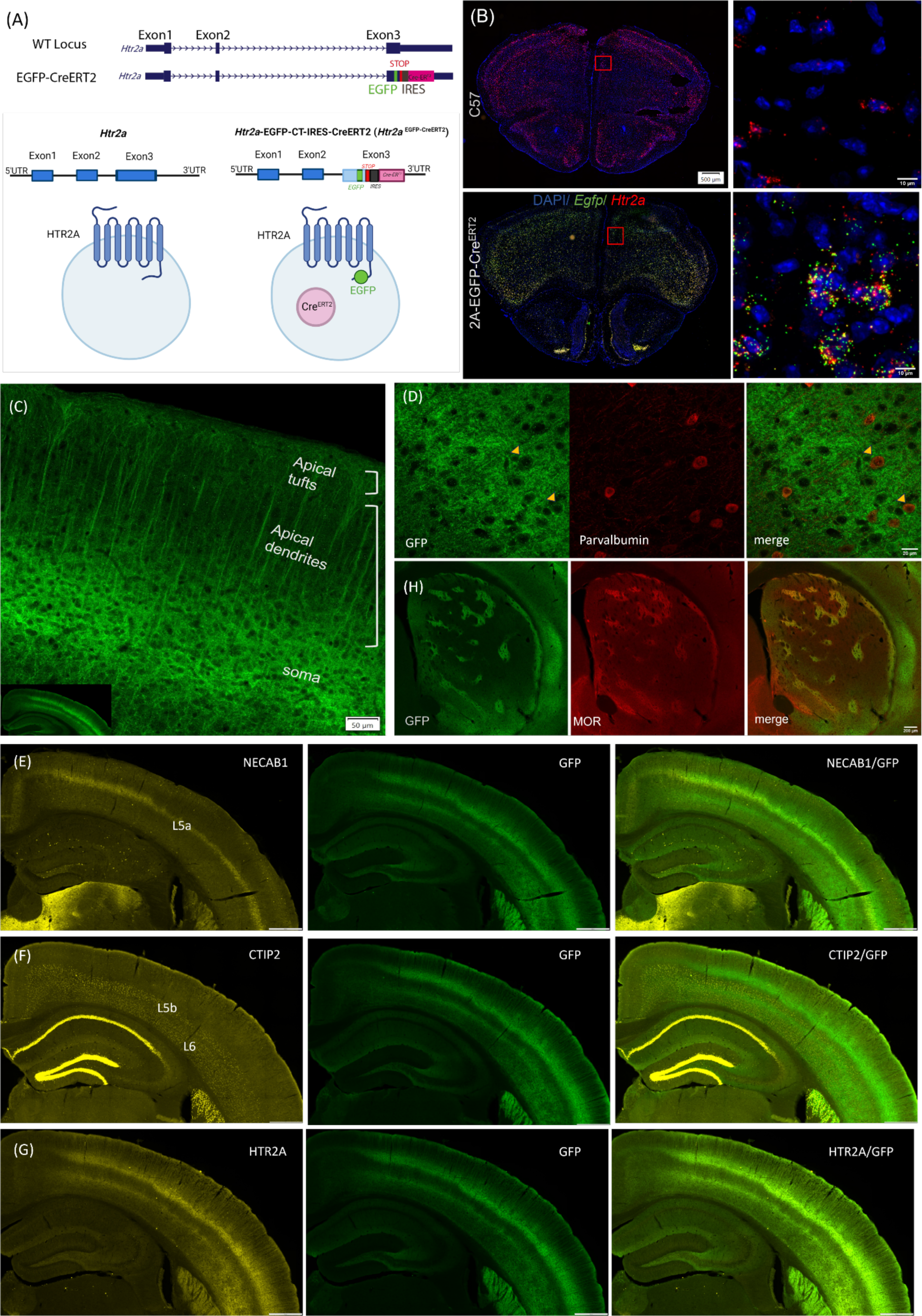
The genetic design of *Htr2a*-EGFP-CT-IRES-CreERT2 knockin mice and validation. (A) Schema of the transgenic modifications at the murine *Htr2a* gene in exon 3. For *Htr2a*-EGFP-CT-IRES-CreERT2 (*Htr2a*^EGFP-CreERT2^) mouse line, EGFP was inserted into the C terminus (CT) of the receptor, after residue 452. An IRES-CreERT2 cassette was inserted between the *Htr2a* stop codon and the 3’UTR. Blue boxes represent exons of murine *Htr2a*; the green box is EGFP; the red box shows the STOP codon. The gray box is the IRES followed by a pink box for CreERT2. The schematic was created with https://www.biorender.com/. (B) Validation of *Egfp* and *Htr2a* mRNA expressions in the mouse brain: RNAscope experiments were performed to probe *Egfp* and *Htr2a* mRNA in C57 (*Htr2a*^+/+^) and *Htr2a* ^EGFP-CreERT2/EGFP-CreERT2^ mice. The *Htr2a* (red) probe can be detected in C57 mice; *Egfp* (green) and *Htr2a* (red) probes showed co-localization (yellow) in *Htr2a*^EGFP-CreERT2/EGFP-CreERT2^ mice. Images were taken under an Olympus slide scanner with 10X objective and an Olympus confocal microscope with a 60X objective. (C) The expression pattern of HTR2A in cortical pyramidal neurons (soma, apical dendrites, and apical tufts). Images were taken with a 20X objective under an Olympus confocal microscope. (D) Paravalbumin is the GABAergic interneuron marker. The brain sections from *Htr2a* ^EGFP-CreERT2/+^ mice were stained with anti-parvalbumin and anti-GFP antibodies. The yellow arrowhead denotes parvalbumin-positive and GFP-positive interneurons. Images were captured under a 60X objective with an Olympus confocal microscope. (E) Cortical L5a marker NECAB1 was used to locate the cortical layer 5a distribution. HTR2A-EGFP-CT fusion receptors with anti-GFP staining showed its distribution in the NECAB1-positive layer. (F) With CTIP2 as a cortical L5b/L6 marker, HTR2A-EGFP-CT did not show overlap with the Ctip2-positive layers. (G) An anti-HTR2A antibody showed the same pattern as the anti-GFP distribution associated with HTR2A-EGFP-CT. Note, all brain sections in Figures 1E-1H from *Htr2a*^EGFP-CreERT2/+^ mice were stained with an anti-GFP antibody for HTR2A-EGFP-CT and different cortical markers (NECAB1, CTIP2), as well as with an anti-HTR2A antibody. (H) The distribution pattern of HTR2A-EGFP-CT labeling in the dorsal striatum. The Mu opioid receptor (MOR) is rich in the striosome and subcallosal stria of the Caudate-Putamen (CPu). HTR2A-EGFP-CT signals showed the same expression pattern with MOR distribution in the patch-like (striosome) area and stria. Experiments in this figure were conducted with 3-4 mice with similar results. Images were taken using an Olympus VS120 slide scanner or an Olympus confocal microscope under 60X objective.

To characterize these mice, we first examined whether EGFP cDNA inserted into exon3 of *Htr2a* genetic locus was expressed in a manner similar to the endogenous receptor. Here, *in situ* hybridization was used to detect the mouse *Htr2a* and *Egfp* transcripts in C57BL/6J (C57, *Htr2a*^+/+^) and *Htr2a*-EGFP-CT-IRES-CreERT2 mice. As shown in Figure 1B, *Egfp* mRNA was expressed in the same cells as *Htr2A mRNA* in knock-in mice, but not in C57 animals. Next, we examined expression patterns/levels of HTR2A-EGFP-CT with GFP antibodies and immunofluorescent microscopy in fixed brain slices (Fig 2; Table 1) and by light-sheet microscopy in cleared brains (Movie S1-S4). Both strategies revealed similar results in corresponding coronal sections. A comparison of the expression patterns between HTR2A-EGFP-CT (Suppl. Fig S1E) and the previously determined pattern of ^3^H-M100907**-**labelled HTR2A (Lopez-Gimenez et al., 1997) (Suppl. Fig S1D) revealed excellent correspondence with expression of the receptor in our *Htr2a*^EGFP-CreERT2^ mice. A summary analysis of this receptor expression pattern in different brain regions follows in the next section, whereas a detailed description is found in **Supplement #1**.

**Fig 2.**
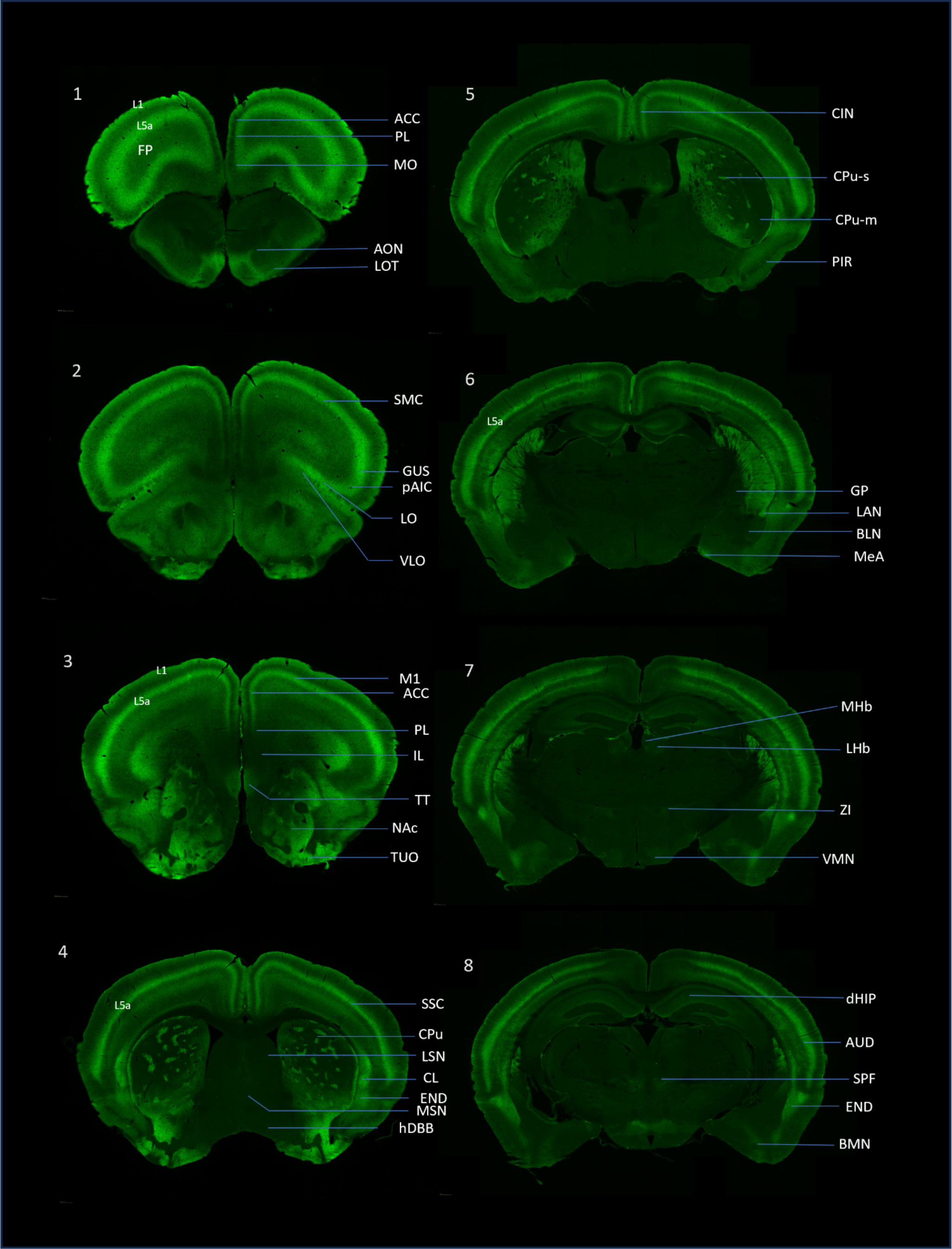
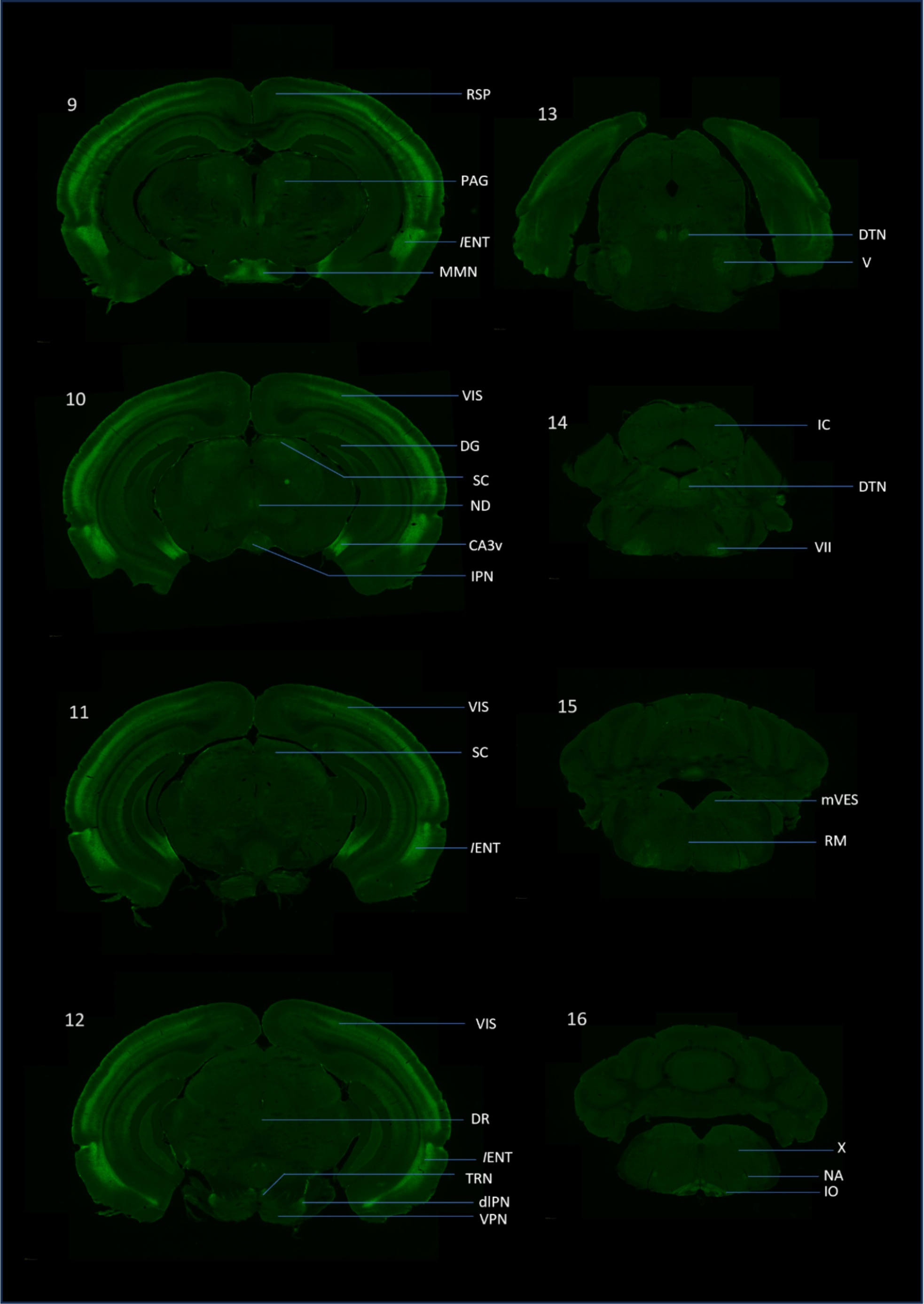
The distribution of the HTR2A-EGFP-CT fusion protein in mouse brain. Brain sections from *Htr2a*^EGFP-CreERT2/+^ mice were stained with a GFP antibody to amplify the HTR2A-EGFP-CT signal and then were acquisitioned with an Olympus slide scanner under 10X objective. The abbreviations for the brain areas are located in the Supplementary Figure. The raw images were uploaded to open-source website, A Mouse Imaging Server (AMIS, https://amis2.docking.org/.

**Table 1.**
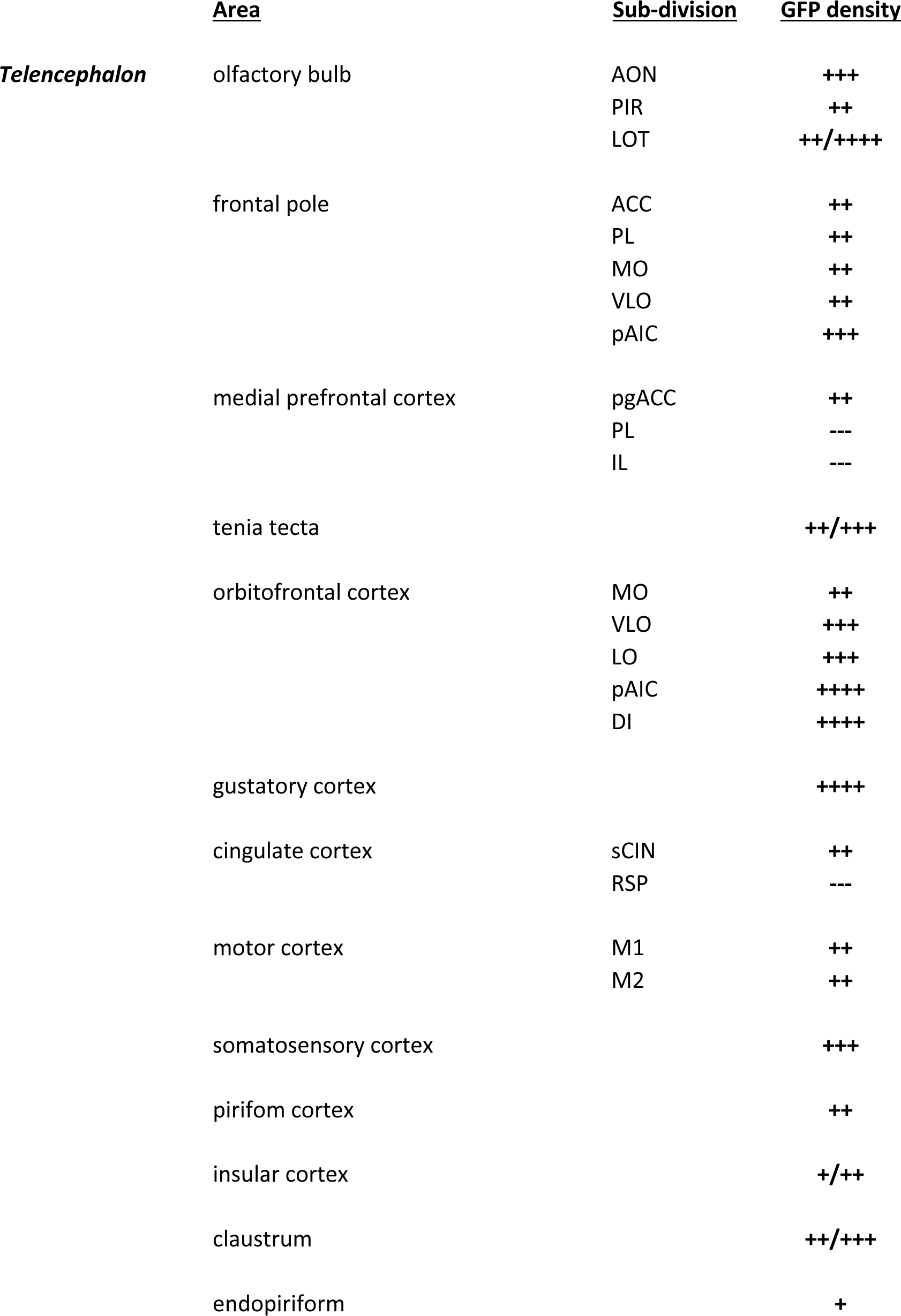
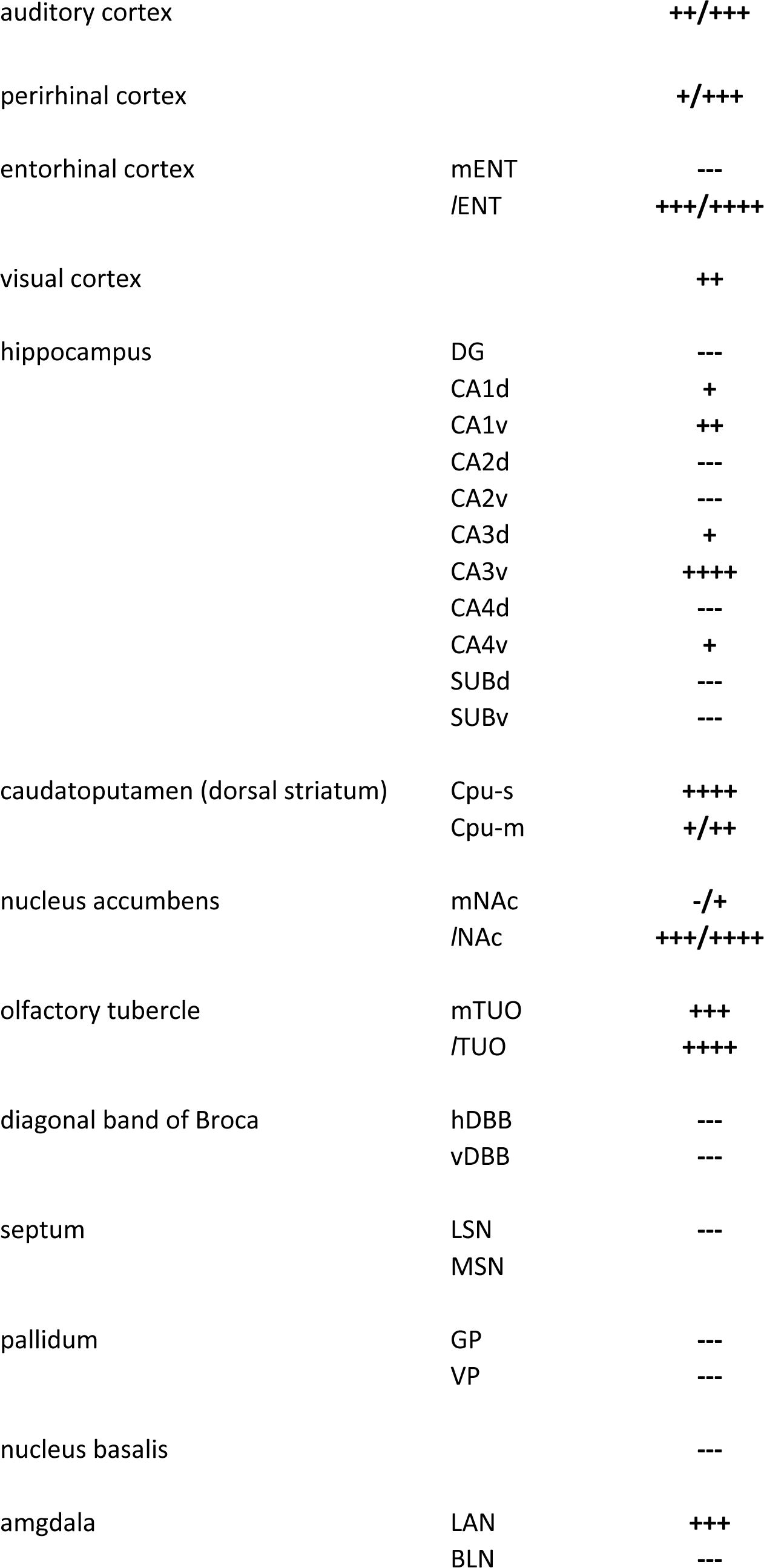
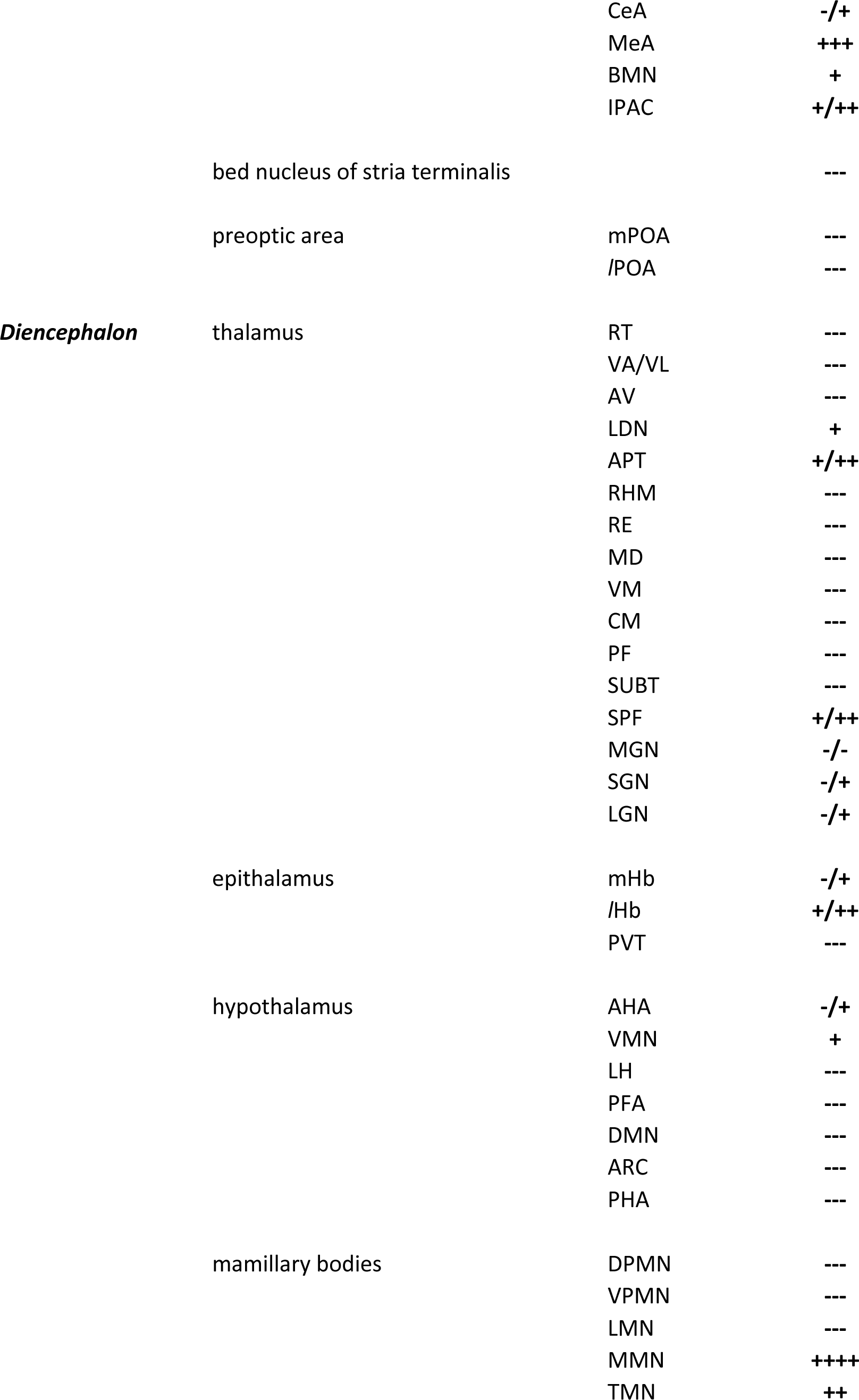
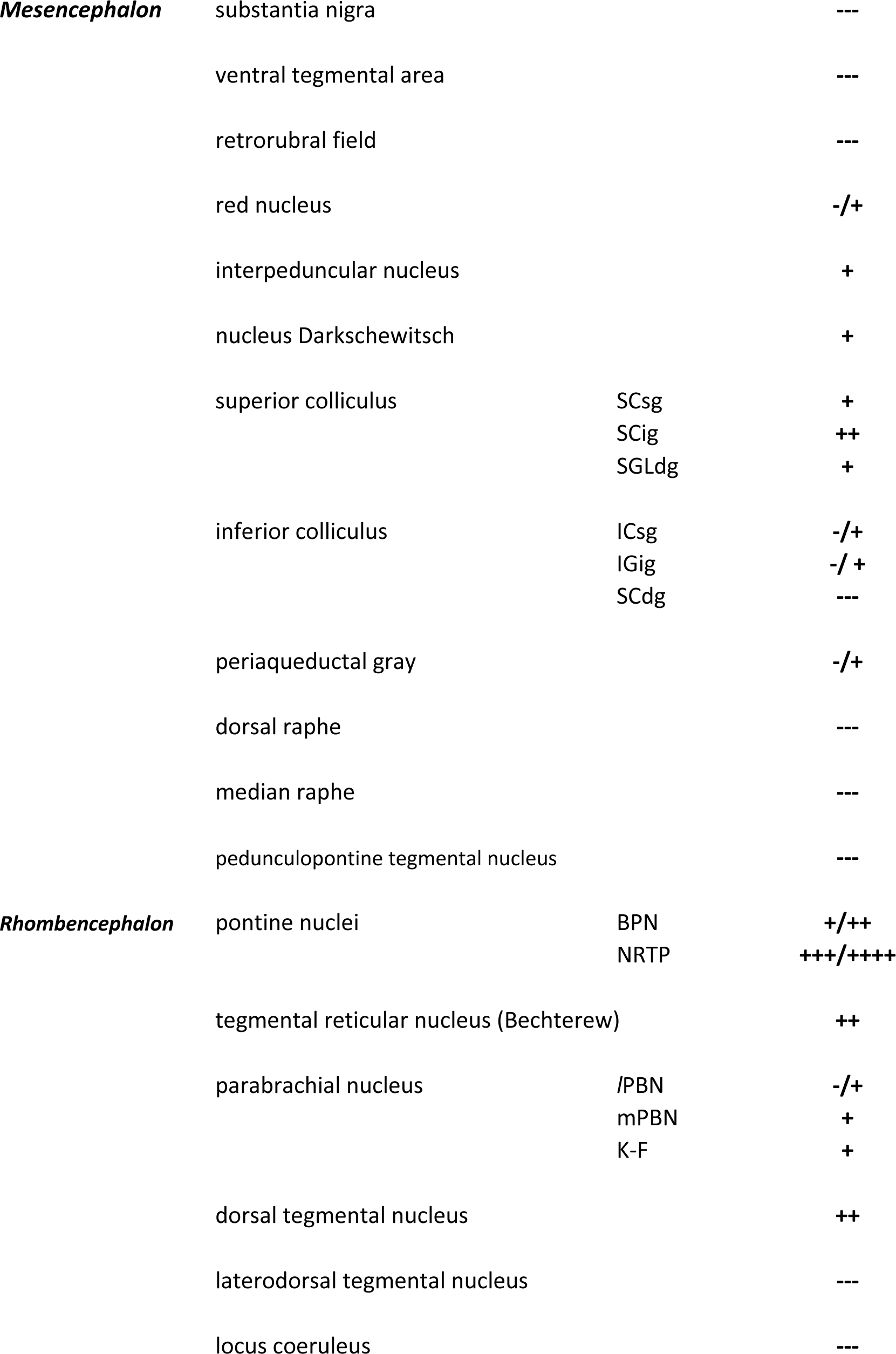
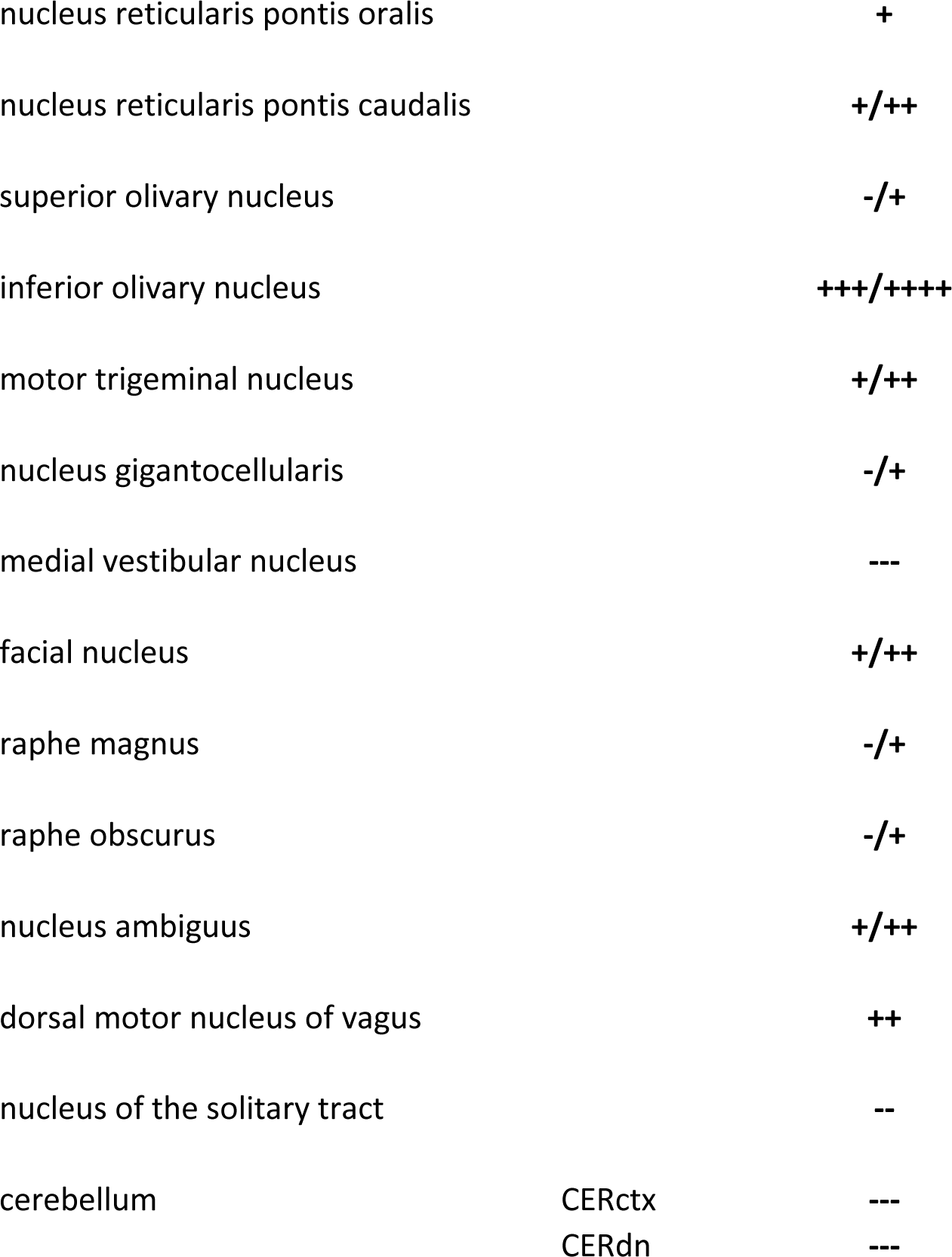
Whole brain distribution of HTR2A-EGFP-CT in *Htr2a*^EGFP-CreERT2^ mice.

### Distribution of HTR2A-EGFP-CT receptor in mouse brain

The EGFP fluorophore property was used to locate HTR2A-EGFP-CT fusion proteins for the whole brain mapping. The HTR2A-EGFP-CT receptor was densely expressed in several forebrain areas, while in more caudal brain regions, receptor expression was restricted to only a few areas, most of which had relatively low expression. In neocortical areas, there was a dense band of HTR2A-EGFP-CT receptors in the most superficial aspect of layer 5a (L5a). Scattered HTR2A-EGFP-CT receptors were also present in L6 and supra-granular lamina with the exception of layer 1, where the apical tufts of HTR2A-positive pyramidal cells could be clearly visualized.

In the frontal pole (FP), HTR2A-EGFP-CT in L5a formed a distinct continuous band that encircled the entire hemisphere, running parallel to the pial surface (Fig 2.1). In the medial prefrontal area caudal to the frontal pole, where the infralimbic cortex (IL) first appears (Fig 2.2), the continuous nature of the band of HTR2A-EGFP-CT-labeled L5a cells was disrupted by a zone in the medial PFC in which the HTR2A cells were absent in the ventral prelimbic (PL) and infralimbic cortices (IL) (Fig 2.3). Caudal to the level of the genu of the callosum, HTR2A-positive L5a cells extended from the cingulate cortex medially to the piriform cortex (PIR) laterally (Fig 2.4 to Fig 2.6). At the level of the mesodiencephalic juncture (Fig 2.6 to Fig 2.12), where the PIR was no longer present, low HTR2A-EGFP-CT signal was present in the retrosplenial cortex (RSP) (Fig 2.6), while the most laterally situated L5a labeled cells were in the lateral entorhinal cortex (*l*ENT) (Fig 2.9 to Fig 2.12). Still more posteriorly, HTR2A-EGFP-CT was seen in the visual cortices (VIS) (Fig 2.10 to Fig 2.12).

The densely-labeled L5a HTR2A-EGFP-CT cortical band was primarily comprised of pyramidal neurons with characteristic apical dendrites extending across L2/3 to enter and terminate in L1 as apical tufts (Figure 1C). The dense HTR2A-EGFP-CT immunoreactivity that filled the somatodendritic compartments initially suggested cells in this cortical band were located in both superficial L5a and L4 (Pazos et al., 1985; Lopez-Gimenez et al., 1997). However, the cell bodies of these HTR2A-EGFP-CT signals were predominantly found in L5a, with a characteristic pyramidal morphology rather than the small (stellate) cells of L4. Moreover, the cortical band of the HTR2A-EGFP-CT signal was present in the prelimbic cortex, which lacks a defined L4. We also stained sections for parvalbumin, a marker of the largest subset of cortical interneurons (Tamamaki et al., 2003). Only a minority of neurons in the HTR2A-EGFP-CT cortical band expressed parvalbumin immunoreactivity (Figure 1D), consistent with both pyramidal neurons and occasional interneurons expressing HTR2A (Willins et al., 1997a).

Early studies of the localization of HTR2A using autoradiographic methods concluded that the pyramidal cells that form a strongly-labeled band of cells across the cortex were in layer 4 (Pazos et al., 1985) and layer 5 (Lopez-Gimenez et al., 1997) However, these studies suffered from a lack of resolution. We therefore defined the laminar position of HTR2A-expresssing pyramidal cells by comparing HTR2A-EGFP-CT and well-characterized cortical laminar markers. These markers included N-terminal EF-hand calcium binding protein 1 (NECAB1), which is primarily localized to L5a, and the L5b/L6 marker COUP-TF-interacting protein 2 (CTIP2). NECAB1 staining overlaped with the major band of cells expressing HTR2A-EGFP-CT (Fig 1E). In contrast, the laminar distributions of CTIP2- and HTR2A-EGFP-CT expression did not overlap (Fig 1F). A validated HTR2A antibody (Magalhaes et al., 2010a; Yadav et al., 2011b) was also used to determine if the expression pattern of the HTR2A protein corresponded with the GFP-stained HTR2A-EGFP-CT fusion protein; both antibodies displayed similar staining patterns (Fig 1G). Therefore, the prominent layer expressing dense HTR2A-EGFP-CT-positive pyramidal cells is L5a and not L4.

HTR2A-EGFP-CT in the telencephalon was not restricted to the cortex but was also present in subcortical sites. In the dorsal striatum (caudatoputamen, CPu), HTR2A-EGFP-CT densely filled the striosomal compartments (CPu-s) (defined on the basis of mu opioid receptor (MOR) immunoreactivity). HTR2A-EGFP-CT was co-localized with MOR in striosomes, including the subcallosal stria (Fig1H). HTR2A-EGFP-CT was present but much less dense in the matrix (CPu-m) compartment (Fig 2.4 to Fig 2.5). Ventral striatal areas, including the nucleus accumbens (NAc) and olfactory tubercle (TUO) were among the most densely labeled nuclei in the brain (Fig 2.3). In the ventral striatum, there was a distinct lateral-to-medial gradient of HTR2A-EGFP-CT labeling, with the most lateral NAc areas (the ventrolateral shell and core) exhibiting a greater density of HTR2A-EGFP-CT than more medial accumbens (medial shell and septal pole) territories (Fig 2.3). A similar gradient was observed across the mediolateral extent of the TUO. Pallidal projection targets of the dorsal and ventral striatum lacked HTR2A-EGFP-CT expression, as did other forebrain areas in the medial telencephalon, including the septal (LSN and MSN, lateral and medial septal nucleus) and diagonal band complexes (hDBB, diagonal band, horizontal limb) (Fig 2.4).

The entire rostrocaudal extent of the claustrum (CL) expressed HTR2A-EGFP-CT, with labeling of the adjacent endopiriform cortex (END) being slightly less dense (Fig 2.4). Still weaker was labeling in the laterally contiguous insular cortex (IC) (Fig 2.2 to Fig 2.4). In the amygdala (Fig 2.6), the lateral nucleus (LAN) expressed high levels of HTR2A-EGFP-CT, whereas the ventromedially adjacent basolateral nucleus (BLN) was devoid of signal. The medial nucleus (MeA) was moderately filled with HTR2A-EGFP-CT, while the remainder of the amygdala expressed trace (e.g., the basomedial nuclei) or no HTR2A-EGFP-CT (e.g., the central nuclei). The hippocampus was notable for a very dense aggregation of HTR2A-EGFP-CT in the ventral-most CA3 field of the ventral (temporal) (Fig 2.9 to Fig 2.11), but not dorsal (septal) hippocampus (Fig 2.6 to Fig 2.8).

HTR2A-EGFP-CT was present in very few diencephalic areas. In the thalamus there was low expression of HTR2A-EGFP-CT in the anterior pretectal area and subparafascicular nucleus (SPF) (Fig 2.8), while in the hypothalamus low-to-moderate density filling of the zona incerta (ZI) and ventromedial nucleus (VMN) were noted (Fig 2.7). The only densely-labeled site in the diencephalon was the medial mammillary nucleus (MMN) (Fig 2.9). No strongly labeled areas were seen in the midbrain. Trace-to-light density HTR2A-EGFP-CT labeling was present in the superior colliculus (SC), especially in the intermediate gray (Fig 2.11), with a somewhat lower density of labeling in the inferior colliculus (IC) (Fig 2.14). Low levels of HTR2A-EGFP-CT were seen in the interpeduncular nucleus (IPN) (Fig 2.10).

In the pons, HTR2A-EGFP-CT expression was restricted to a few pontine nuclei. The dorsolateral and dorsomedial aspects of the pontine nuclei and the nucleus reticularis tegmenti pontis were densely labeled. The dorsal tegmental nucleus (DTN) (Fig 2.13) was moderately labeled, as was the motor trigeminal (V) (Fig 2.13), facial nuclei (VII) (Fig 2.14), and the nucleus ambiguous (NA) (Fig 2.16). Low levels of HTR2A-EGFP-CT were seen in the parabrachial and Kolliker-Fuse nuclei. With the exception of the raphe obscurus and raphe magnus (RM) (Fig 2.15), which contained very sparse labeling, the raphe nuclei did not express HTR2A-EGFP-CT. In the medulla, trace-to-low levels of expression were present in the dorsal motor nucleus of the vagus (X) and the nucleus of the solitary tract (Fig 2.16). The only densely-labeled medullary site was the inferior olive (IO) (Fig 2.16).

### Behavioral analyses reveal prototypical responses to psychedelics in *Htr2a-*EGFP-CT-IRES-CreERT2 mice

In mice and rats, acute administration of psychedelic drugs produces a stereotypical head twitch response (Glennon et al., 1984; Gonzalez-Maeso et al., 2007; Keiser et al., 2009b), disruption of prepulse inhibition (PPI) (Sipes et al., 1995), and variable changes in motor behaviors (Rodriguiz et al., 2021). Accordingly, we next evaluated these responses to vehicle, 0.3 mg/kg LSD, 1 mg/kg 2,5-dimethoxy-4-iodoamphetamine (DOI), and 1 mg/kg psilocin (i.p.) in C57 and *Htr2a*^EGFP-CreERT2^ mice to examine whether HTR2A-EGFP-CT function was similar to endogenous HTR2A.

#### Effects of psychedelics on motor activities

In the open field baseline activities were collected in 5-min blocks over 30 min and post-injection responses were followed over the next 90 min. Locomotor activities were increased overall in both genotypes by DOI relative to vehicle (*p*-values≤0.031) (Suppl. Fig S2A-C). In contrast, only LSD and DOI augmented overall rearing (*p*-values≤0.001) (Suppl. Fig S2D-E), while only DOI increased stereotypy (*p*=0.035) (Suppl. Fig S2G-H) in C57 controls and *Htr2a*^EGFP-CreERT2^ animals. Both genotype and treatment effects can be visualized more readily when the results are presented as cumulative responses. In all cases, baseline levels of locomotor, rearing, and stereotypical activities were higher overall in *Htr2a*^EGFP-CreERT2^ than in the C57 mice (*p*-values≤0.004) (Fig 3A-C, *left*). To control for these baseline differences, the post-injection responses were normalized using analysis of covariance (ANCOVA).

**Fig 3.**
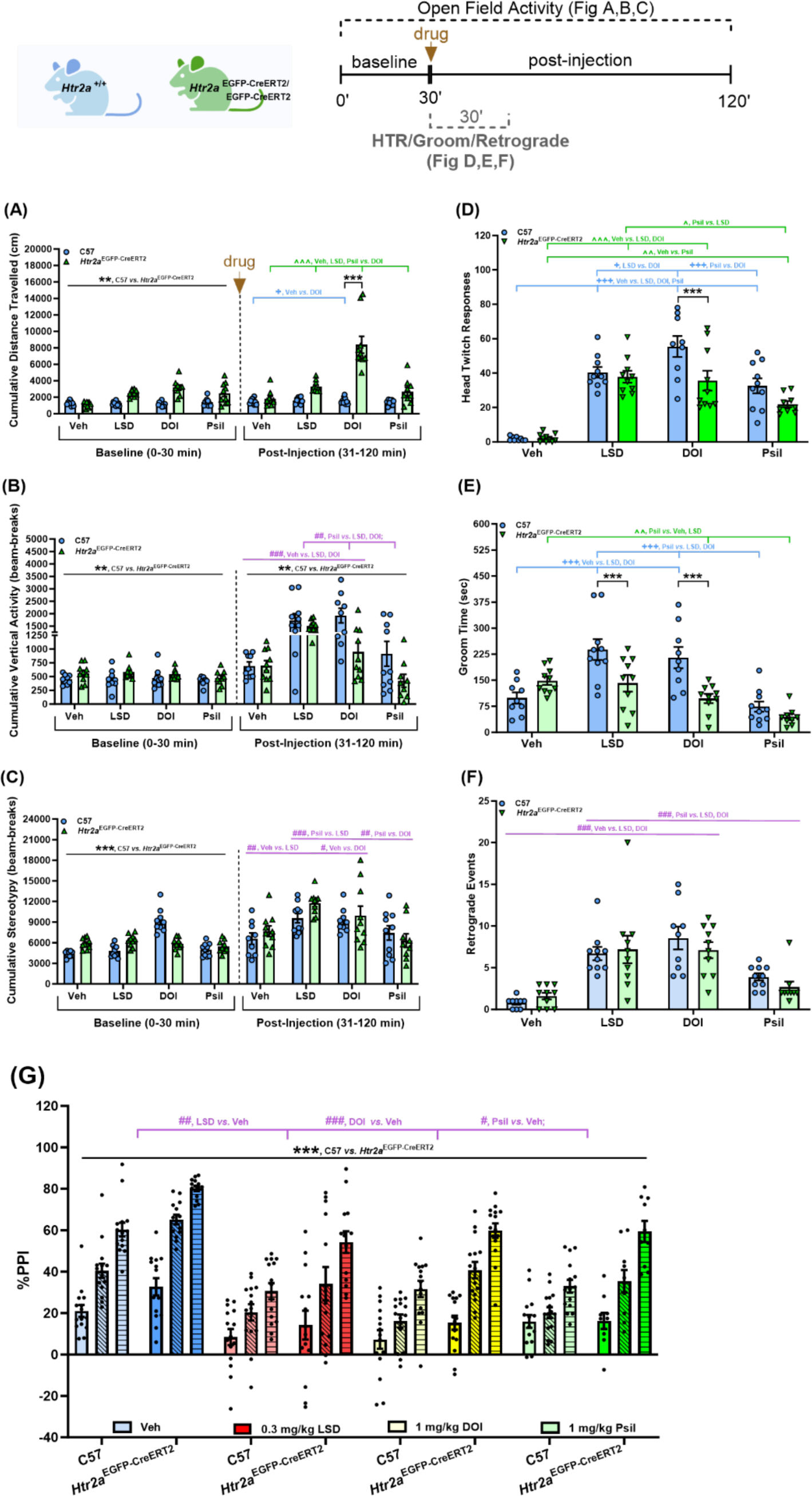
Pharmacological effects of psychedelics (LSD, DOI, and Psilocin) on behaviors in C57BL/6J and *Htr2a*^EGFP-CreERT2/EGFP-CreERT2^ mice. **(A-C) *Cumulative motor activities*** Cumulative baseline activities (0-30 min; pre-administration) and cumulative post-injection activities following administration (31-120 min) of the vehicle (Veh), LSD (0.3 mg/kg), DOI (1 mg/kg) or psilocin (Psil, 1 mg/kg); n=9-10 mice/genotype/treatment. Note, the C57BL/6J and *Htr2a*^EGFP-CreERT2/EGFP-CreERT2^ mice are termed C57 and *Htr2a*^EGFP-CreERT2^ mice, respectively, below and in the figure panels. The symbol “*” is for genotype difference (black); “+” is for within C57 effect (blue); “^” for within *Htr2a*^EGFP-CreET2^ effect (green), and “#” for overall treatment effects (purple). All data are represented as means ± SEMs. (A) Locomotor activities in C57 and *Htr2a*^EGFP-CreERT2^ mice. Two-way ANOVA for baseline: genotype [F(1,69)=8.968, *p*=0.004]; for C57 *vs*. *Htr2a*^EGFP-CreERT2^ mice (black): \*\**p*<0.01, overall genotype effects. Two-way ANCOVA for post-injection: genotype [F(1,68)=19.433, *p*<0.001], treatment [F(3,68)=29.894, *p*<0.001], and genotype by treatment interaction [F(3,68)=11.143, *p*<0.001]; Bonferroni *post-hoc* tests; for C57 *vs*. *Htr2a*^EGFP-CreERT2^ mice (black): \*\*\**p*<0.001, DOI; for C57 mice (blue): ^+^*p*<0.05, Veh *vs.* DOI; for *Htr2a*^EGFP-CreERT2^ mice (green): ^^^^^p<0.001, Veh, LSD, Psil *vs.* DOI. (B) Rearing activities in the same mice. Two-way ANOVA for baseline: genotype [F(1,69)=9.307, *p*=0.003]; for C57 *vs*. *Htr2a*^EGFP-CreERT2^ mice (black): \*\**p*<0.01, overall genotype effects. Two-way ANCOVA for post-injection: genotype [F(1,68)=17.900, *p*<0.001] and treatment [F(3,68)=13.765, *p*<0.001]; Bonferroni *post-hoc* tests; for C57 *vs*. *Htr2a*^EGFP-CreERT2^ mice (black): \*\**p*<0.01, overall genotype effects; overall treatment effects (purple): ^##^*p*<0.01, Psil *vs.* LSD and DOI; ^###^*p*<0.001, Veh *vs.* LSD and DOI. (C) Stereotypical activities in these same mice. Two-way ANOVA for baseline: genotype [F(1,69)=44.086, *p*<0.001]; for C57 *vs*. *Htr2a*^EGFP-CreERT2^ mice (black): \*\*\**p*<0.001, overall genotype effects. Two-way ANCOVA for post-injection: treatment [F(3,68)=8.338, *p*<0.001]; Bonferroni *post-hoc* tests; overall treatment effects (purple): ^#^p<0.05, Veh *vs.* DOI; ^##^p<0.01, Veh *vs.* LSD or Psil *vs.* DOI; ^###^p<0.001, Psil *vs.* LSD. **(D-F) *Head twitch responses, grooming, and retrograde walking*** These responses were scored beginning immediately after vehicle or psychedelic administration and followed over the next 30 min; n=9-10 mice/genotype/treatment. (D) Head twitch responses in C57 and *Htr2a*^EGFP-CreERT2^ animals following administration (31-60 min) of the Veh, 0.3 mg/kg LSD, 1 mg/kg DOI, or 1 mg/kg Psil. Two-way ANOVA: genotype [F(1,69)=9.236, *p*=0.003], treatment [F(3,69)=51.113, *p*<0.001], and genotype by treatment interaction [F(3,69)=2.878, *p*=0.042]; Bonferroni *post-hoc* tests: for C57 *vs. Htr2a*^EGFP-CreERT2^ mice (black): \*\*\**p*<0.001, DOI; for C57 mice (blue): ^+^*p*<0.05, LSD *vs.* DOI; ^+++^*p*<0.001, Veh *vs.* all psychedelics, or Psil *vs.* DOI; for *Htr2a*^EGFP-CreERT2^ mice (green): ^*p*<0.05, Psil *vs.* LSD; ^^*p*<0.01, Veh *vs.* Psil; ^^^*p*<0.001, Veh *vs.* LSD and DOI. (E) Groom time in the same mice. Two-way ANOVA: genotype [F(1,69)=11.465, *p*<0.001], treatment [F(3,69)=15.125, *p*<0.001], and genotype by treatment interaction [F(3,69)=6.855, *p*<0.001]; Bonferroni *post-hoc* tests: for C57 vs *Htr2a*^EGFP-CreERT2^ mice (black): \*\*\**p*<0.001, LSD and DOI; for C57 mice (blue): ^+++^*p*<0.001; Veh *vs.* LSD and DOI or or Psil *vs.* LSD and DOI; for *Htr2a*^EGFP-CreERT2^ mice (green): ^^*p*<0.01, Psil *vs.* Veh and LSD. (F) Retrograde walking in these same mice. Two-way ANOVA for treatment [F(3,69)=21.713, *p*<0.001]; Bonferroni *post-hoc* tests: overall treatment effects (purple): ^###^*p*<0.001, Veh *vs.* LSD and DOI, or Psil *vs.* LSD and DOI. **(G) Prepulse inhibition** C57 and *Htr2a*^EGFP-CreERT2^ mice were injected with the Veh, 0.3 mg/kg LSD, 1 mg/kg DOI, or 1 mg/kg Psil and tested 10 min later; n=10-15 mice/genotype/treatment. RMANOVA: PPI [F(2,210)=329.097, *p*<0.001], PPI by genotype interaction [F(2,210)=23.856, *p*<0.001], PPI by treatment interaction [F(6,210)=3.342, *p*=0.004], genotype [F(1,105)=45.808, *p*<0.001], and treatment [F(3,105)=19.562, *p*<0.001]; Bonferroni *post-hoc* tests: for C57 vs *Htr2a*^EGFP-CreERT2^ mice (black): \*\*\**p*<0.001, overall genotype effects; for treatment effects (purple): ^#^*p*<0.05, Veh *vs.* Psil; ^##^*p*<0.01, Veh *vs.* LSD; ^###^*p*<0.001, Veh *vs.* DOI. Some schematics were created with https://www.biorender.com/.

DOI enhanced locomotion in the open field to a greater extent overall in *Htr2a*^EGFP-CreERT2^ than in C57 animals (*p*<0.001) (Fig 3A). Within C57 mice, DOI augmented locomotion over that of the vehicle control (*p*=0.020), whereby in *Htr2a*^EGFP-CreERT2^ animals DOI was more potent in stimulating activity than all other treatments (*p*-values<0.001).

Rearing responses to DOI were differentiated by main effects of genotype and treatment (Fig 3B). Rearing was lower overall in *Htr2a*^EGFP-CreERT2^ than C57 mice (*p*<0.001). With regards to treatment, responses to the vehicle and psilocin were similar, but both were lower than the corresponding LSD- and DOI-elicited responses (*p*-values≤0.002). When stereotypy was analyzed, treatment effects emerged (Fig 3C). Overall responses to the vehicle and psilocin were not distinguished from each other, while both LSD and DOI augmented stereotypical behaviors over these two groups (*p*-values0.026).

Together, these results show that basal motor activities were higher in the *Htr2a*^EGFP-CreERT2^ than in C57 mice and that DOI was more potent in stimulating locomotion in the knock-in animals. Following vehicle or psychedelic administration, these genotype effects persisted in locomotion and rearing with ANCOVA, whereas DOI in locomotion and LSD and DOI in rearing and stereotypes were more potent in stimulating activities compared to psilocin and the vehicle control.

#### Psychedelic-induced head twitch and ethological responses

LSD, DOI, and psilocin stimulated more head twitch responses (HTRs) in the C57 and *Htr2a*^EGFP-CreERT2^ mice relative to the vehicle controls *p*-values<0.003), where responses were very low (Fig 3D). DOI was more potent in stimulating head twitch responses in C57 than in *Htr2a*^EGFP-CreERT2^ mice (*p*<0.001), with a strong trend for psilocin (*p*=0.053). No genotype differences were evident between the vehicle and LSD treatments. Within C57 mice, all psychedelics stimulated more head twitch responses than vehicle (*p*-values<0.001). Additionally, DOI stimulated more head twitch responses than LSD or psilocin (*p*-values≤0.043), which were not statistically different from each other. In *Htr2a*^EGFP-CreERT2^ animals, the numbers of head twitch responses were lower with vehicle than all hallucinogen-treated groups (*p*-values≤0.003), and LSD was more efficacious than psilocin (*p*=0.026). Thus, both genotype and treatment effects in head twitch responses were observed.

Time spent grooming was also examined (Fig 3E). Grooming was more prolonged in the C57 mice given LSD and DOI relative to that in the *Htr2a*^EGFP-CreERT2^ animals (*p*-values<0.001). Within C57 subjects, both LSD and DOI stimulated grooming for longer periods of time than those observed in the psilocin- and vehicle-treated mice (*p*- values<0.001), which were not different from each other. By comparison, in *Htr2a*^EGFP-CreERT2^ animals groom time was reduced with psilocin relative to the LSD and vehicle groups (*p*-values≤0.008). In summary, grooming is largely reduced by psilocin in both genotypes relative to vehicle, whereas genotype differences are evident with LSD and DOI. No genotype distinctions were found for retrograde walking (Fig 3F). Here both LSD and DOI stimulated this response to greater extents than in the psilocin- and vehicle-treated mice (*p*-values<0.001).

#### Differential effects of psychedelics on prepulse inhibition (PPI)

Genotypic differences were evident where PPI was lower overall in C57 than in *Htr2a*^EGFP-CreERT2^ mice (*p*<0.001) (Fig 3G). Moreover, within genotypes all three hallucinogens disrupted PPI relative to vehicle (*p*-values<0.001). Both null and startle activities were analyzed. Overall null activity was higher in *Htr2a*^EGFP-CreERT2^ than in C57 animals (*p*-values<0.001) -(Suppl. Fig S3A). An assessment of treatment effects revealed overall null activities were lower in the vehicle than in the LSD and DOI groups (*p*-values<0.001). In addition, null activity was higher in DOI- than in LSD- and psilocin-treated mice (*p*- values<0.001). Genotype effects were observed also for startle activities (Suppl. Fig S3B). Here responses to the vehicle, LSD, and psilocin were reduced in *Htr2a*^EGFP-CreERT2^, compared to the C57 animals (*p*-values<0.001). Within C57 mice startle activities were higher overall in the psilocin than in the vehicle or LSD groups (*p* -values<0.044). In *Htr2a*^EGFP-CreERT2^ mice, startle to DOI was higher than in the vehicle and LSD animals (*p*- values<0.010). Collectively, these findings indicate that PPI was disrupted in both genotypes with each of the psychedelics; however, PPI was more perturbed in C57 than in the *Htr2a*^EGFP-CreERT2^ mice.

### Biochemical validation of *Htr2a* ^EGFP-CreERT2^ mice

[^3^H]-ketanserin saturation binding assay was used to quantify the affinity (K_d_) and number of HTR2A-EGFP-CT receptors (B_max_) in the whole cerebral cortex (Fig 4A). C57 and *Htr2a* ^EGFP-CreERT2^ mice showed no statistically significant differences in K_d_ values (p=0.35) although receptor expression in the *Htr2a*^EGFP-CreERT2^ mice was statistically enhanced compared to C57 controls (p=0.0005). Receptor mRNA levels *via* real-time qPCR were examined from the whole brain cortex in C57, as well as in heterozygous and homozygous *Htr2a*^EGFP-CreERT2^ mice (Fig 4B). *Htr2A* mRNA expression levels in *Htr2a*^EGFP-CreERT2^ mice were statistically enhanced in the heterozygous (*p*=0.0002) and in the homozygous *Htr2a*^EGFP-CreERT2^ mice (*p*<0.0001) relative to C57 controls. Increased transcript levels have been previously reported for other 3’ manipulations of GPCRs (Erbs et al., 2015) (Scherrer et al., 2006) (Chen et al., 2020).

**Fig 4.**
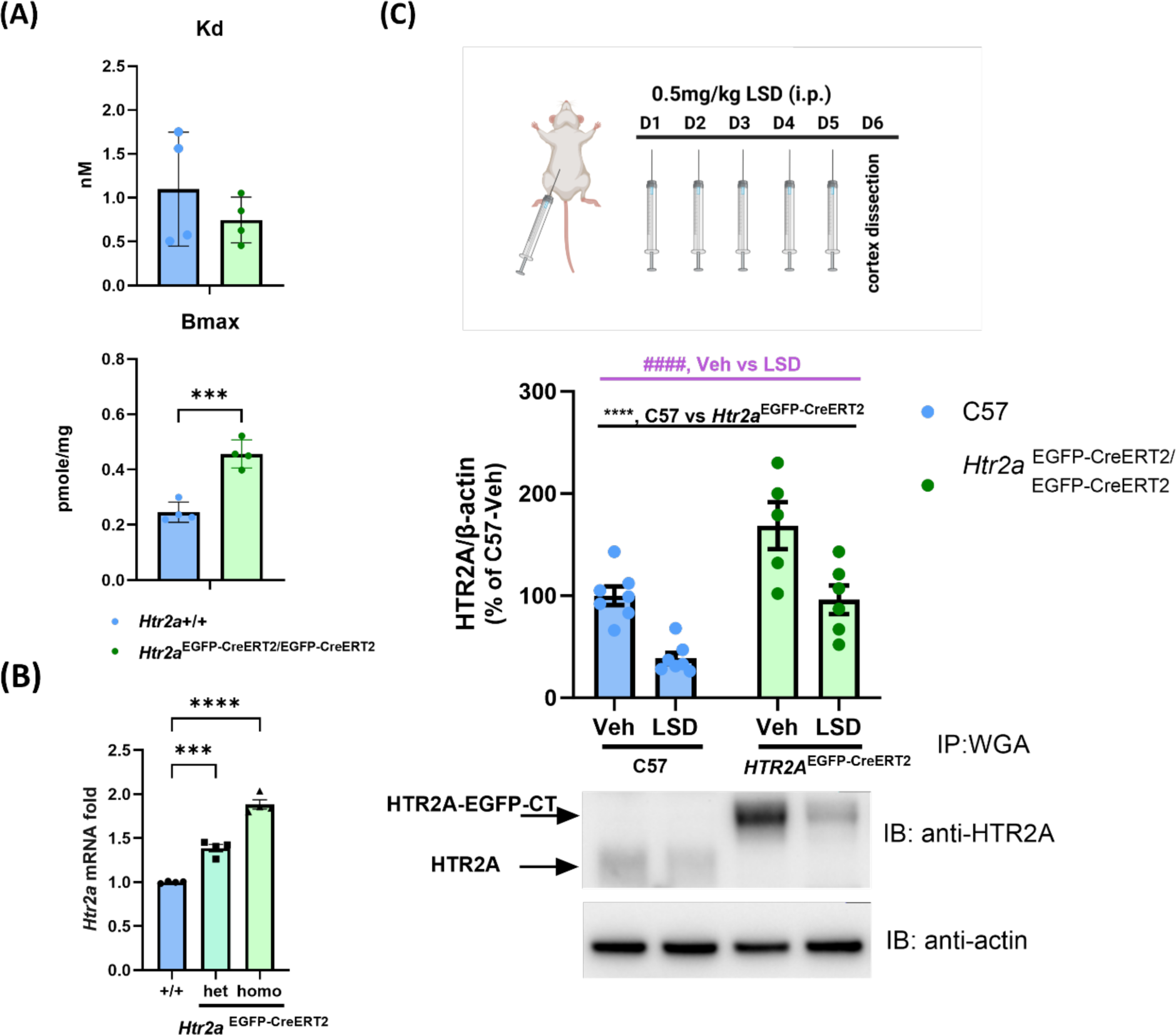
Receptor functionality validations of HTR2A in transgenic *Htr2a*^EGFP-CreERT2^ mice. (A) [^3^H] ketanserin saturation binding assay was conducted to examine the affinity (Kd, nM) and receptor levels (Bmax, pmole/mg) of HTR2A in the whole cortex of C57 and *Htr2a*^EGFP-CreERT2^ mice. Non-specific binding was determined with 10 µM clozapine. Data are means ± SEMs (n=4 samples/genotype) and analyzed by unpaired t-test. Kd values: C57(1.09 ± 0.324 nM) *vs. Htr2a*^EGFP-CreERT2^ (0.75 ± 0.13nM), [t(6)=1.005, *p*=0.354]; Bmax values: C57 (0.25 ±0.018 pmole/mg) *vs. Htr2a*^EGFP-CreERT2^ (0.46 ± 0.025 pmole/mg), [t(6)=6.745, *p*=0.0005] (\*\*\**p*<0.001, C57 *vs. Htr2a*^EGFP-CreERT2^). (B) Real-time qPCR was used to determine *Htr2a* mRNA levels from the whole cortex of C57 mice ^(*Htr2a*+/+)^, *Htr2a*^EGFP-CreERT2/+^, and *Htr2a*^EGFP-CreERT2/EGFP-CreERT2^ mice. Data are means ± SEMs (n=4 samples/genotype). One-way ANOVA [(F(2,9)=129.4, *p*<0.0001] followed by Tukey’s *post-hoc* test showed \*\*\**p*<0.001, C57 *vs. Htr2a*^EGFP-CreERT2/+^; *\*\*\*\*p*<0.0001, C57 *vs. Htr2a*^EGFP-CreERT2/EGFP-CreERT2^. (C) LSD-mediated HTR2A downregulation: Mice (C57 and *Htr2a*^EGFP-CreERT2^) were injected with vehicle or LSD (0.5 mg/kg, i.p.) for 5 consecutive days and then were euthanized 24 hr later. The whole cortices were dissected followed by receptor purification using Wheat-Germ beads pull-down. Western blot against anti-HTR2A antibody detected the receptor expression levels. Data represent the means ± SEMs (n=5-7). Two-way ANOVA: treatment [F(1,21)=27.11, *p*<0.0001], genotype [F(1,21)=24.09, *p*<0.0001], and treatment by genotype interaction [F(1,21)=0.1832, *p*=0.673] (overall genotype effects (black): ****p<0.0001, C57 *vs. Htr2a*^EGFP-CreERT2/EGFP-CreERT2^; overall treatment effect (purple): ^####^*p* <0.0001, vehicle *vs.* LSD). The schematic was created with https://www.biorender.com/.

It is well known that LSD and other psychedelic drugs induce substantial HTR2A down-regulation following repeated *in vivo* administration (Buckholtz et al., 1988; Buckholtz et al., 1990; McKenna et al., 1989). Here, C57 and *Htr2a*^EGFP-CreERT2^ mice were treated with 0.5 mg/kg LSD (i.p.) for 5 consecutive days and 24 hr later the whole cerebral cortex was removed for receptor purification and western blot analyses as previously described (Yadav et al., 2011) (Fig 4C). After 5-days of daily LSD administration, endogenous HTR2A in C57 and HTR2A-EGFP-CT fusion receptors from *Htr2a*^EGFP-CreERT2^ mice were downregulated to similar extents compared to the vehicle-treated group (p<0.0001). These results indicate that the EGFP insertion in the C-terminus of *Htr2A* does not affect *in vivo* receptor down-regulation response to LSD which is in agreement with prior studies using a similar strategy to investigate internalization and down-regulation *in vitro* (Bhatnagar et al., 2001; Bhatnagar et al., 2004; Gray et al., 2001).

### Characterization and application of inducible CreERT2-driving HTR2A-expressing cells

#### Validation of inducible CreERT2 activity and HTR2A-tdTomato cell labeling via the CreERT2 driver

The bi-cistronic design of *Htr2a*-EGFP-CT-IRES-CreERT2 produced a HTR2A-EGFP-CT fusion protein and CreERT2 recombinase. The expression of CreERT2 recombinase under *Htr2a* promoter provided a genetic manipulation platform via Cre-dependent reporter (Ai9 mice here) for specifically expressing tdTomato reporter in HTR2A containing cells. HTR2A-tdTomato expressing neurons represent HTR2A-positive neurons, while the HTR2A-EGFP-CT fusion protein represents HTR2A receptor itself (Fig 5A). Whole brain mapping of induced tdTomato expression (Suppl. Fig S4) was compared with HTR2A-EGFP-CT expression in slices and cleared whole brain. The result from whole brain cleared tissue showed HTR2A-EGFP-CT and tdTomato to have quite similar distributions in the 3D rendering of coronal, horizontal, and sagittal views, respectively (movie S1, S2, S3 and S4). For controls, the *Htr2a*^EGFP-CreERT2/+^x Ai9 and C57x Ai9 mice were treated with vehicle (corn oil) or tamoxifen to determine whether tdTomato expression was specifically regulated by tamoxifen for driving CreERT2 recombinase. Only *Htr2a*^EGFP-CreERT2/+^x Ai9 mice with tamoxifen treatment showed tdTomato expression, whereas none of the other treated groups showed this expression (Suppl. Fig S5).

**Fig 5.**
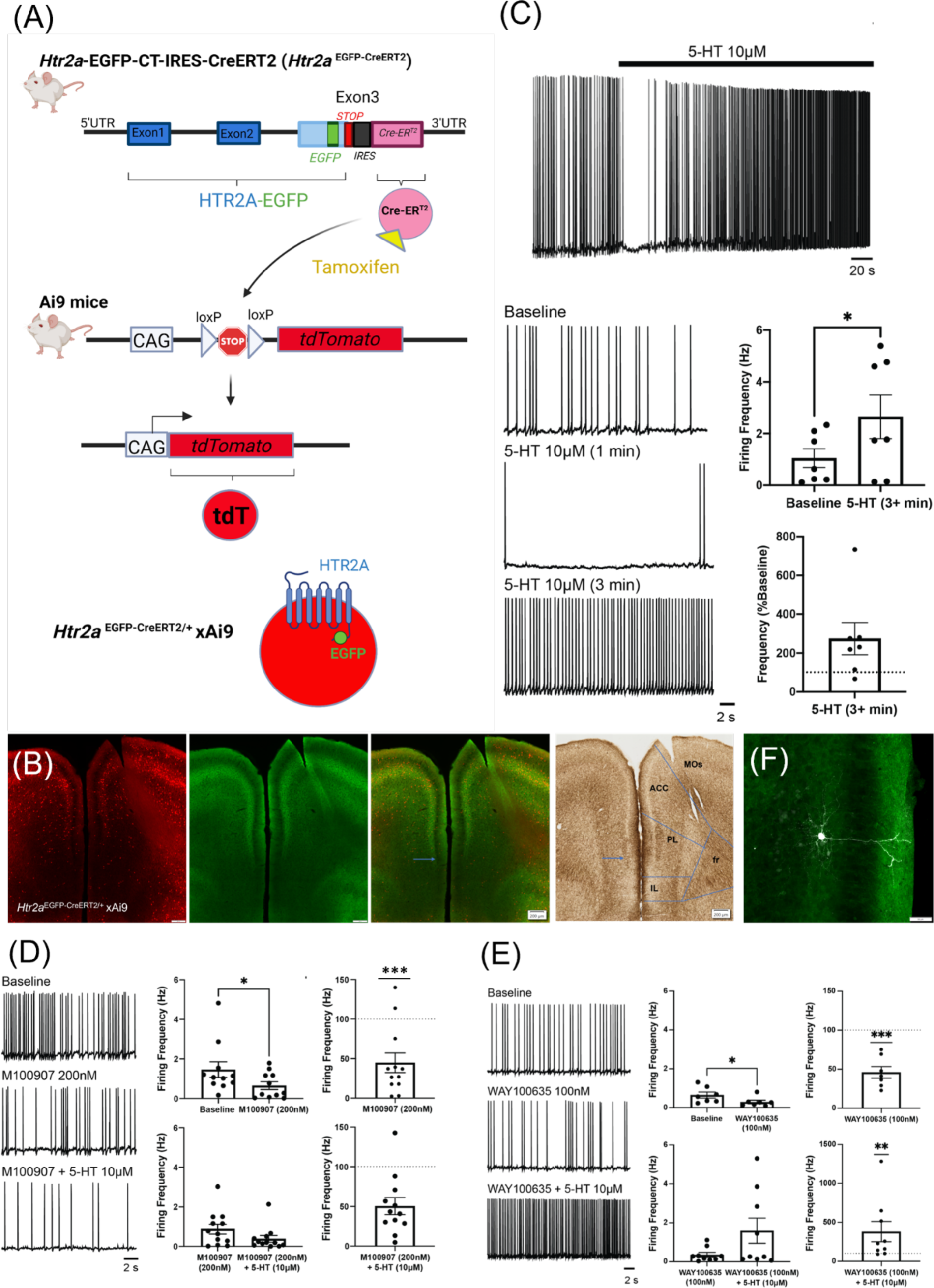
Inducible tdTomato visualization and Electrophysiological effects on neuronal firing in HTR2A-expressing neurons. (A) *Htr2a*^EGFP-CreERT2^ mice crossed with the Cre-dependent reporter line (Ai9 mice here) were used to visualize HTR2A expressing cells with tdTomato expression. The bicistronic design of Htr2a-EGFP-IRES-CreERT2 produced a HTR2A-EGFP-CT fusion protein and CreERT2 recombinase. Tamoxifen activates CreERT2 recombinase, leading to Cre-loxP recombination and then turning on tdTomato expression. The schema was created with https://www.biorender.com/. (B) The expression pattern of tdTomato (red) and HTR2A-EGFP-CT fusion proteins (green) in the mPFC subregion were identified using immunohistochemistry staining for HTR2A expression patterns and AchE staining for the cytoarchitecture. MOs: secondary motor cortex; AAC: anterior cingulate area; PL: prelimbic cortex; IL: infralimbic cortex; and fr: anterior forceps (C) L5a pyramidal neurons were used for recording neuronal firing activity during focal 5-HT (10 µM) application. Addition of 5-HT resulted in an initial decrease in firing followed by increased firing shown 3 min after application. Data are presented as means ± SEMs (n=7) and were analyzed using paired t-test to compare basal activity and firing after 5-HT application, [t(6)=3.177, p=0.0191] (*p<0.05) and one sample t-test was used for normalization of basal activity for the 5-HT response, p=0.0469 (*p<0.05). (D) Representative recording of HTR2A^+^/L5a pyramidal neuronal firing after M100907 (200 nM) and 5-HT (10 µM). M100907 effects on firing were analyzed using paired t-test, [t(10)=2.667, p=0.0236] (*p<0.05 *vs.* baseline) and normalized M100907 effects on firing were analyzed with one-sample t-test, [t(11)=4.421, *p*=0.001] (***p=0.001). Data are represented as means ± SEMs (n=11). The effects of M100907 + 5-HT on firing were analyzed with paired t-test, [t(11)=2.036, p=0.067] and normalized effects were analyzed using one-sample t-test. Data are presented as means ± SEMs (n=12). (E) Representative recording of HTR2A^+^/L5a neuronal firing after administration of WAY100635 (100 nM) and 5-HT (10 μM). WAY100635 effects on firing were analyzed with paired t-test, [t(6)=3.115, p=0.021] (*p<0.05, *vs* baseline) and normalized effects on firing were analyzed with one-sample t-test, [t(6)=7.228, p=0.0004] (***p<0.001). Data are presented as means ± SEMs (n=7). WAY100635 + 5-HT effects on firing were analyzed with paired t-test, [t(8)=2.159, p=0.0629] and normalized WAY100635 + 5-HT effects on firing were analyzed with one sample t-test *p*=0.0039 (**p=0.01). Data are represented as means ± SEMs (n=9). (F) A representative biocytin-labeled L5 pyramidal neuron showed its location in the HTR2A-EGFP-CT positive cortical band layer in the dorsal prelimbic cortex. After recording, the patched neuron was injected with biocytin. The brain sections were stained with an anti-GFP antibody to locate the L5a cortical band. Images were taken under 20X objective with an Olympus confocal microscope.

#### 5-HT effects on neuronal firing in HTR2A-positive L5a mPFC pyramidal neurons

We leveraged the visualization properties of tdTomato-expressing neurons to conduct electrophysiological whole-cell recording in *ex vivo* brain sections, allowing us to study 5-HT-mediated responses in HTR2A-containing pyramidal neurons in the cortical L5a of the mPFC. We first characterized the mPFC subregion distributions of HTR2A-EGFP-CT and HTR2A-tdT in *Htr2a*^EGFP-CreERT2/+^x Ai9 mice. The cytoarchitecture features by AchE staining revealed an AchE-weak zone in the ventral PL and IL. Nevertheless, HTR2A-EGFP-CT was robustly expressed in cortical L5a of the ACC and dorsal PL, but not in the ventral PL and IL; tdTomato-positive neurons also had similar subregion distributions (Fig 5B). Thus, we specifically targeted tdTomato-expressing neurons in the ACC and dorsal PL (dPL) for electrophysiological recordings.

Recent research has questioned whether 5-HT can activate HTR2A *in vivo* (Vargas et al., 2023) given that the HTR2A protein has been shown to reside both in intracellular compartments (Cornea-Hebert et al., 2002; Gelber et al., 1999) and at post-synaptic sites (Abbas et al., 2009; Jones et al., 2009; Miner et al., 2003). To test this hypothesis, we performed whole-cell current-clamp recordings of neuronal firing in visually-identified tdTomato-expressing neurons in L5a of the ACC or dPL. Focal application of 5-HT (10 µM) produced a biphasic response, with an immediate but transient reduction in firing followed by a sustained increase in firing frequency (Fig 5C). When firing was normalized to individual baseline values, 5-HT increased the firing rate by ∼230% (*p*=0.0469; Fig 5C). The HTR2A antagonist M100907 was used to determine whether the HTR2A mediated the change in 5-HT-mediated firing. M100907 (200 nM) significantly decreased the frequency of firing (*p*=0.0236, Fig 5D) to ∼50% of baseline values (p=0.001; Fig 5D). Subsequent application of 5-HT in the presence of M100907 produced no significant change in sustained firing frequency, indicating the HTR2A-mediated the 5-HT-induced increase in sustained firing in HTR2A-expressing cortical L5a ACC/dPL neurons.

Since the HTR2A and HTR1A have been reported to reside in the same neurons and because HTR1A agonists exert an inhibitory effect on neuronal activity in the rat dorsal raphe and PL cortical areas (Martin-Ruiz et al., 2001), we examined the role of HTR1A in HTR2A-expressing cortical L5a ACC/dPL neurons. Focal application of the HTR1A antagonist WAY100635 (100 nM) significantly decreased the firing frequency (*p*=0.021; Fig 5E) to ∼30% of normalized baseline values (*p*=0.0004). Subsequent application of 5-HT in the presence of WAY100635 increased the firing frequency, and this increase was statistically significant when firing frequencies were normalized to baseline values (*p*=0.0039). These results indicate that the HTR1A is not involved in the sustained increases in firing produced by 5-HT in L5a HTR2A-expressing pyramidal neurons in the ACC/dPL.

Following the electrophysiological recordings, the recorded tdTomato neuron was filled with biocytin, which facilitated the verification of its anatomical location. The location of the biocytin-labeled tdTomato neuron was confirmed subsequently using a fluorescent streptavidin and co-staining with a GFP antibody for HTR2A-EGFP-CT labeling. This co-staining revealed the soma of the recorded neuron resided in the GFP-positive layer, L5a within the ACC or dPL. Moreover, the biocytin filling of the entire neuron revealed its pyramidal neuron morphology (Fig 5F). During the recording process, each recorded neuron was imaged under a fluorescence microscope to provide a visual representation of its location within the ACC or dPL. The schema depicting the location of individual recorded neurons is shown in Supplementary Figure S6.

### Generation of additional reporter lines and humanized HTR2A mice

We generated two additional *Htr2a* constitutive Cre mouse lines. The first one is the *Htr2a*-IRES-Cre mouse line (*Htr2a*^Cre^) that was designed with aa native *Htr2a* coding sequence followed by IRES-Cre in the 3’UTR. The *Htr2a* genomic coding sequence was not modified in this mouse when inserting the IRES-Cre into the 3’UTR, in contrast to the already described *Htr2a*-EGFP-CT-IRES-CreERT2 mouse. The second line is a humanized line, *Htr2a*-A242S-EGFP-CT-IRES-Cre (*Htr2a*^A242S-EGFP-Cre^), which was designed so that a single point mutation in alanine residue 242 was replaced by a serine from the human *HTR2A* and the EGFP was inserted into the C-terminus followed by the IRES-Cre sequence (Fig 6A). This single point mutation has been shown previously to enhance the affinity and potency of a variety of psychedelic and non-psychedelic drugs to levels similar to those seen at the human receptor (Johnson et al., 1994; Kim et al., 2020). Receptor distributions were visualized with an anti-HTR2A antibody for HTR2A in the *Htr2a*^Cre^ mice (Suppl. Fig S7A) and an anti-GFP antibody for HTR2A-A242S-EGFP-CT in the *Htr2a*^A242S-EGFP-Cre^ mice (Suppl. Fig S7B), respectively. Protein expression of the HTR2A was demonstrated both by immunohistochemistry (Fig 6B) and by Western blot (Fig 6C) assays among the four mouse lines (*Htr2a*^+/+^, *Htr2a*^Cre^, *Htr2a*^EGFP-CreERT2^, and *Htr2a*^A242S-EGFP-Cre^). The HTR2A protein distribution from the three *Htr2a* knock-in mice showed similar expression patterns as in the *Htr2a*^+/+^ mice (Fig 6B). Samples from *Htr2a*^EGFP-CreERT2^ and *Htr2a*^A242S-EGFP-Cre^ mice found their mutant HTR2A-EGFP-CT and HTR2A-A242S-EGFP-CT fusion proteins to shift around 100 KD, while endogenous HTR2A proteins from *Htr2a*^+/+^ and *Htr2a*^Cre^ mice showed the expected original HTR2A molecular weight (between 50-75KD) (Fig 6C).

**Fig 6.**
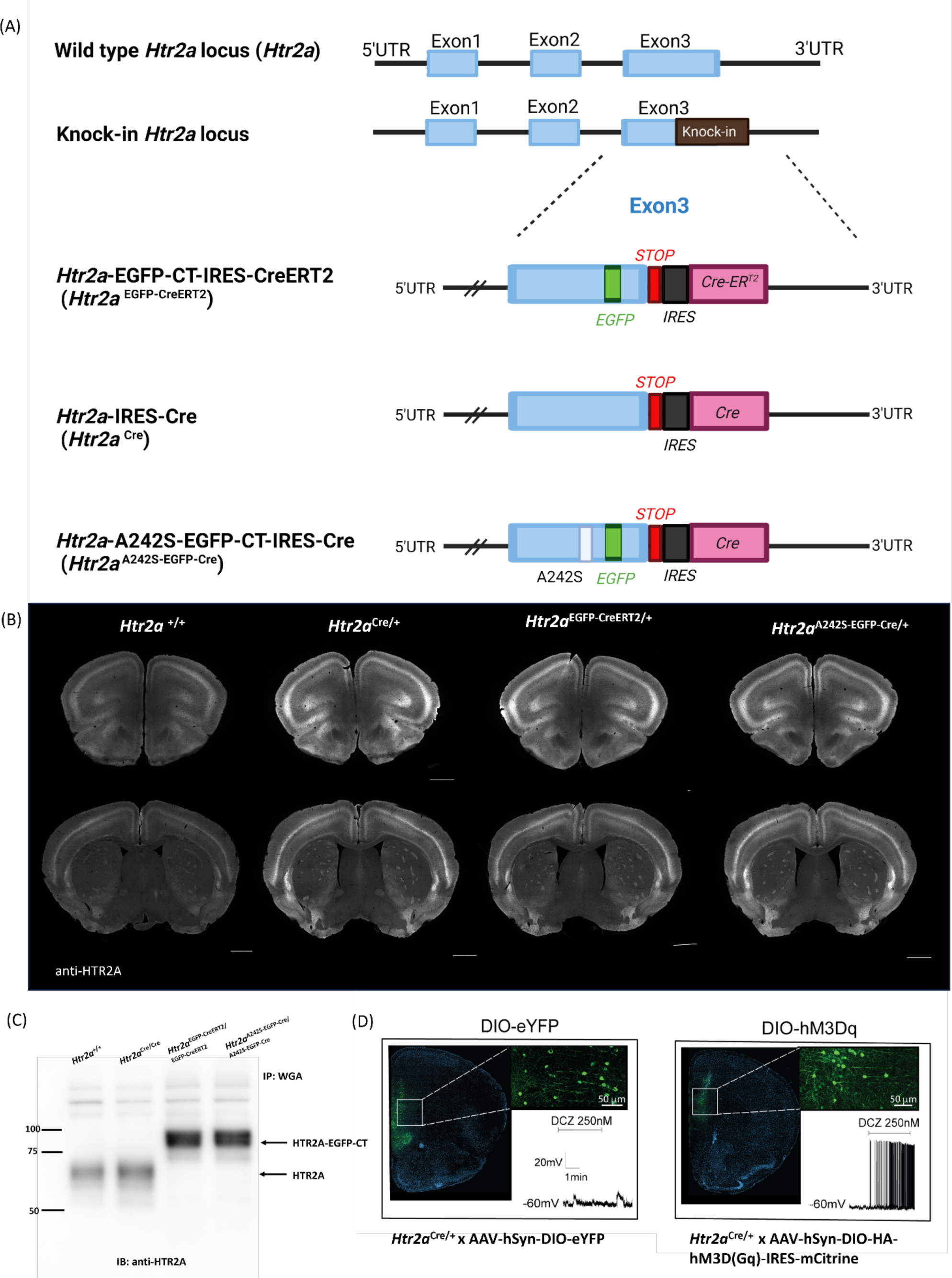
A series of HTR2A transgenic mouse lines. (A) The schema depicts the transgenic designs of each mouse line. As described previously, the *Htr2a*-EGFP-CT-IRES-CreERT2 (*Htr2a*^EGFP-CreERT2^) was inserted with EGFP in the C-terminus of the *Htr2a* followed by adding the IRES-CreERT2 sequence after exon 3 as an inducible *Htr2a* mouse line. *Htr2a*-IRES-Cre mouse line (*Htr2a*^Cre^) was engineered IRES-Cre sequence at the end of exon 3. A humanized mouse line, *Htr2a*-A242S-EGFP-CT-IRES-Cre (*Htr2a*^A242S-EGFP-Cre^), was created with one point mutation at the alanine 242 residue of the murine *Htr2a* to the serine residue of the human *HTR2A* and this was followed at the murine 452 residue by an EGFP insertion; the IRES-Cre was inserted after the exon 3 coding sequence. In the diagrams, the blue boxes represent exons of the murine *Htr2a* gene; the green box is *Egfp*; the red box is the stop codon. The gray box is the IRES (internal ribosome entry site) followed by a pink box for Cre or CreERT2 recombinase. The white box is the A242S point mutation. The schematic was created with https://www.biorender.com/. (B) HTR2A distribution in the brain among four mouse lines (*Htr2a*^+/+^, *Htr2a*^Cre/+^, *Htr2a*^EGFP-CreERT2/+^*, and Htr2a*^A242S-EGFP-Cre/+^). For this study, mice were perfused, and the brain sections were immunostained with an anti-HTR2A antibody to visualize the HTR2A distribution. (C) The HTR2A receptor from four mouse lines (*Htr2a*^+/+^, *Htr2a*^Cre/Cre^, *Htr2a*^EGFP-CreERT2/EGFP-CreERT2^*, and Htr2a*A242S-^EGFP-Cre/A242S-EGFP-Cre^). Here, mice were euthanized and then cortices were dissected followed by receptor purification using Wheat-Germ beads pull-down. Western blot against anti-HTR2A antibody to detect receptor expression level. In *Htr2a*^+/+^ and *Htr2a*^Cre/Cre^ mice, the HTR2A was detected between 50-75 KD; for EGFP insertion lines, the HTR2A-EGFP-CT fusion protein was detected around 100 KD. (D) Representative image of *Htr2a*^Cre/+^ mice expressing DIO-eYFP in the mPFC was stained with anti-GFP for eYFP signal amplification at 20x (left) and 63x magnification (left inset). Example electrophysiological trace of a HRT2A^+^ neuron expressing DIO-eYFP during bath application of DCZ. Representative image of *Htr2a*^Cre/+^ mice expressing DIO-HA-hM3Dq-IRES-mCitrine in the mPFC was stained with anti-GFP for mCitrine at 20x (right) and 63x magnification (right inset). Example electrophysiological trace of a HRT2A^+^ neuron expressing DIO-HA-hM3Dq-IRES-mCitrine during bath application of DCZ.

The distribution of the HTR2A-containging cells was examined also through recombinant expression of tdTomato. As Cre-recombinase activity is driven at various stages of development (Roth et al., 1991), many cortical neurons displayed tdTomato fluorescence in a pattern quite distinct from that of endogenous HTR2A expression (Suppl. Fig S7C). Cortical tdTomato expression patterns in constitutive *Htr2a*^Cre^ and *Htr2a*^A242S-EGFP-Cre^ mice crossed with Ai9 mice were quite different from the HTR2A expression pattern and tdTomato signals observed in the inducible *Htr2a*^EGFP-CreERT2^ x Ai9 mice, which may be useful for future fate mapping studies.

For constitutive Cre-expressing mice, virally-mediated transduction is frequently used for spatiotemporal Cre-loxP dependent recombination. Here, peripheral administration of PHP.eB AAV virus via retro-orbital injection was performed to test for brain-wide transduction. Two constitutive mouse lines (*Htr2a*^Cre^ x PHP.eB.FLEX.tdT and *Htr2a^A242S-EGFP-^*^Cre^ x PHP.eB.FLEX.tdT) displayed similar tdTomato distribution patterns, although these patterns were distinct from those of HTR2A receptors (Suppl. Fig S7D1). As a comparison, inducible *Htr2a*^EGFP-CreERT2^ mice were retro-orbitally injected with PHP.eB.FLEX.tdTomato to assess their tdTomato expression patterns. Interestingly, differences were observed in the tdTomato expressing patterns between the virally-mediated approach and the Ai9 reporter mice in the inducible mice (Suppl. Fig S7C and S7D1). For example, the viral approach revealed trace levels of the tdTomato signal in the striosome (CPu-s) and olfactory tubercle (TUO), whereas the signal was high in the induseum griseum (IG) and septohippocampal nucleus (SH) (Suppl FigS7D1). In the cortex, virally-mediated tdTomato expressing neurons were found in the deeper cortical layers L5b and L6 (minor), while Ai9-mediated inducible tdTomato was located in L5a and L6--but not in L5b (Suppl Fig S7D2). Despite the similar distribution of HTR2A receptors observed in all three mouse lines, different Cre-dependent approaches (reporter viruses and reporter mice) displayed dissimilar expression patterns of HTR2A-containing cells due to distinct Cre recombination occurring timing and efficiencies (Cre *vs*. CreERT2), and viral infection efficiencies (resulting in incomplete cell infected) as well as various promoters of Cre-dependent reporters. Notably, when using Cre-dependent reporters as a readout investigates HTR2A-mediated effects, it will be important to validate whether reporter expression patterns correspond with HTR2A receptor distributions in the same brain region.

As previously mentioned, the human HTR2A differs from the rodent version of the receptor by virtue of a single amino acid change in the binding pocket (S242A) which attenuates the activity of a variety of psychedelics (Kim et al., 2020) and non-psychedelic HTR2A agonists (Johnson et al., 1994). Accordingly, we created mutant mice in which murine Ala242 was mutated to the human serine residue (i.e., A242S). Initial behavioral studies were conducted to compare head twitch responses and PPI performance between the C57 and *Htr2a*^A242S-EGFP-Cre^ mice. In C57 animals, DOI elicited more robust head twitch responses than in the *Htr2a*^A242S-EGFP-Cre^ mice (*p*<0.001), whereas the converse was observed with psilocin (*p*=0.026) (Suppl. Fig S8A). In both genotypes, all psychedelics stimulated head twitch responses relative to the vehicle control (*p*-values<0.001), while the response to DOI was more pronounced than that to psilocin but only in the C57 mice (*p*=0.001). An examination of PPI revealed responses were higher in the *Htr2a*^A242S-EGFP-Cre^ than in the C57 mice (*p*=0.009) and all psychedelics were disruptive of PPI (*p*-values<0.001) (Suppl. Fig 8B). In addition, overall PPI responses were higher to psilocin than DOI (*p*=0.044). Genotype effects emerged for null activities (*p*<0.001), and treatment effects were evident (Suppl. Fig 8C). Compared to vehicle, null activities were enhanced by all psychedelics (*p*-values<0.016). Startle activities were higher in C57 than *Htr2a*^A242S-EGFp-Cre^ mice given the vehicle or psilocin (*p*-values<0.025) (Suppl. Fig 8D). In C57 animals, startle was higher to psilocin than to LSD (*p*=0.004), whereas in *Htr2a*^A242S-EGFP-Cre^ mice responses to DOI were enhanced over vehicle (*p*<0.001). Collectively, these results show that while overall responses to the different psychedelics are relatively uniform for PPI, there are some genotype differences in HTRs.

Biochemical studies in *Htr2a*^A242S-EGFP-Cre^ mice have found the HTR2A affinities as Kd values to be 2.38 ± 1.48nM and receptor expression levels as Bmax values to be 0.49 ± 0.2 pmole/mg using [^3^H]-ketanserin radioligand binding assays. Given the well-documented differences between human and rodent HTR2A psychedelic pharmacology (Kim et al., 2020), the A242S mouse line will likely prove useful for interrogating the action of psychedelic drugs *in vivo*.

We also compared the distribution of HTR2A in neurons and those labelled with the Thy1-GFP reporter which previously had been used to quantify spine formation after psychedelic drug administration (Shao et al., 2021; Vargas et al., 2023). For these studies, our *Htr2a^EGFP-^*^CreERT2^ x Ai9 mice were crossed with Thy1-GFP mice. As can be seen in the cortex (Suppl. Fig S9), Thy1-GFP neurons did not co-localize with the tdTomato-expressing neurons in L5a. Only rare Thy1-GFP-positive/HTR2A-tdTomato-positive neurons were observed in L5b/L6 (tdT/Thy1-GFP in L5b/L6 of the ACC/dPL cortex: 1.23 ±1% from 445 neurons of 3 animals). Given the lack of 1:1 co-localization with Thy1-GFP, our results imply that studies on psychedelic drug-induced plasticity using this reporter mouse may not faithfully quantify spine formation in HTR2A-expressing neurons. Thy1-GFP mice are useful, however, to determine the effects of psychedelics and other drugs on non-HTR2A-expressing neurons.

### Designer Receptor Exclusively Activated by Designer Drugs (DREADD) application in *Htr2a*^Cre^ mice

We also manipulated the activity of HTR2A-expressing neurons with a chemogenetic actuator (deschloroclozapine, DCZ) in *Htr2a*^Cre^ mice with local viral injection of DREADD into the mPFC. *Htr2a*^Cre/+^ mice that expressed excitatory DREADD (DIO-HA-hM3Dq-IRES-mCitrine) displayed a significant depolarization response to bath application of DCZ, whereas HTR2A^+^ neurons expressing DIO-eYFP did not show a physiological response (Fig 6D). Consequently, DCZ was shown to specifically activate hM3Dq^+^/HTR2A^+^ neurons. This illustrates how *Htr2a*^Cre/+^ mice prove to be valuable in chemogenetic studies for revealing the functions of specific cell populations expressing HTR2A.

## DISCUSSION

Psychedelic drugs have emerged as transformative therapeutics for a large number of neuropsychiatric disorders (Roth et al., 2023) and all classical psychedelics induce their typical responses in mice (Gonzalez-Maeso et al., 2007; Keiser et al., 2009a) and humans (Vollenweider et al., 1998; Preller et al., 2018) via HT2ARs (Gumpper et al., 2023). Despite decades of study, however, we lack key technologies to interrogate the role of the HTR2A in the actions of these drugs (McClure-Begley et al., 2022; Kwan et al., 2022). Here we provide an open-source suite of validated genetically-engineered mouse models (AMIS, https://amis2.docking.org/) which provides platforms to interrogate the species-specificity, cellular signaling, neural circuitry, and behavioral actions of psychedelic drugs.

Previous studies have utilized distinct and orthogonal approaches to examine the distribution of HTR2A by receptor autoradiography and *in situ* hybridization, as well as by studying transgenic mice. The early autoradiography work in rats from the Palacios group (Pazos et al., 1985) and others (Roth et al., 1987) shows a similar distribution of the HTR2A in the brain as the results with our engineered mouse models. HTR2A expression was high in the claustrum, olfactory tubercle, piriform cortex, and what was (erroneously) designated layer 4 of the cortex (Pazos et al., 1985). Moderate levels were seen in the striatal complex and certain nuclei of the amygdala, but outside of the forebrain, in regions such as the thalamus, midbrain, and lower brainstem, HTR2A was not present or seen in low levels (Pazos et al., 1985).

The distribution of the HTR2A has been studied also in human brain using [^3^H]-ketanserin, [^3^H]-mesulergine, [^3^H]-LSD, and [^3^H]-spiperone (Pazos et al., 1987). Of these ligands, ketanserin is the most selective for the HTR2A and the expression patterns with this radioactive ligand more closely correspond with patterns we observed using our mouse *Htr2a*^EGFP-CreERT2^ line. The similarities were evident in comparisons of high expressing cortical areas; intermediate expression was seen in the claustrum and striatum as well as restricted areas of the amygdala, and very low or absent labeling in the thalamus, brainstem, and cerebellum.

In our knock-in mice, HTR2A-EGFP-CT showed a similar trend. High expression levels were observed throughout the cortex, in the striatal complex, and claustrum. Other forebrain loci with high expression included CA3 in the ventral (temporal) hippocampus and the lateral nucleus of the amygdala. In more posterior areas high expression was observed in the medial mammillary nucleus and both the basis pontis and the inferior olive. HTR2A-EGFP-CT expression of was either absent or seen in trace levels in diencephalic (thalamus and hypothalamus) and midbrain areas, as well as the ponts and medulla. Notable was the lack of expression in the brain raphe nuclei, with the exception of low levels in the raphe obscurus and raphe magnus (see Table 1).

The GENSAT resource (Gong et al., 2003; Heintz, 2004) (Schmidt et al., 2013) provides transgenic mice for interrogating HTR2A circuits and function (see *Htr2a*-EGFP-DQ118, *Htr2a*-Cre-KM207 (FigS1A) and *Htr2A*-Cre-KM208 (FigS1B) (http://www.gensat.org/imagenavigatorfileselection.jsp?gene.id=575&allages=true)) among others. Unfortunately, these lines do not faithfully recapitulate the endogenous distribution of the HTR2A as has been seen with other BAC-transgenic mice (Matthaei, 2007). Despite this limitation, some groups have been used for HTR2-related research, such as electrophysiological studies (Weber et al., 2010), retina-related studies (Simmons et al., 2020)and circuitry-related research. The resultant data should be interpreted with caution since the distribution of HTR2A in the *Htr2a* BAC mouse lines are distinct from that of the endogenous receptor based on studies using receptor autoradiography (Pazos et al., 1985; Roth et al., 1987), immunohistochemistry (Willins et al., 1997a; Backstrom et al., 1997; Miner et al., 2003; Mengood et al., 1990), *in situ* hybridization (Wright et al., 1995) and our mouse lines.

We also observed *Htr2a* transcript distribution, which showed some modest differences compared with HTR2A-EGFP-CT protein expression (Suppl. Fig S10A). *Htr2a* transcripts displayed cortical distribution from L2/3, L5 to L6, while HTR2A-EGFP-CT was mainly expressed in L1, L5a and L6. These results suggest some level of dissociation between the mRNA and protein levels of the HTR2A in specific cortical layers as should be expected since the HTR2A protein is localized primarily to processes rather than cell bodies. As a follow-up for mRNA expression, we examined whether *Egfp*-tagged *Htr2a* transcripts were normally expressed in *Lamp5*-positive neurons (a marker for L2/3 and L5b), parvalbumin inhibitory interneurons, and *Cplx3*-positive neurons (a marker for L6b). We found *Htr2a-Egfp* transcripts were co-localized with *Lamp5* in L2/3 and L5b, as well as with the interneuron marker parvalbumin and with *Cplx3* in L6b (Suppl. Fig S10B). These results are consistent with the Allen Institute for Brain Science Brain Map for the distribution of *Htr2a* mRNA in cortical L2/3, L5, and L6 (http://casestudies.brain-map.org/celltax#section_explorea).

The data we have obtained on the localization of HTR2A-EGFP-CT may offer some clues as to the sites of action of psychedelic drugs. Doss et al (Doss et al., 2022)have recently reviewed and evaluated the three major current hypotheses on brain circuits that underlie the psychedelic actions of psychedelic drugs. All three involve a large expanse of cortical areas, but otherwise differ.

One of these models is a cortico-striato-fugal circuit that returns to (and goes beyond) the originating cortical site. This Cortico-Striatal-Thalamo-Cortical (CSTC) model posits a key role for thalamic projections from pallidal sites that receive striatal inputs, with prominent roles for the thalamic reticular nucleus and the mediodorsal nucleus (MD). In an immunohistochemical study (Rodriguez et al., 2011) commented on the presence of low but detectable levels of HTR2A in the thalamic reticular nucleus, the antibody used was not characterized. Using in situ hybridization histochemistry, Wright et al. (1995) observed very low levels of the *Htr2a* transcript in the reticular or mediodorsal thalamic nuclei. Almost all other studies, including the autoradiographic studies using radiolabeled MDL100,907, LSD, and DOI (McKenna et al., 1989) (Lopez-Gimenez et al., 2013) (Lopez-Gimenez et al., 1999) (Lopez-Gimenez et al., 1997) as well as our own findings, note that the thalamus stands out by virtue of lacking significant HTR2A.

A second major model of the brain system(s) that mediates psychedelic drug action is the Relaxed Beliefs Under Psychedelics (REBUS) model. This circuit has fewer nodes than the CSTC model, and critically posits a hippocampal-cortical processing loop, through which the ability of cortical areas to modulate hippocampal function is diminished by the actions of psychedelics. One group (Bombardi, 2012) using a poorly characterized antibody, reported that HTR2A-like immunoreactivity is present in most areas of the hippocampal and parahippocampal area, from the entorhinal cortex to the dentate gyrus. Wright et al. (1995) found very low levels of *Htr2a* mRNA in the CA fields of the hippocampus, but did note that the transcript was present in high abundance in the ventral subiculum, the dentate gyrus, and the entorhinal cortex. We observed that HTR2A-EGFP-CT was present in high abundance in the lateral (but not medial) entorhinal cortex and the CA3 pyramidal cell area in ventral aspects of the temporal (but not medial) hippocampal formation.

The final model evaluated by Doss et al. (2022) is the cortico-claustro-cortical (CCC) model. The claustrum is a thin structure just lateral to the external capsule that enjoys wide-spread reciprocal connections with many cortical areas (Mathur et al., 2009) (White et al., 2017) (Wang et al., 2017) and as such is capable of serving as a central node regulating cortical activity. The CCC model hypothesizes that psychedelic drugs uncouple functional connectivity of the claustrum and cortex, thereby attenuating cortical networks. Moderately dense HTR2A-EGFP-CT expression is seen throughout the extent of the claustrum.

The mouse lines we have developed as a resource for the scientific community should be of considerable utility in testing models of brain circuits subserving psychedelic action. In particular, our data and those of other investigators point to the REBUS and CCC models as being the most plausible.

We also examined several behaviors that have been linked to psychedelic drug actions in rodents (Corne and Pickering, 1967; Glennon et al., 1984; Kyzar et al., 2016; Rodriguiz et al., 2021). For instance, drug-induced changes in locomotor activity have been reported with LSD, DOI, and psilocin (Halberstadt et al., 2011; Halberstadt et al., 2009; Rodriguiz et al., 2021). The HTR2A has been implicated in the modulation of locomotor activity in rodents (Halberstadt et al., 2009; Rodriguez et al., 2021) and because this receptor is expressed in the striatum, HTR2A agonists could modulate the dopaminergic and serotonergic systems to affect motor activity (Halberstadt et al., 2011; Rodriguiz et al., 2021). Notably, the selective HTR2A antagonist M100907 blocks these effects (Rodriguiz et al., 2021). Due to their robust polypharmacology, psychedelics like LSD potently activate several dopamine receptors (Kroeze et al., 2015) and this dopaminergic activity could contribute to these locomotor responses.

The head twitch response is a validated animal model for identifying psychedelic drugs in rodents (Glennon et al., 1984) and it is mediated by the activation of HTR2A in the mPFC (Willins et al., 1997b). Previous studies have demonstrated that the LSD-, DOI-, DMT-, and psilocybin-induced head twitch responses in rodents are blocked by HTR2A antagonists (Glennon et al., 1983) and by genetic deletion of *Htr2a* in mice (Gonzalez-Maeso et al., 2007; Keiser et al., 2009). Another proxy for the actions of psychedelic drugs is PPI which is abnormal in schizophrenic patients (Mena et al., 2016) and is disrupted in both humans (Gouzoulis-Mayfrank et al., 1998) and rodents (Halberstadt et al., 2010) by psychedelic, but not by non-psychedelic HTR2A agonists. In behavioral tests of our HTR2A reporter mice, we used psychedelic drugs as positive controls because they induce hallucinations in humans and are reported to have potential therapeutic benefits (Roth et al., 2023). We observed LSD and DOI stimulated motor activity, head twitches, retrograde walking, and produced disruption of PPI; psilocin increased HTR and disrupted PPI. Taken together, our results indicate that our engineered mouse models display typical responses to psychedelic drugs using a variety of behavioral readouts.

In electrophysiological studies, 5-HT acted on genetically-identified L5a pyramidal neurons in the mPFC, producing a biphasic effect wherein a brief inhibition of firing (1 min) was immediately followed by a prolonged HTR2A-mediated excitatory effect in the late phase (>3 min). The HTR1A antagonist partially reversed the 5-HT mediated inhibitory effect on firing, while the HTR2A antagonist completely blocked 5-HT-mediated increase in firing. This result is consistent with previous findings showing opposite effects of HTR1A (hyperpolarization) and HTR2A (depolarization) in single cell activities (Martin-Ruiz et al., 2001). Thus, the HTR2A plays a major role in 5-HT-induced increases in neuronal activity in HTR2A^+^/L5a pyramidal neurons in the mPFC. These findings are important because of a recent report suggesting that many HTR2A agonists may exert their actions via intracellular rather than plasma membrane receptors (Vargas et al., 2023). As we recently reported (Schmitz et al., 2022), psychedelic drugs increase the firing of genetically-identified HTR2A neurons in a manner similar to that reported here for 5-HT. As 5-HT is extracellularly applied, our results are consistent with the hypothesis that the psychedelic actions of HTR2A agonists are mediated by direct actions of pyramidal neuron firing via surface HTR2A receptors.

With respect to the HTR2A mouse lines we have generated, they can be used as an integrated toolset for studying psychedelic-mediated mechanisms. In the present study, we demonstrated the whole brain mapping of HTR2A expression, psychedelic-mediated HTR2A behavioral pharmacology, HTR2A-mediated receptor downregulation, and 5-HT effects on HTR2A-mediated neuronal firing. Our mouse toolkit thus provides an invaluable platform for investigating HTR2A-mediated responses underlying psychedelic drug mechanisms, ranging from receptor trafficking, signal pathways, neuronal circuitry, and behavior.

## ACKNOWLEDGEMENTS

The mu opioid receptor antibody was a generous gift from Dr. Lee-Yuan Liu-Chen, Center for Substance Abuse Research of Temple University. We wish to thank Mr. Christopher Means, Ms. Sarah Page Steffens, Ms. Julia Hoyt, and Ms. Ann Njorge for helping to conduct the behavioral studies; Ms. Jiechun Zhou for breeding, genotyping, and maintaining the mice for behavioral studies; Dr. Vladimir Pogorelov for preparing the C57BL/6J and HTR2A transgenic mice for morphology studies; and Dr. Ramona Rodriguiz for organizing the behavioral data and helping to statistically analyze these results; all at Duke University. Some of the behavioral experiments were conducted with equipment and software purchased with a North Carolina Biotechnology Center grant. Confocal microscope images were taken at the UNC Neuroscience Microscopy Core (RRID:SCR_019060), supported, in part, by funding from the NIH-NINDS Neuroscience Center Support Grant P30 NS045892 and the NIH-NICHD Intellectual and Developmental Disabilities Research Center Support Grant P50 HD103573. Use of the Marsico Hall, Cystic Fibrosis Center Olympus VS120 slide scanner and file server is appreciated. CRISPR-made transgenic mice were generated at the UNC Animal Model Core Facility. This study was funded by Department of Defense DARPA grant (DARPA-5822.e003) and by R37DA045657.

## Authorship contributions

Experimental design: Chiu, Deutch, Wu, Wetsel, Chen, Niehaus, Schmitz, Kocak, Llorach, Walsh, Scherer, Herman and Roth

Conducted experiments: Chiu, Wang, Bowyer, Schmitz, Huang, Kocak and Llorach Data analysis: Chiu, Deutch, Wetsel, Schmitz, Kocak, Liu and Sciaky

Transgenic mice design and maintenance: Kocak, DiBerto, Hua and English Writing manuscript: Chiu, Deutch, Schmitz, Kocak, Walsh, Herman, Wetsel, and Roth

## MATERIALS AND METHODS

### Animals

Adult male and female C57BL/6J (#000664), Ai9 mice (#007909), and Thy1-GFP-M mice (#007788) were from Jackson Labs (Bar Harbor, ME). In house, HTR2A series of knock-in mice [*Htr2a*-EGFP-CT-IRES-CreERT2 (*Htr2a*^EGFP-CreERT2^), *Htr2a*-IRES-Cre (*Htr2a*^Cre^) and *Htr2a*-A242S-EGFP-CT-IRES-Cre (*Htr2a*^A242S-EGFP-Cre^)] were used in this study. All mice were on a C57BL/6J genetic background. Mice were housed 3-5 per cage in a temperature- and relative humidity-controlled room with a 12:12-hr light/dark cycle. All animals were provided food and tap water *ad libitum*. Animal experiments for saturation binding assay, immunohistology, RNAScope, western blot and electrophysiology were performed according to the National Institutes of Health guidelines for the Care and Use of Laboratory Animals and with an approved protocol from University North Carolina at Chapel Hill Institutional Animal Care and Use Committee (IACUC).

All behavioral experiments were conducted with an approved protocol from the Duke University IACUC. All experiments were performed under relevant regulations and ARRIVE guidelines. Prior to behavioral testing, the C57BL/6J mice had been screened in the zero maze for anxiety-like behavior (Fukui et al., 2007), open field for locomotor activity, and prepulse inhibition for sensorimotor gating and mice that responded “normally” on all three tests were used to further backcross the knock-in mice onto a C57BL/6J genetic background (Rodriguiz et al., 2021). These C57BL/6J mice were bred for 2 generations with the knock-in mice to render them most similar in behavioral phenotype to their native inbred animals.

#### Knock-in mice design

We generated two mouse HTR2A lines, *Htr2a*-IRES-Cre and the *Htr2a*-EGFP-CT-IRES-Cre^ERT2^. We then produced an additional reporter line that humanized the HTR2A in its orthosteric binding site. The *Htr2a*-A242S-EGFP-CT-IRES-Cre line humanizes the ligand binding pocket by introducing the A242S as a single point mutation. We used the GRCm38/mm10 genome assembly for all annotations listed below. The *Htr2a* gene is on chromosome 14 and spans nucleotides 74640840-74709494.

EGFP was inserted into the C terminal tail of the receptor, flanked on both ends by a serine- and glycine-rich linker. This insertion occurs after residue 452, corresponding to a DNA insertion site after chr14:74706337. This location is ideal in that it is distal to the final helix of the receptor so as not to disturb protein folding and also distal to the PDZ domain so as to not disturb protein trafficking. The IRES-Cre or IRES-CreERT2 sequence was inserted directly following the coding sequence and preceding the 3’ UTR, corresponding to a genomic insertion after nucleotide chr14:74706397. The A242S point mutation was introduced by changing GCA at nucleotides 74705705-74705707 to TCC.

#### Production of CRISPR protein, gRNA, and donor vectors

For the production of recombinant Cas9 protein, a human codon optimized FLAG-Cas9 cDNA (Addgene 42230) was modified by C-terminal insertion of an additional nuclear localization signal and a 6-His tag, and cloned into the pET-28a(+) vector (Novagen/Sigma-Aldrich, St. Louis, MO, USA). Cas9 protein expression and purification were performed by the UNC Protein Expression and Purification Core Facility. The Cas9 protein was purified using a HisTrap Ni–NTA column followed by a SP cation exchange column (reference: http://www.pnas.org/content/113/11/2868), followed by size-exclusion chromatography. The final protein was stored in 20 mM HEPES (pH 7.5), 150 mM KCl, 1 mM DTT, 50% glycerol.

Cas9 guide RNAs in target regions of the mouse *Htr2a* locus were identified using Benchling software (www.benchling.com). Selected guide RNAs were cloned into a T7 promoter vector followed by T7 *in vitro* transcription and spin column purification, with elution in microinjection buffer [5 mM Tris-HCl (pH 7.5), 0.1 mM EDTA]. Functional testing was performed by co-transfecting a mouse embryonic fibroblast cell line with guide RNA and Cas9 protein. The guide RNA target site was amplified from transfected cells and PCR products were analyzed by T7 Endonuclease I digestion (New England Biolabs) or Sanger sequencing followed by ICE analysis (Synthego) to detect Cas9 cleavage and indel formation.

The guide RNAs selected for production of knock-in mice were:

*Htr2a*-EGFP-CT-IRES-Cre-ERT2: sg58T2 (5’-gACTAGGGAACCAACACT-3’) and sg77T (5’-GTGATGGACCGGATGCT-3’)

*Htr2a*-IRES-Cre: sg77T (sequence above)

*Htr2a*-A242S-EGFP-CT-IRES-Cre: 3g52T (5’-gGTCCTCATAGGCTCTTTTG-3’) and sg77T (sequence above)

Double-stranded supercoiled DNA donor plasmids were used to generate each insertion event. Donor plasmids were prepared with the HiSpeed Plasmid Maxi Kit (Qiagen). Eluted DNA was spot-dialyzed in a microinjection buffer. Sequences of donor plasmids for each allele are in the Appendix.

#### Transgenic mice production

All animal handling and experiments were conducted in accordance with the National Institutes of Health Guide for Care and Use of Laboratory Animals and as approved by the Animal Care and Use Committee of the University of North Carolina (UNC). All animals were housed in temperature- and humidity-controlled rooms under standard 12-h cycle lighting with food and water provided *ad libitum*. The mouse strain used to obtain the embryos was from C57BL/6J (Jackson Laboratory, Bar Harbor, ME, USA 000664).

C57BL/6J zygotes were microinjected with 400 nM Cas9 protein, 25 ng/μl of each guide RNA, and 20 ng/μl donor vector. In some cases, *in vitro* transcribed mRNA encoding codon-optimized Flp recombinase was included in the microinjection mix at 50 ng/μl to reduce tandem integration events by recombining the single FRT sites in tandem copies of the donor vector. To prepare the microinjection mix, guide RNAs were diluted in microinjection buffer, heated to 95°C for 3 min, and placed on ice prior to addition of Cas9 protein. The mixture was then incubated at 37°C for 5 min and placed on ice, after which the donor vector was added, and the mixture was held on ice prior to pronuclear microinjection. Microinjected embryos were implanted in recipient pseudopregnant B6D2F1/J females (#100006; Jackson Labs). Resulting pups were screened by PCR and sequencing for the presence of the correct insertion allele. One or more founders with the correct allele were mated to inbred C57BL/6J animals to transmit the modified allele through the germline. Offspring from a single founder line were selected for additional breeding to maintain and characterize the line.

### Drugs

The drugs used in the behavioral experiments were (±)-2,5-dimethoxy-4-iodoamphetamine hydrochloride (DOI; Bio-Techne Corp., Minneapolis, MN), (+)-lysergic acid diethylamide-(+)-tartrate (LSD), and psilocin (NIDA Drug Supply Program, Bethesda, MD). The vehicle was composed of N,N-dimethyllacetamide (final volume 0.5%; Sigma-Aldrich) that was brought to volume with 5% 2-hydroxypropoyl-β-cyclodextrin (Sigma-Aldrich) in water (Mediatech Inc., Manassas, VA). All drugs were administered (i.p.) in a 5 ml/kg volume. For electrophysiological experiments, serotonin hydrochloride and M100907 were purchased from Sigma-Aldrich and RS102221 was purchased from Tocris Bioscience (Ellisville, MO). Stock solutions were prepared in ultra-pure water or DMSO, stored at −20°C and diluted to working concentration in aCSF on the day of testing.

### In situ hybridization

The RNAscope experiment followed the ACDBio protocol (RNAscope Multiplex Fluorescent Reagent Kit v2 assay) with slight modifications. Fresh brains from *Htr2a*^+/+^ and *Htr2a*^EGFP-CreERT2^ mice were dissected and immediately embedding into O.C.T specimen matrix (Tissue-Tek® O.C.T Compound, Sakura ®Finetek). Brain sections at 18 µm thickness were cut using a cryostat and directly mounted onto slides (Fisherbrand^TM^ Superfrost^TM^ Plus microscope slides) followed by a 1 hr post-fixation with 4% paraformaldehyde (PFA) in PBS at 4°C. Brain sections were passed through a serial gradient ethanol dehydration step, followed by H_2_O_2_ treatment to quench endogenous peroxidase activity. The sections were next subjected to protease 3 digestion for 20 min. Mouse *Htr2a* (#401291-C3) and *Egfp* (#400281) probes were used for hybridization over 2 hr at 40°C in a humidity-controlled oven (HybEZII, ACDBio) and then the signal was amplified using Opal Dye570 for the *Egfp* probe and Opal Dye690 for the *Htr2a* probe. *Pvalb* (#421931-C2), *Lamp5* (#451071), and *Cplx3* (#467821-C3) were used as probes for interneuronal and cortical layer markers. Slides were counterstained with DAPI and mounted. Images were collected under an Olympus VS120 virtual slide microscope and Olympus FV3000RS Confocal Microscope (Olympus, Tokyo, Japan).

### Immunohistochemistry

C57, *Htr2a*^EGFP-CreERT2^, *Htr2a*^Cre^ and *Htr2a*^A242S-EGFP-Cre^ mice were crossed with the Cre-dependent reporter Ai9 strain. Mice were injected (i.p.) with vehicle (corn oil, Sigma) or 100 mg/kg tamoxifen (Sigma) in a 5 ml/kg volume from p39 to p42. At 14 days post injection (14 dpi), mice were euthanized at p56, and brains were dissected for further HTR2A expression surveys. In a separate experiment, C57, *Htr2a*^EGFP-CreERT2^, *Htr2a*^Cre^ and *Htr2a*^A242S-EGFP-Cre^ mice were retro-orbitally injected with AAV-PHP.eB-FLEX-tdTomato. Brains were collected later for analysis. The pAAV-FLEX-tdTomato was a gift from Edward Boyden (Addgene plasmid #28306, Addgene, Watertown, MA).

Mice were intracardially perfused with 4% PFA in PBS. Brains were harvested, post-fixed in 4% PFA/PBS overnight, and dehydrated in 30% sucrose/PBS until sinking. Brains were sectioned by cryostat at 40 µm. The free-floating brain sections were washed 3 times with 0.1% TX-100/PBS prior to a 1 hr incubation in blocking buffer (5% normal donkey serum in 0.4%TX-100/PBS) and then incubated overnight at 4°C with anti-goat GFP (1:1000, Rockland, #600-101-215), anti-rabbit NECAB1 (1:1000, Sigma, # HPA023629), anti-Parvalbumin (1:4000, Swant, #235), and anti-rabbit mu opioid receptor (MOR) (1:4000, a gift from Dr. Lee-Yuan Liu-Chen, Temple University; (Huang et al., 2008) (Wang et al., 2016)). The anti-rabbit HTR2A antibody (1:250, Neuromics, #RA24288) and anti-rat CTIP2 (1:1000, Abcam, #ab18465) primary antibodies were incubated at room temperature (RT) overnight. Brain sections were washed with 0.1% TX-100/PBS 3 times and then incubated with secondary antibodies (donkey anti-goat-Alexa Fluor® 488 for GFP primary antibody; anti-rabbit-Alexa Fluor® 647 for 5-HT2A, NECAB1, and MOR primary antibodies; and anti-rat-Alexa Fluor® 647 for CTIP2 at 1:1000 dilution; Jackson Immunoresearch, West Grove, PA). Tissue sections were imaged on an Olympus VS120 virtual slide microscope or Olympus FV3000RS Confocal Microscope (Olympus, Tokyo, Japan).

### Whole brain clearing and imaging by lightsheet microscopy

The C57 or *Htr2a*^EGFP-CreERT2^x Ai9 mice (treated with tamoxifen) were used in this study. Mouse brains were processed with a newly optimized protocol (U.Clear, Wang et al., *in submission*) to image both EGFP and tdTomato fluorescent proteins and the immunolabeling of GFP was performed in the whole mount. Perfused and 4% PFA post-fixed mouse brains were washed with PBS, de-lipidated by several washes (1, 2, 4 hr, and overnight) followed by 3 washes 1 day later with SBiP buffer [200 µM Na_2_HPO_4_, 0.08% sodium dodecyl sulfate, 16% 2-methyl-2-butanol, 8% 2-propanol in H_2_O (pH 7.4)] at RT. Next, brains were transferred into blocking B1n buffer (0.1% Triton X-100, 2% glycine, 0.01% 10 N sodium hydroxide, 0.008% sodium azide in H_2_O) and mixed at RT overnight followed by incubation at 37°C for 1 hr. Brains were washed twice at 1 and 2 hr with PTxwH buffer [0.1% Triton X-100, 0.05% Tween-20, 0.002% heparin (w/v), 0.02% sodium azide in PBS]. For immunolabeling HTR2A-EGFP-CT fusion proteins and goat anti-GFP primary antibody (1:300, Rockland, #600-101-215), diluted in PTxwH buffer, were added to brains. The brains were maintained 37°C with gentle rocking for 2 weeks. Excess antibodies were removed in 4 washes (1, 2, 4 hr, overnight) with PTxwH. Subsequently, brains were incubated with Alexa647-conjugated donkey anti-goat secondary antibodies (1:100, Jackson Immunoresearch, #705-607-003) diluted in PTxwH at RT with gentle rocking for two weeks and subsequently washed 4 times (1, 2, 4 hr, overnight) with PTxwH. Samples were fixed in 1% PFA at 4°C overnight, washed in PTxwH at RT (1, 2, 4 hr, overnight), then blocked in B1n at RT overnight, and washed (1, 2, 4 hr) in PTxwH at RT. Samples were then bleached in 0.3% H_2_O_2_ at 4°C overnight, washed 3 times in 20 mM PB (16 mM Na_2_HPO_4,_ 4mM NaH_2_PO_4_ in H_2_O) at RT for 2 hr. For further de-lipidation, samples were immersed in SBiP buffer 4 times (1, 2, 4 hr, overnight), and then extra 6 times each for 1 day. Next, brains were washed twice (2 and 4 hr) in 20 mM PB buffer and finally twice in PTS solution (25% 2,2’-thiodiethanol/10 mM PB) for 2 hr and overnight, then equilibrated with 75% histodenz buffer (#AXS-1002424; Cosmo Bio USA) with the refractive index adjusted to 1.53 using 2,2’-thiodiethanol. Samples were stored at −20 °C until acquisition. The cleared brain samples were imaged horizontally with tiling using the LifeCanvas SmartSPIM lightsheet microscope. 561/647 nm lasers were used for GFP/RFP/IHC imaging with the 3.6×/0.2 detection lens. Lightsheet illumination was focused with a NA 0.2 lens and axially scanned with an electrically-tunable lens coupled to a camera (Hamamatsu Orca Back-Thin Fusion) in slit mode. The camera exposure was set at fast mode (2 ms) with 16b imaging. The *X*/*Y* sampling rate was 1.866 μm and tha *Z* step was set at 2 μm.

### AchE staining

The method for AchE staining method was as reported (Lim et al., 2004) with modifications as follows. Brain sections (40 μm) were cut and then mounted onto slides. The slides were incubated for 6 hr with solution A [0.0072% ethopropazine, 0.075% glycine, 0.05% cupric sulfate, 0.12% acetyl thiocholine iodide, 0.68% sodium acetate, (pH 5)] for enzymatic reactions. Slides were washed with ddH_2_O 3 times for 5 min and then were developed in solution B (0.77% sodium sulfide, pH 7.8) for 30 min. Next, slides were washed 3 times for 5 min and were exposed to solution C (1% silver nitrate) for 10 min in the dark for silver intensification and then rinsed for 3 times over 5 min. The slides were air-dried overnight and then were cleared in two changes of xylene followed by Permount mounting media.

### Saturation Binding assay

The whole brain cortices were collected from C57, *Htr2a*^EGFP-CreERT2^, and *Htr2a*^A242S-EGFP-Cre^ mice. Cortices was homogenized by a polytron homogenizer (BioSpec Products, Inc., Bartlesville, OK) on ice in 50 mM Tris buffer (pH7.4) with 0.1 mM PMSF and 1 mM EDTA. The homogenate was centrifuged at 1,000 x g for 10 min at 4°C. The supernatant was reserved for further centrifugation at 31,000 x g for 15 min at 4°C; pellets were collected as crude membranes. The crude membranes were suspended in 50 mM Tris (pH7.4) and 320 mM sucrose and were passed through a 26.5 G needle 5 times and then stored at - 80°C until use. The saturation binding assay was conducted using an 8-point dose-dependent concentration of [^3^H] ketanserin (maximal concentration 20 nM) (Perkin Elmer, Waltham, MA) with brain crude membranes (25 μg membrane proteins) with or without 10 µM clozapine in the binding buffer [50 mM Tris (pH 7.4) with 10 mM MgCl_2_]. The nonlinear saturation analysis was performed with GraphPad Prism 9.4.0 (GraphPad Software, La Jolla, CA) to calculate the Kd and Bmax values.

#### Quantitative reverse transcription-PCR

The whole brain cortices from C57, *Htr2a*^EGFP-CreERT2/+^, and *Htr2a*^EGFP-CreERT2/EGFP-CreERT2^ mice were dissected for total RNA purification. Cortices were homogenized with 1 ml Trizol (ThermoFisher) followed by chloroform (0.2 ml) extraction. The mixture was vortexed, and centrifuged at 12,000 x g for 5 min. The aqueous phase containing the RNA was transferred to a separate tube and 0.5 mL of isopropanol was added to precipitate the RNA. Samples were incubated on ice for 10 min and centrifuged at 12,000 x g at 4°C for 10 min. The supernatant was discarded, and the pellet was washed twice with 1 ml 80% ethanol. The RNA pellet was air-dried and re-suspended in 100 μl molecular-grade water. cDNA synthesis and PCR were performed using TaqMan™ Fast Virus 1-Step Multiplex Master Mix (#4331182; ThermoFisher). Probes were used to detect the mouse transcripts of *Htr2a* and *Gapdh* (Mm00555764_m1 and Mm99999915_g1 4448892, respectively; ThermoFisher). The input RNA (1 ng) was used at 20 μl per RT-qPCR reaction, and *Htr2a* and *Gapdh* signals were quantitated simultaneously using FAM and VIC fluorescence. Real-time PCR was performed with the CFX96 Real-Time PCR Detection System (Bio-Rad) and with Taqman probes. Primer specificity was confirmed by melting curve analyses. Reaction efficiencies over the appropriate dynamic range were calculated to ensure linearity of the standard curve. The results are expressed as fold-increase mRNA expression of the gene of interest normalized to *Gapdh* expression by the ΔΔ*C*_t_ method.

### HTR2A downregulation: drug treatment, receptor purification and western blot assay

C57 and *Htr2a*^EGFP-CreERT2^ mice were treated (i.p.) with vehicle or 0.5 mg/kg LSD for 5 days. Twenty-four hr later whole cortex was dissected for Western blot.

Receptor purification and subsequent immunoblotting were conducted according to our previously published procedure (Yadav et al., 2011). The whole cortex was homogenized with a Dounce homogenizer in membrane buffer [50 mM Tris (pH 7.4), 0.1 mM EDTA, 5% glycerol, and with protease and phosphatase inhibitors]. Tissue lysates were centrifuged at 1,000 x g for 10 min at 4°C. The supernatant was collected and centrifuged at 25,000 x g for 15 min at 4°C. The pellets were washed once with membrane buffer followed by re-centrifugation. Pellets (crude membranes) were lysed with lysis buffer [50 mM Tris (pH 7.4), 150 mM NaCl, 10% glycerol, 1% NP-40, 0.5% Na-deoxycholate, 0.5% CHAPS and protease inhibitors] for 1 hr on a rotating mixer at 4°C. Detergent-soluble proteins (supernatants) were collected following centrifugation at 12,500 x g for 20 min at 4°C. The supernatant was incubated at least 2 hr with 30 μl package volume of Wheat-germ (WGA) -conjugated agarose beads (Vector Laboratories, Inc., Burlingame, CA) on a rotating mixer at 4°C and washed 3 times with lysis buffer. WGA-bound proteins were eluted with 50 μl 2x Laemmli buffer [2% SDS, 0.25 M Tris (pH 7.4), 50% glycerol, and 0.01% Bromophenol Blue].

Samples were resolved with 4-12% SDS-PAGE (Life Technologies) and transferred onto polyvinylidene fluoride membranes. Membranes were incubated overnight with anti-rabbit-HTR2A antibody (1:500, Neuromics, #RA24288) at 4°C followed by horseradish peroxidase–conjugated goat anti-rabbit secondary antibodies (Jackson Immunoresearch) and finally reacted with enhanced chemiluminescence reagents. Images were captured with the ChemiDoc Image system (Bio-Rad Laboratories, Hercules, CA). Membranes were stripped and re-probed for β-actin (1:1000, Sigma, #A3854). Intensities of HTR2A were normalized against β-actin in the same lane. Samples from vehicle-treated *Htr2a*^+/+^ mice served as the control and were designated as 100%; other samples were normalized against it. The specificity of this HTR2A antibody has been confirmed in HTR2A KO mice in our previous study (Magalhaes et al., 2010).

### Behaviors

#### Open field activity

Locomotor, rearing, and stereotypical activities were examined in an open field (21 x 21 x 30 cm; Omnitech Electronics, Columbus, OH) under 180 lux illumination (Rodriguiz et al., 2021). Motor activities were assessed automatically by beam-breaks, all other behaviors were video-taped and scored by hand in a blinded fashion. Mice were placed into the open field for 30 min to collect baseline activities, they were injected (i.p.) with the vehicle, 0.3 mg/kg LSD, 1 mg/kg DOI, or 1 mg/kg psilocin, and returned immediately to the open field for 90 min. Locomotor activity (distance traveled), rearing (vertical beam-breaks), and stereotypical activities (repetitive beam-breaks less than 1 sec) were monitored in 5-min segments or as cumulative activities using Fusion Versamax software (5.3 Edition; Omnitech Electronics, Columbus, OH).

#### Head twitch responses (HTRs), grooming, and retrograde walking

These responses were filmed over 30 min during the open field studies following administration of drugs (Rodriguiz et al., 2021). Behavioral scoring for HTRs was conducted by researchers blinded to the sex, genotype, and treatment conditions of the mice. Scoring of grooming and retrograde walking were performed using the TopScan program (version 3; CleverSys Inc., Reston, VA). The results are expressed as the numbers of HTRs, duration of grooming (sec), and numbers of retrograde walking events. ***Prepulse inhibition (PPI)*** PPI of the acoustic startle response was conducted in SR-LAB apparati (San Diego Instruments, San Diego, CA) (Rodriguiz et al., 2021). Mice received (i.p.) the vehicle, 0.3 mg/kg LSD, 1 mg/kg DOI, or 1 mg/kg psilocin and were placed into the PPI chambers. Following habituation to a white-noise background (64 dB), testing began 10 min later. Each test was composed of 42 trials (6 null trials + 18 pulse-alone trials+ 18 prepulse-pulse trials). Null trials consisted of the white noise background. Pulse trials were composed of 40 msec bursts of 120 dB white noise (see below). Prepulse-pulse trials were comprised of 6 trials with the 20 msec prepulse stimuli (4, 8, and 12 dB above the white-noise background) that were followed 100 msec later with the pulse stimulus (120 dB). Testing commenced with 10 pulse-alone trials followed by combinations of the prepulse-pulse and null trials, and it ended with 10 pulse-alone trials. PPI was calculated as the %PPI = [1 - (pre-pulse trials/startle-only trials)] *100.

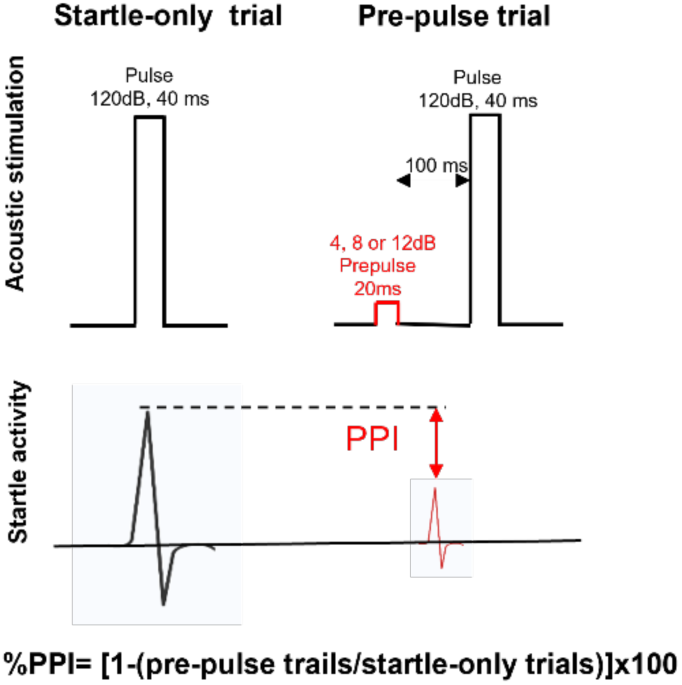

#### Electrophysiology in Htr2a^EGFP-CreERT2/+^xAi9 mice

*Htr2a*^EGFP-CreERT2/+^x Ai9 mice were rapidly decapitated, and the brains were placed into a beaker containing an ice-cold high sucrose solution (in mM): sucrose 206.0; KCl 2.5; CaCl_2_ 0.5; MgCl_2_ 7.0; NaH_2_PO_4_ 1.2; NaHCO_3_ 26; glucose 5.0; HEPES 5. Coronal sections (300 μm thickness) were sliced on a vibrating microtome (Leica VT1000S; Leica Microsystems, Buffalo Grove, IL) and incubated in oxygenated (95% O_2_/5% CO_2_) artificial cerebrospinal fluid (aCSF) composed of (in mM): NaCl 130; KCl 3.5; glucose 10; NaHCO_3_ 24; MgSO_4_-7H_2_O 1.5; NaH_2_PO_4_-H_2_O 1.26; CaCl_2_ 2.0 for 30 min at 37°C, followed by 30 min at RT (21-22°C) for equilibration. Patch pipettes (4-6 MΩ; King Precision Glass Inc., Claremont, CA) were filled with internal solution (mM: potassium gluconate 145; EGTA 5; MgCl_2_ 5; HEPES 10; Na-ATP 2; Na-GTP 0.2). Data were acquired with a Multiclamp 700B amplifier (Molecular Devices, Sunnyvale, CA), low-pass filtered at 2-5 kHz, digitized (Digidata 1550B; Molecular Devices), and stored using pClamp 10 software (Molecular Devices). Series resistance was continuously monitored with a hyperpolarizing 10 mV pulse; cells with axis resistance >20 MΩ were excluded from the data set. HTR2A expressing neurons were identified via tdTomato expression using fluorescent optics and brief (<2 sec) episodic illumination as previously described (Schmitz et al., 2022). During voltage clamp recording (V_hold_ = −70mV), electrophysiological properties of cells were determined by pClamp 10 Clampex software using a 10 mV pulse delivered after breaking the cellular membrane. Current Clamp recordings (I_hold_ = 70pA) were performed immediately afterwards to determine the firing type of each neuron included in the data sets.

#### hM3Dq DREAAD application in the *Htr2* ^Cre/+^ mice

##### Surgeries

Mice underwent intracranial surgeries for local viral injection [AAV-hSyn-DIO-eGFP or AAV-hSyn-DIO-HA-hM3D(Gq)-IRES-mCitrine (Addgene)] into the mPFC. Mice (7–9 weeks of age) were anesthetized with a mixture of ketamine (100 mg/kg) and xylazine (10 mg/kg), positioned in a small-animal stereotaxic instrument (David Kopf Instruments, Tujunga, CA) and the surface of the skull was exposed. Thirty-three gauge syringe needles (Hamilton) were used to bilaterally infuse 0.3 μl of virus into the mPFC (bregma coordinates: angle 10°, anteroposterior, +1.7; mediolateral, +/-0.8; dorsoventral, −2.4) at a rate of 0.1 μl/min. Needles were removed slowly over 5 min after infusions were complete.

##### Fixed tissue, perfusions/slicing/immunohistochemistry/imaging

Animals were transcardially perfused with a saline flush followed by 4% PFA and then further post-fixed in PFA overnight. Coronal sections (50 μm) were cut on a vibratome (Leica V1000s) in PBS. Sections were blocked with 10% normal donkey serum (NDS) and 2% bovine serum albumin (BSA) in PBS and 0.5% Triton-X100 (PBST) for 1 hr, then incubated with primary antibody in 10% NDS in PBST overnight at RT. Sections were rinsed 3 times for 30 min each at RT, followed by incubation in secondary antibody in 1% NDS in PBST at RT for 2 hr. Sections were then rinsed 3 times (10 min/rinse), mounted onto slides, cover-slipped with Fluoromount-G mounting medium (Southern Biotech, Birmingham, AL). Concentrations and sources for antibodies were as follows: chicken anti-GFP 1:1000 (Invitrogen) and secondary antibody 1:200 (Alexa Fluor 488). Image acquisition was performed with ZEN software on a Zeiss LSM 800 confocal system using 20X and 63X objectives. All image acquisition and post-processing settings were held constant to permit qualitative comparisons.

##### DREADD Electrophysiology

Mice were anesthetized with isoflurane and underwent intracardiac perfusions with ice-cold artificial cerebrospinal fluid (aCSF) which contained (in mM): NaCl 128, KCl 3, NaH_2_PO_4_ 1.25, D-glucose 10, NaHCO_3_ 24, CaCl_2_ 2, and MgCl_2_ 2 [oxygenated with 95% O_2_ and 5% CO_2_ (pH 7.4), 295–305 mOsm]. Acute coronal brain slices (200 μm) containing HTR2A^+^ mPFC neurons were prepared using a microslicer (Leica VT1200S) in ice-cold sucrose aCSF, which was derived by fully replacing NaCl with sucrose (254 mM) and saturated with 95% O_2_ and 5% CO_2_. Slices were maintained in holding chambers with aCSF for 1 hr at 32°C. Patch pipettes (3–5 mΩ) for whole-cell current-clamp recordings were pulled from borosilicate glass and filled with an internal solution containing (in mM): potassium gluconate 135, HEPES 10, KCl 4, MgATP 4, and NaGTP 0.3 (pH 7.31, 287 mOsm). Recordings from mPFC HTR2A^+^ neurons (identified visually by the presence of eYFP or mCitrine following AAVs injection into *Htr2a*^Cre/+^ mice) were performed in slices perfused with aCSF at 32°C (flow rate = 2.5 ml/min). Recordings were made using a Multiclamp 700B amplifier (Molecular Devices). Signals were digitized at 8 kHz using a Digidata 1440A (Molecular Devices), and filtered at 4 kHz. To measure the intrinsic membrane properties of mPFC HTR2A^+^ neurons, whole-cell recordings were conducted in current-clamp mode and series resistance (10–30 MΩ) was monitored throughout the recording session with neurons discarded if resistance changed by >20%. DCZ (250 nM) was bath-applied after 2.5 min of baseline recording and monitored for responses. Data were acquired and analyzed using pCLAMP11 (Molecular Devices). Firing rates and resting membrane potentials were analyzed during baseline and after bath application of DCZ.

### Statistics

All behavioral data analyses were performed with the IBM SPSS Statistics 27 programs (IBM, Chicago, IL). The results were presented as means and standard errors of the mean (SEMs). Student’s t-test, one- or two-way ANOVA, repeated measures ANOVA (RMANOVA), or analyses of covariance (ANCOVA) were used, and these analyses were followed by Bonferroni corrected pair-wise acomparisons. When Levene’s test of homogeneity of variabce was violated, the data were analysed by Mann-Whitney U or Wilcoxin tests for two groups or by Kruskal-Wallis tests for more than two groups, the latter was followed with Bonferroni tests. A *p*<0.05 was taken as significant. The results were graphed using GraphPad Prism (San Diego, CA). The results of the binding assay and western blot were analyzed by GraphPad Prism with t-test and two-way ANOVA, respectively. The electrophysiology results were analyzed by GraphPad Prism with paired t-tests and one-sample t-tests.

## Supplementary materials

**Fig S1.**
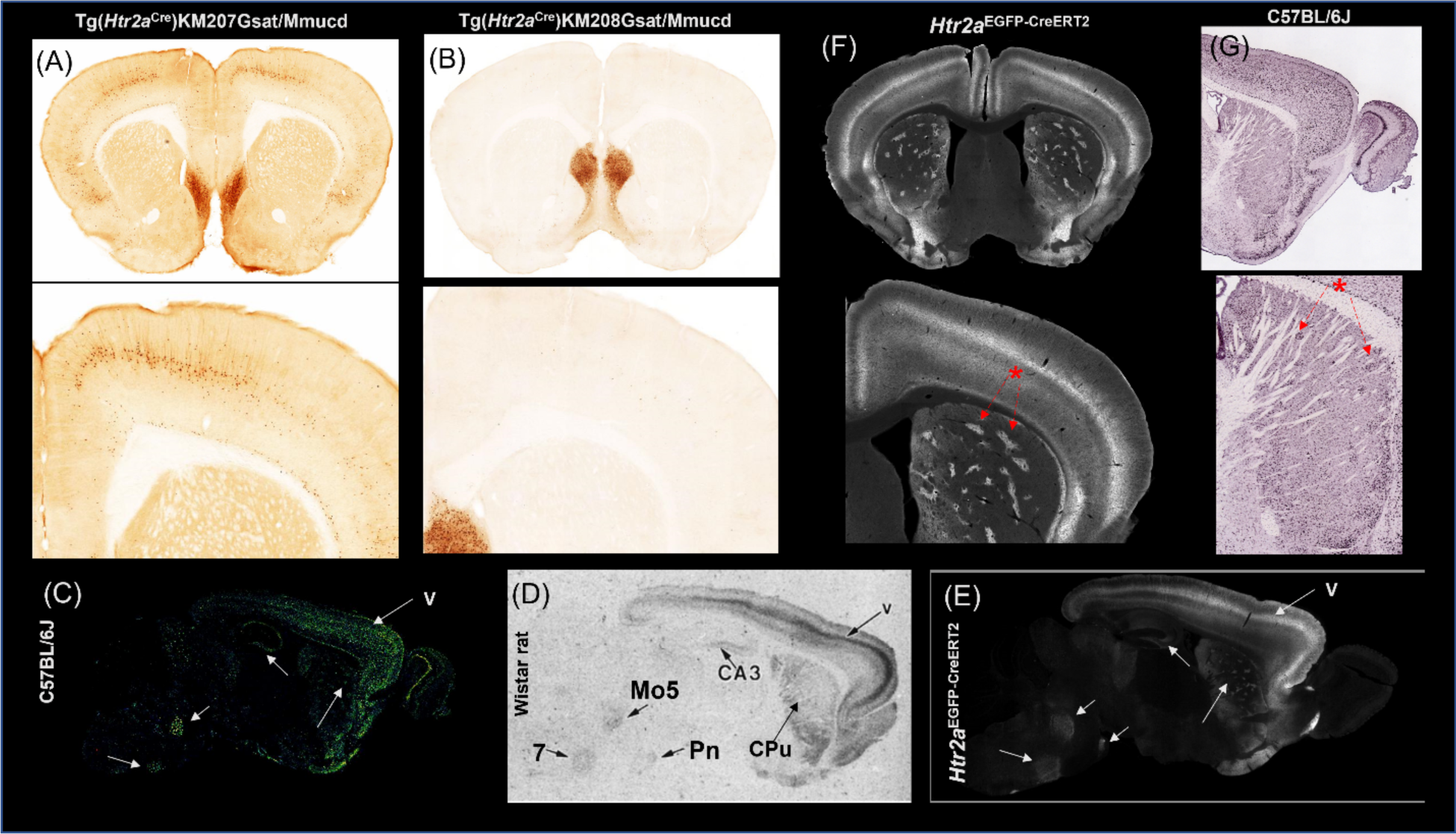
Comparison of HTR2A distributions with distinct approaches among different *Htr2a* mouse lines, C57 mouse and Wistar rat. **(A)/(B)** Coronal sections including the caudate putamen region from two *Htr2a*^Cre^ BAC mouse lines (KM207 and KM208) with Cre-dependent GFP reporter distribution: Image S1A is from http://www.gensat.org/imagenavigator.jsp?imageID=69385. Image S1B is from http://www.gensat.org/imagenavigator.jsp?imageID=83872. Comparison of *Htr2a* mRNA (Fig S1C), autoradiography (Fig S1D) and HTR2A-EGFP-CT protein (Fig S1E): arrows shown in these 3 figures indicated the corresponding brain regions with these three different methods. **(C)** Expression of *Htr2a* mRNA in the adult C57 mouse brain: Images are from Allen Mouse Brain Atlas (http://mouse.brainmap.org/experiment/siv?id=81671344). **(D)** [^3^H] MDL100,907 (0.4 nM) autoradiography labeling of HTR2A binding sites in the Wistar rat brain section (V: layer V cortex, CPu: caudate putamen, Pn: pon, 7: facial nucleus and M05: motor trigeminal nucleus.) (Lopez-Gimenez et al., 1997) **(E)** Sagittal view of distribution of HTR2A-EGFP-CT fusion protein in the whole brain cleared tissue from *Htr2a*^EGFP-CreERT2/+^ mouse line. Comparison of HTR2A-EGFP-CT protein and *Htr2a* transcripts in the CPu: The red star with dashed line with arrows in Fig S1F and S1G denoted the patch-like pattern (striosome) in the CPu. **(F)** The distribution of the HT2A-EGFP-CT fusion protein in coronal section at the level of the CPu from *Htr2a*^EGFP-CreERT2/+^ mouse line. **(G)** Distribution of *Htr2a* mRNA in the brain from C57 mice. Data resource from Allen Mouse Brain Atlas (http://mouse.brainmap.org/experiment/siv?id=81671344.

**Fig S2.**
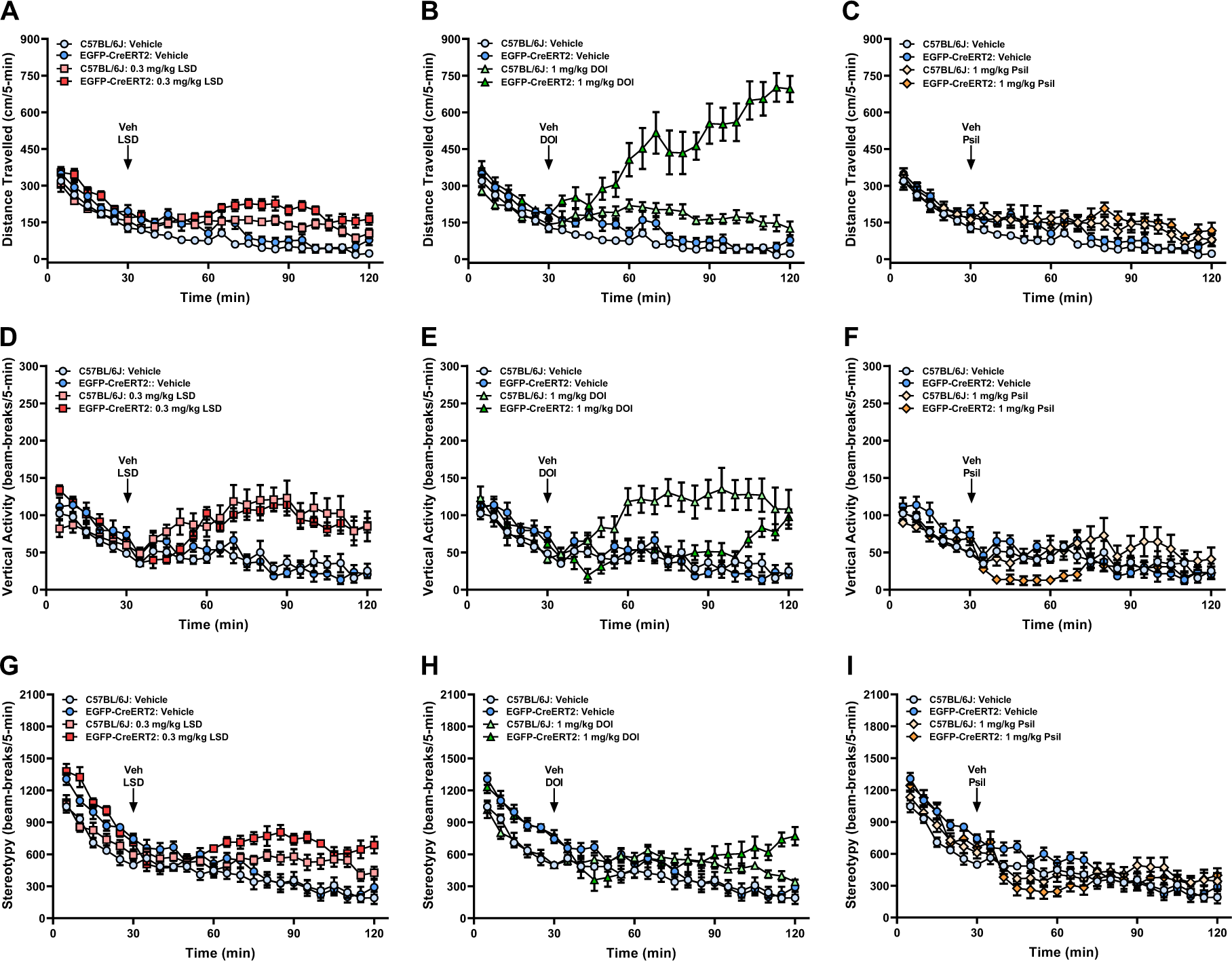
Pharmacological effects of psychedelics (LSD, DOI, and Psilocin) on motor activities in C57 and *Htr2a*^EGFP-CreERT2/EGFP-CreERT2^ mice. **(A-I) *Motor activities in the open field*** Baseline activities (0-30 min; pre-administration) and post-injection activities following administration (31-120 min) of the vehicle (Veh), LSD (0.3 mg/kg), DOI (1 mg/kg) or psilocin (Psil, 1 mg/kg); n=9-10 mice/genotype/treatment. Note, the C57BL/6J and *Htr2a*^EGFP-CreERT2/EGFP-CreERT2^ mice are termed C57 and *Htr2a*^EGFP-CreERT2^ mice, respectively, below and in the figure panels. All data represented as means ±SEMs; N=9-10 mice/genotype/treatment. (A-C) RMANOVA for baseline: time [F(23,1587)=23.922, *p*<0.001], time by genotype [F(23,1587)=8.900, *p*<0.001], time by treatment [F(69,1587)=14.677, *p*<0.001], time by genotype by treatment [F(69,1587)=7.859, *p*<0.001], genotype [1,69)=30.277, *p*<0.001], treatment [F(3,69)=24.114, *p*<0.001], and genotype by treatment interaction [F(3,69)=11.598, *p*<0.001]. RMANCOVA for post-injection: time [F(17,1071)=2.600, *p*<0.001], time by genotype [F(17,1071)=12.309, *p*<0.001], time by treatment [F(51,1071)=10.977, *p*<0.001], time by genotype by treatment [F(51,1071)=8.914, *p*<0.001], genotype [1,63)=16.278, *p*<0.001], treatment [F(3,63)=26.809, *p*<0.001], and genotype by treatment interaction [F(3,63)=8.972, *p*<0.001]. (D-F) Rearing activities in the same mice. RMANOVA for baseline: time [F(23,1587)=23.548, *p*<0.001], time by genotype [F(23,1587)=9.221, *p*<0.001], time by treatment [F(69,1587)=7.544, *p*<0.001], time by genotype by treatment interaction [F(69,1587)=1.840, *p*=0.020], genotype [1,69)=5.228, *p*=0.025], treatment [F(3,69)=12.397, *p*<0.001]. RMANCOVA for post-injection: time by genotype [F(17,1071)=2.670, *p*<0.001], time by treatment [F(51,1071)=6.033, *p*<0.001], time by genotype by treatment [F(51,1071)=1.699, *p*=0.002], genotype [1,63)=15.125, *p*<0.001], treatment [F(3,63)=11.016, *p*<0.001], and genotype by treatment interaction [F(3,63)=4.016, *p*=0.011]. (G-I) Stereotypical activities in these same mice. RMANOVA for baseline: time [F(23,1587)=139.510, *p*<0.001], time by genotype [F(23,1587)=7.485, *p*<0.001], time by treatment [F(69,1587)=5.973, *p*<0.001], time by genotype by treatment interaction [F(69,1587)=2.485, *p*<0.001], genotype [1,69)=7.491, *p*=0.008], and treatment [F(3,69)=7.632, *p*<0.001]. RMANCOVA for post-injection: time by genotype [F(17,1071)=3.353, *p*<0.001], time by treatment [F(51,1071)=7.105, *p*<0.001], time by genotype by treatment interaction [F(51,1071)=3.910, *p*<0.001], and treatment [F(3,63)=7.226, *p*<0.001].

**Fig S3.**
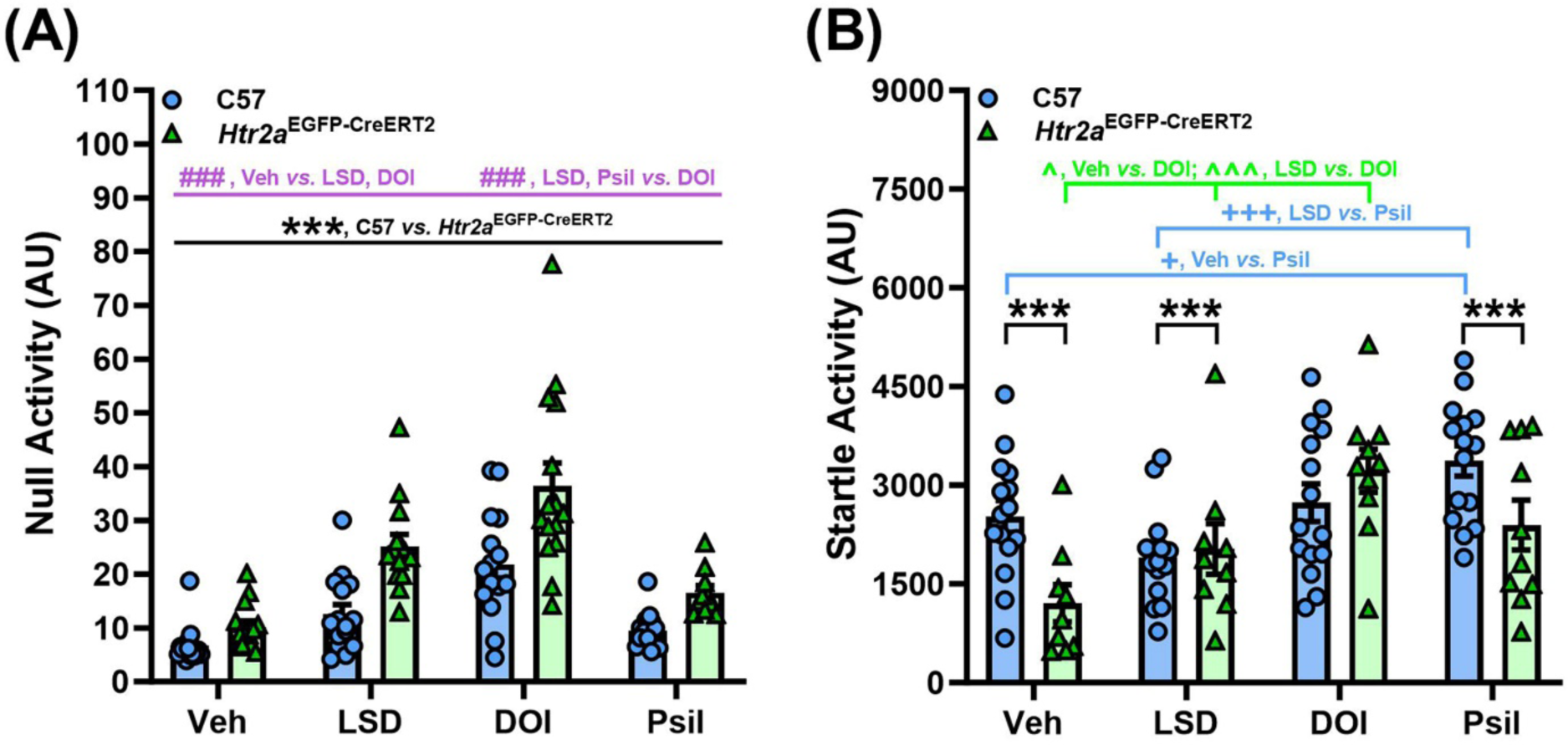
Pharmacological effects of psychedelics (LSD, DOI, and Psilocin) on null and startle activities in C57and *Htr2a*^EGFP-CreERT2/EGFP-CreERT2^ mice. **(A-B) *Null and startle activities in prepulse inhibition*** Mice were injected with the Veh, 0.3 mg/kg LSD, 1 mg/kg DOI, or 1 mg/kg Psil and tested 10 min later; n=10-15 mice/genotype/treatment. Note, the C57BL/6J and *Htr2a*^EGFP-CreERT2/EGFP-CreERT2^ mice are termed C57 and *Htr2a*^EGFP-CreERT2^ mice, respectively, below and in the figure panels. The results are presented as means ± SEMs; n=10-15 mice/genotype/treatment. (A) Null activities in C57 and *Htr2a*^EGFP-CreERT2^ mice. Two-way ANOVA: genotype [F(1,105)=33.886, *p*<0.001] and treatment [F(3,105)=32.101, *p*<0.001]; Bonferroni *post-hoc* tests: for C57 *vs*. *Htr2a*^EGFP-CreERT2^ mice (black): \*\*\**p*<0.001, overall genotype effects; overall treatment effects (purple): ^###^*p*<0.001, Veh *vs.* LSD and DOI or LSD and Psil *vs.* DOI. (B) Startle activities in C57 and *Htr2a*^EGFP-CreERT2^ mice. Two-way ANOVA: genotype [F(1,105)=51.786, *p*<0.001], treatment [F(3,105)=12.810, *p*<0.001], and genotype by treatment interaction [F(3,105)=3.282, *p*=0.024]; Bonferroni *post-hoc* tests: for C57 *vs*. *Htr2a*^EGFP-CreERT2^ mice (black): \*\*\**p*<0.001, Veh, LSD, and Psil; for C57 (blue): ^+^*p*<0.05, Veh *vs.* Psil; ^+++^*p*<0.001, LSD *vs.* Psil; for *Htr2a*^EGFP-CreERT2^ mice (green): ^*p*<0.05, Veh *vs.* DOI; ^^^*p*<0.001, LSD *vs.* DOI.

**Fig S4.**
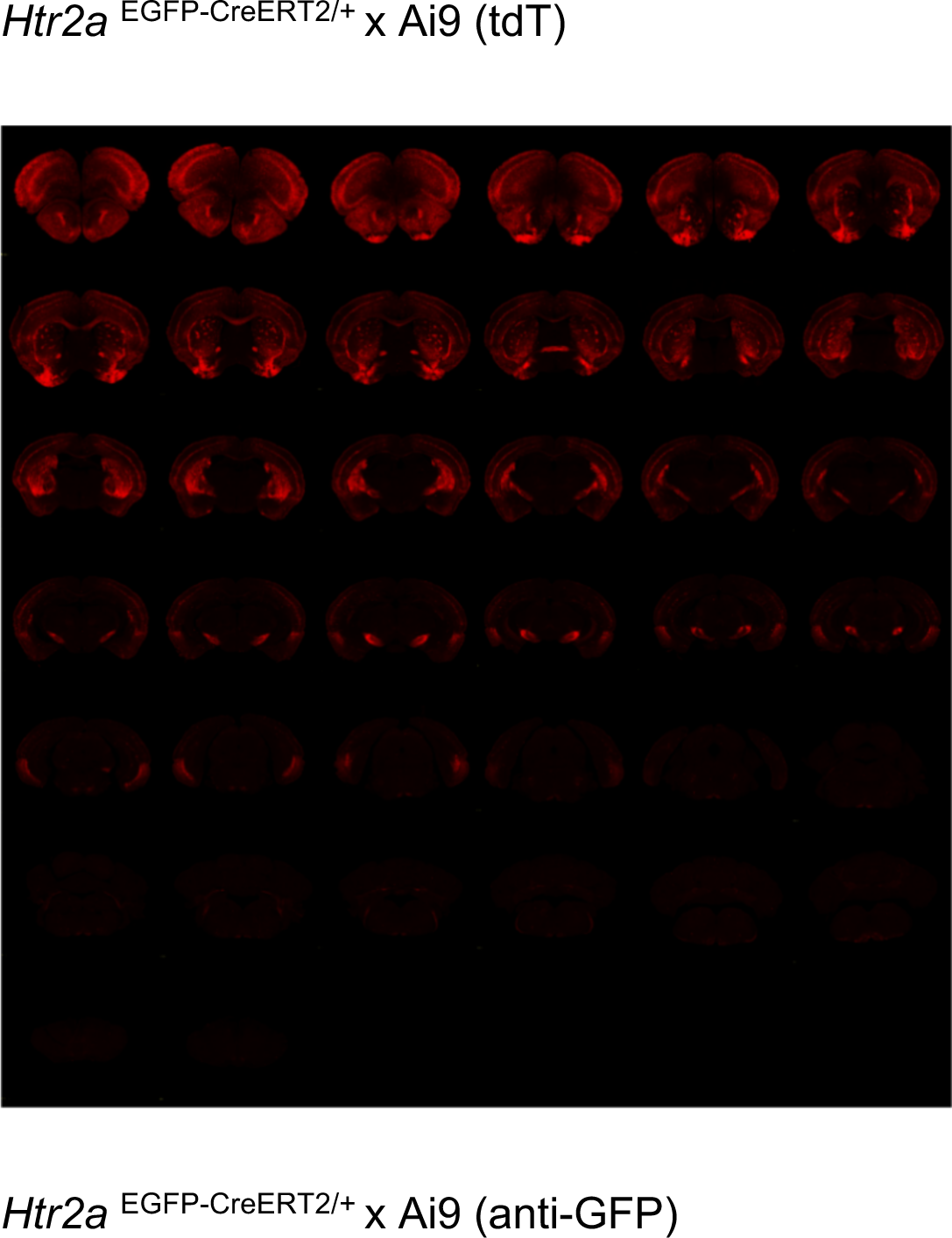
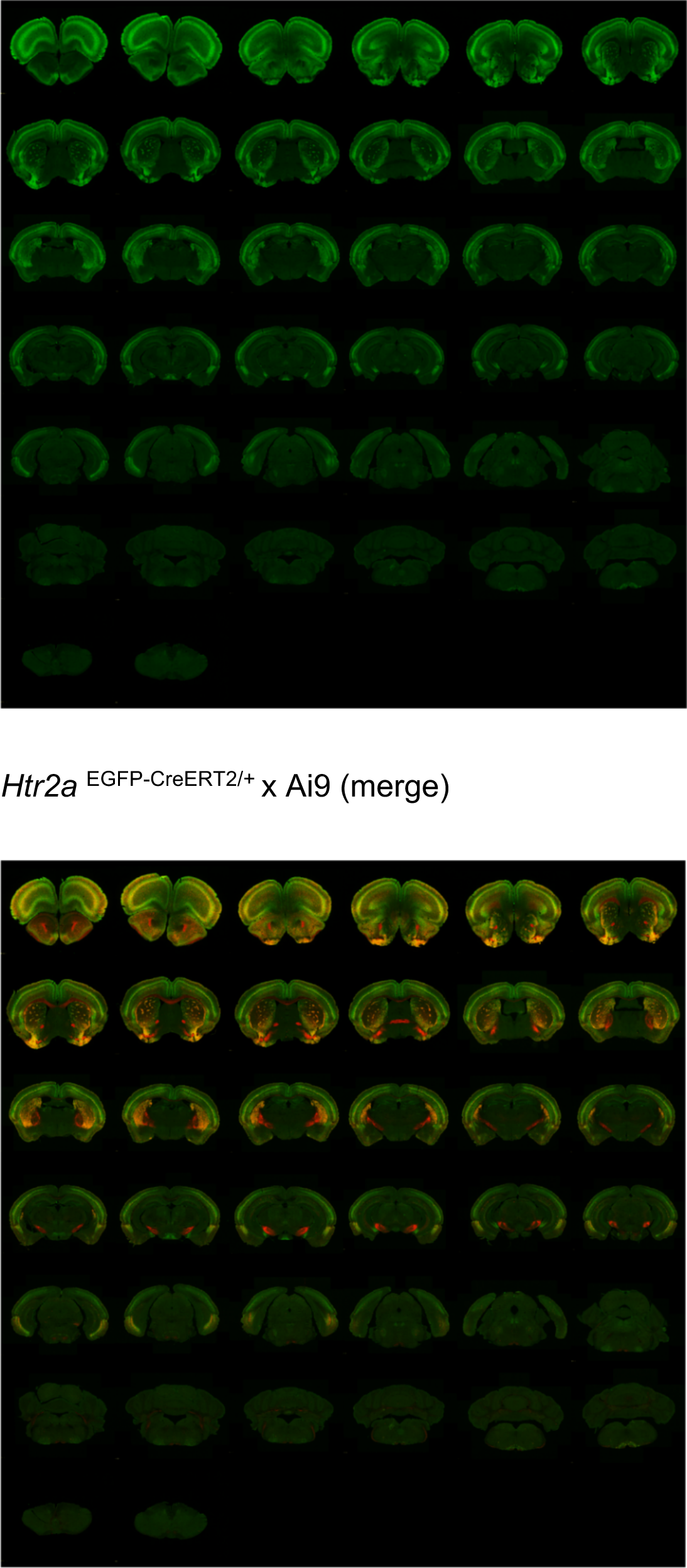
Whole brain mapping in *Htr2a* ^EGFP-CreERT2/+^ x Ai9 mice: *Htr2a* ^EGFP-CreERT2/+^ mice were crossed with Ai9 mice. Mice were treated with tamoxifen (100 mg/kg, i.p.) at p39-p42 for 4 days and then brains were perfused at 14 days post-injection (14 dpi) at p56. The brain sections were stained with GFP antibody. Images were captured in two channels for tdTomato (red) and GFP (green) signals under an Olympus slide scanner with 10X objective. The GFP signal as a HTR2A-EGFP-CT fusion protein and the tdTomato signal as HTR2A-positive cells were shown here. Experiments were conducted on 4 brains with similar results. The raw images have been uploaded to open-source website, A Mouse Imaging Server (AMIS, https://amis2.docking.org/).

### Movies S1. 3D rendering view

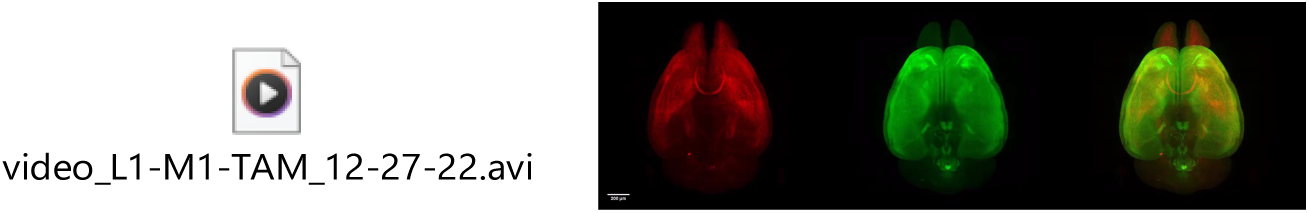

### Movie S2: coronal view

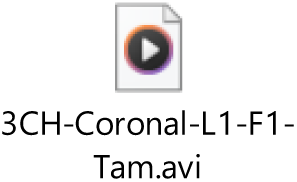

### Movie S3: horizontal view

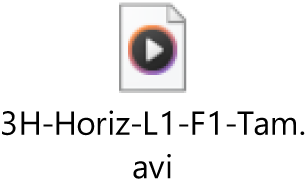

### Movie S4: sagittal view

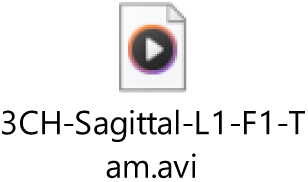

### Whole brain cleared tissues from *Htr2a* ^EGFP-CreERT2/+^ x Ai9 mice

*Htr2a* ^EGFP-CreERT2/+^ x Ai9 mice were treated with tamoxifen (100 mg/kg, i.p.) at p39-p42. Brains were perfused at 14 dpi and then were conducted for tissue clarity (see the details in the materials and methods). The whole cleared brains were scanned under a light-sheet microscope with 3.6X objective. The whole brain was stained with GFP antibody. Signals of tdTomato (Red) for HTR2A positive cells and GFP (Green) for HTR2A-EGFP-CT fusion protein were captured. Three dimensional (3D) rendering are depicted (Movie S1), coronal view (Movie S2), horizontal view (Movie S3) and sagittal view (Movie S4), respectively. The movies have been uploaded to open-source website, A Mouse Imaging Server (AMIS, https://amis2.docking.org/).

**Fig S5.**
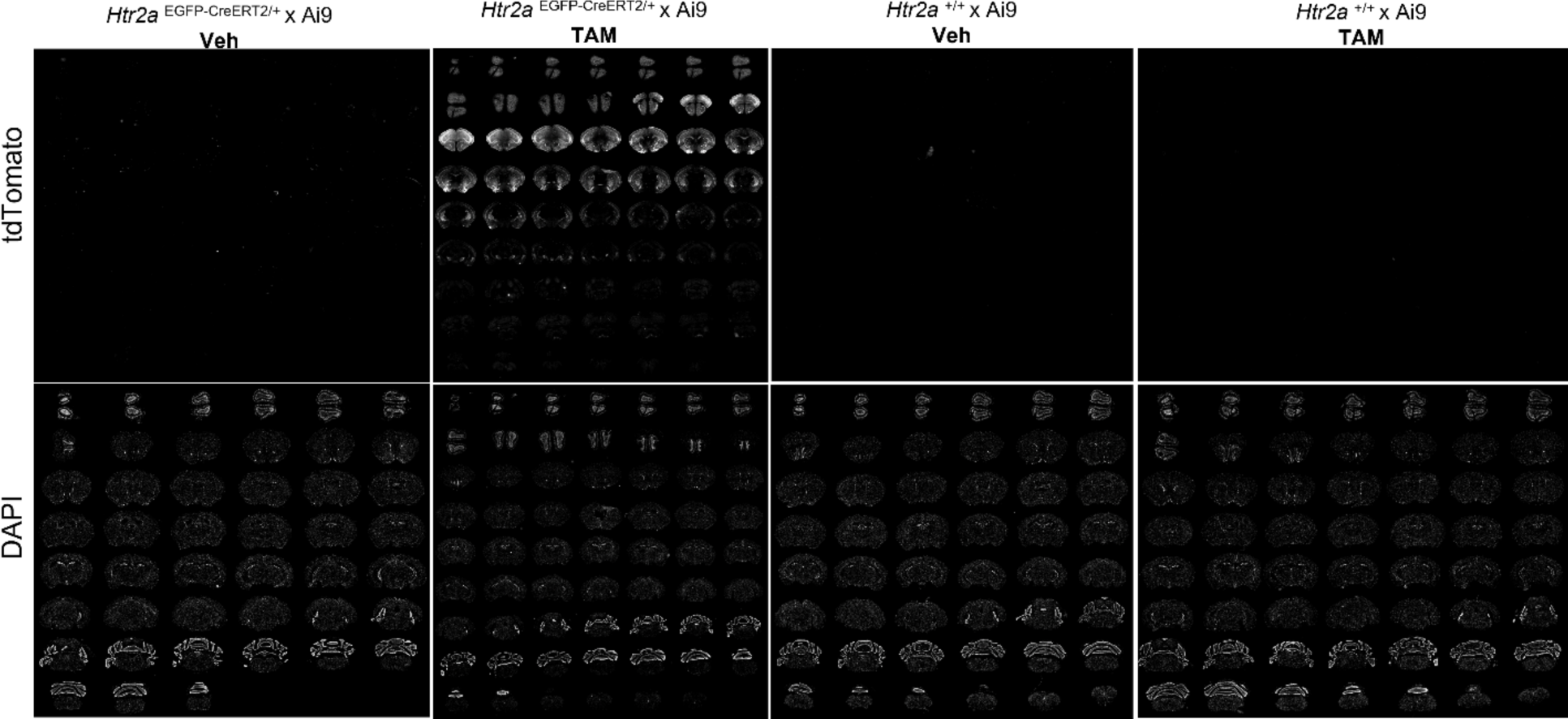
The activity and specificity of CreERT2 recombinase were characterized in in *Htr2a* ^EGFP-CreERT2/+^ x Ai9 mice and C57 x Ai9 (*Htr2a*^+/+^x Ai9) mice. *Htr2a* ^EGFP-CreERT2/+^ x Ai9 or C57 x Ai9 (*Htr2a* ^+/+^ x Ai9) mice were treated with vehicle (corn oil) or tamoxifen (100 mg/kg) at p39 to p42 with 4 injections and then brains were collected at p56. A whole brain mapping was conducted to determine whether CreERT2 recombinase turned on tdTomato protein expression after TAM treatment. The brain sections were stained with DAPI and then images were rendered by an Olympus VS120 slide scanner. Experiments were conducted with 4 animals with similar results.

**Fig S6.**
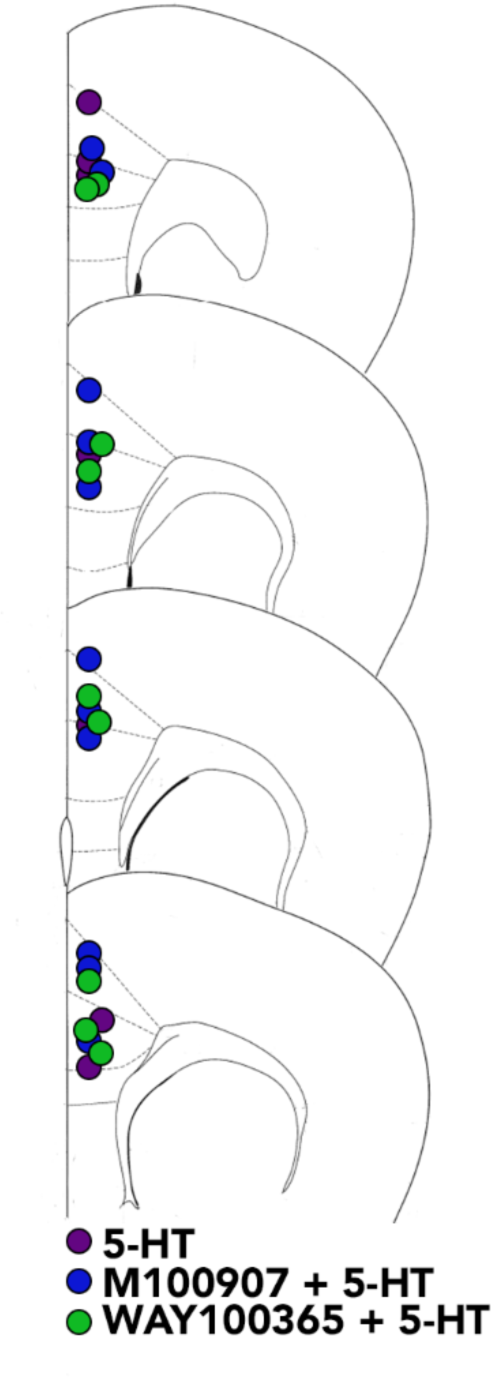
Schema showing the location of HTR2A-tdTomato expressing neurons used in the electrophysiological recordings. The recording neurons were located in the mPFC (ACC or dPL) from *Htr2a*^EGFP-CreERT2/+^ x Ai9 mice. *Htr2a*^EGFP-CreERT2/+^ x Ai9 mice were used for this experiment. Individual recording neurons (HTR2A-tdTomato-positive) from different brain sections of each treated group (purple: 5-HT; blue: M100907+5-HT; and Green: WAY100365+5-HT) are depicted in the ACC or dPL. (A) HTR2A signals with anti-HTR2A staining from *Htr2a*^Cre/+^ mouse (B) HTR2A*-*A242S-EGFP-CT signals with anti-GFP staining from *Htr2a* ^A242S-EGFP-Cre/+^ (C-D) Comparison of different Cre-dependent reporters (Ai9 mice vs PHP.eB.flex.tdT AAV virus) among *Htr2a* mouse lines

**Fig S7.**
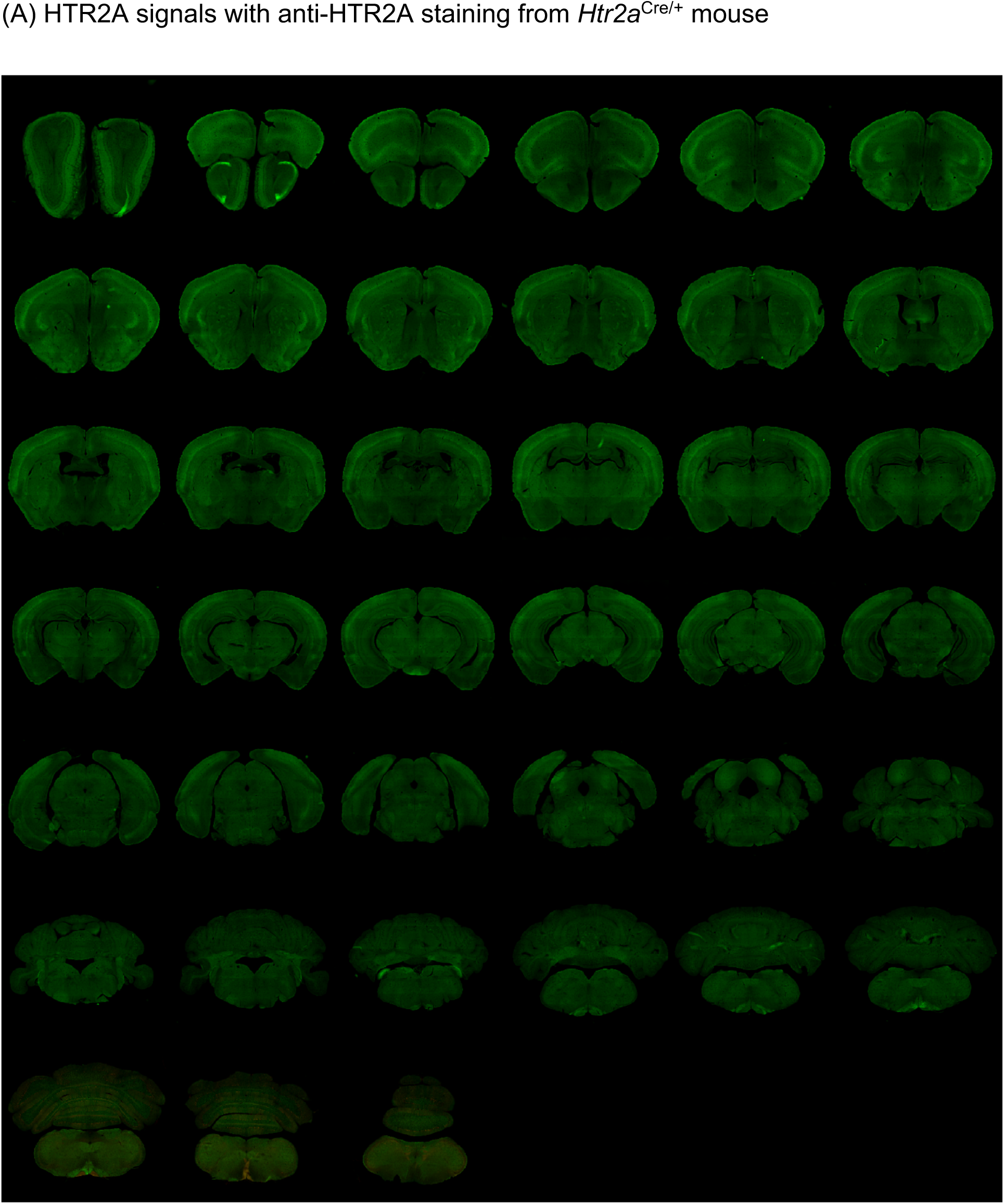
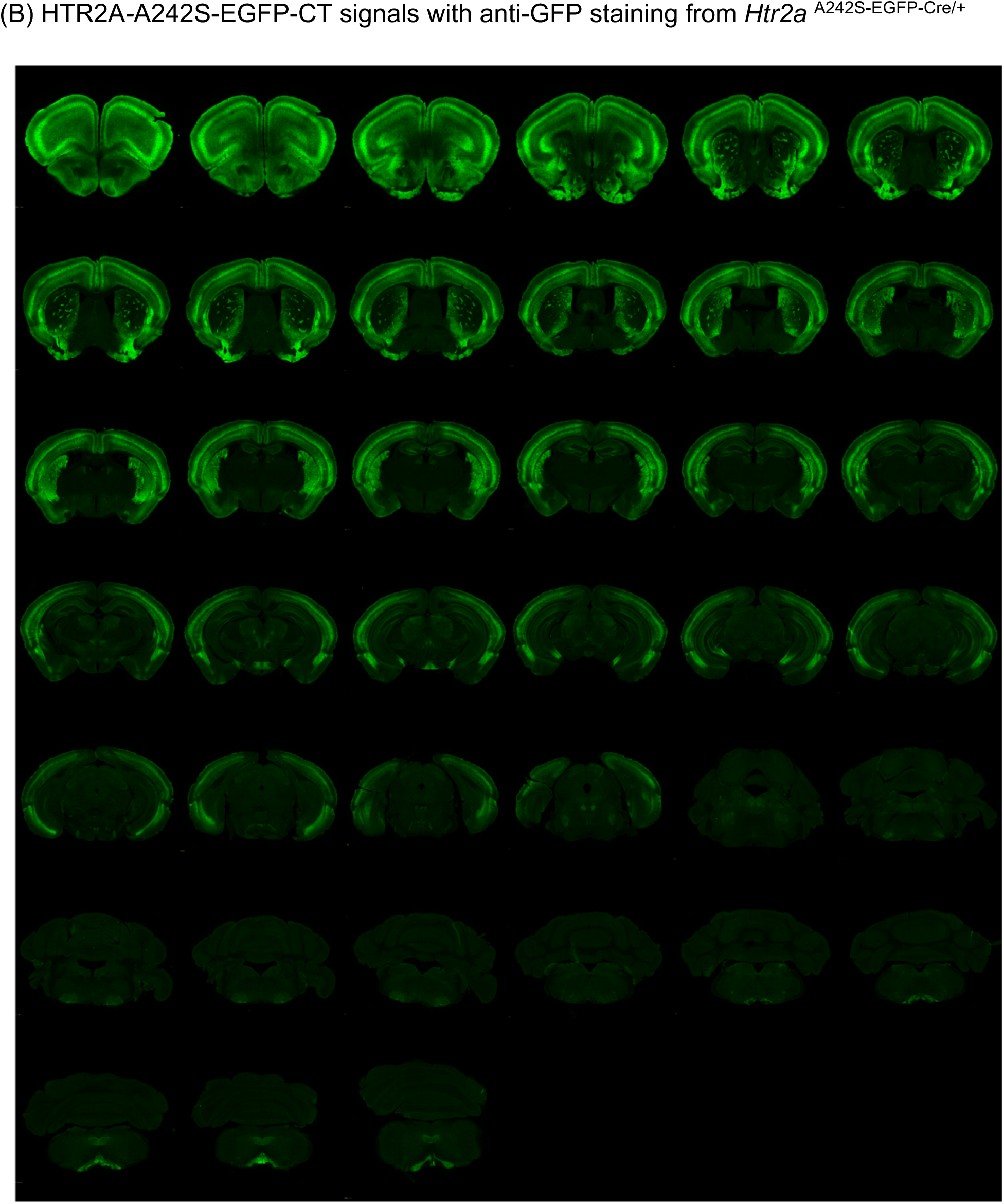
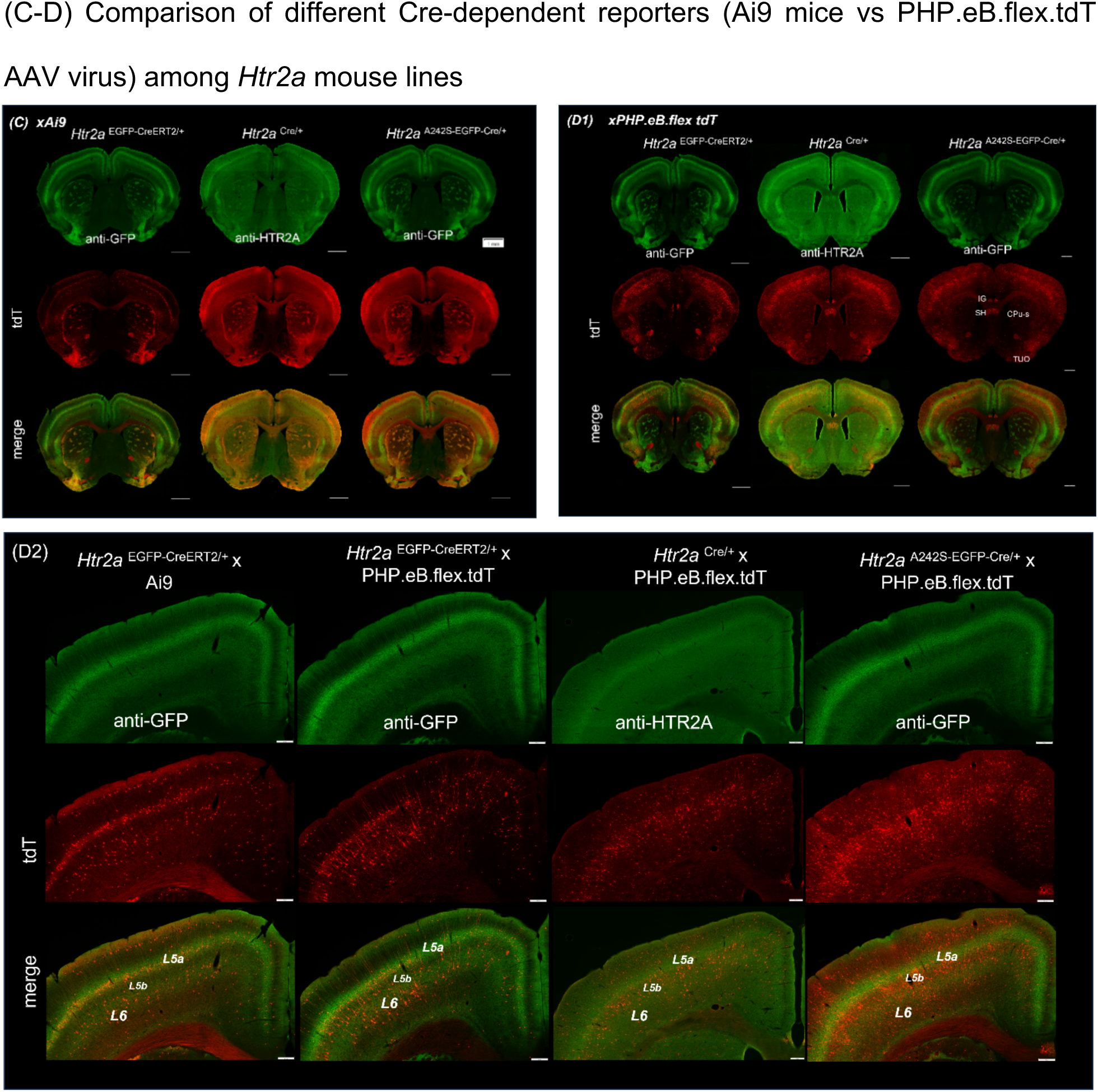
HTR2A distribution in multiple *Htr2a* mouse lines: **(A)** The HTR2A proteins in the whole brain mapping from *Htr2a* ^Cre/+^ mice were shown using an anti-HTR2A antibody. **(B)** The distribution of HTR2A-A242S-EGFP-CT fusion proteins from *Htr2a* ^A242S-EGFP-Cre/+^ mice were mapped in the whole brain using an anti-GFP antibody. The raw images have been uploaded to open-source website, A Mouse Imaging Server (AMIS, https://amis2.docking.org/). **(C)** Cre reporter mice (Ai9) crossed with three *Htr2a* mouse lines. The distribution patterns of HTR2A and HTR2A^+^ tdTomato from an inducible *Htr2a*^EGFP-CreERT2/+^ x Ai9 mouse and from constitutive Cre lines (*Htr2a*^Cre/+^ xAi9 and *Htr2a*^A242S-EGFP-Cre/+^ x Ai9) are shown. All mice were perfused at p56. The brain sections were stained with anti-HTR2A or anti-GFP antibodies. Images were captured in two channels for tdTomato (red) and anti-HTR2A or anti-GFP (green) signals under and Olympus slide-scanner with 10X objective. Experiments were conducted with 3-4 brains with similar results. **(D)** Cre reporter virus (PHP.eB.flex.tdT) with inducible CreERT2 and constitutive Cre-mouse Htr2a lines. The *Htr2a* mouse lines were retro-orbitally injected with PHP.eB.flex.tdTomato (Fig S7D1). The tdTomato patterns of expression among the 3 mouse lines from virally-treated groups are compared with *Htr2a*^EGFP-CreERT2/+^ x Ai9 mice in the area of the somatosensory cortex (Fig S7D2). Mice were perfused and then stained with anti-HTR2A for *Htr2a*^Cre/+^mice and with anti-GFP for *Htr2a*^EGFP-CreERT2/+^ and *Htr2a*^A242S-EGFP-Cre/+^ mice. Images were captured under an Olympus slide-scanner with 10X objective. (CPu-s: striosome, TUO: olfactory tubercle, IG: induseum griseum, SH: septohippocampal nucleus).

**Fig S8.**
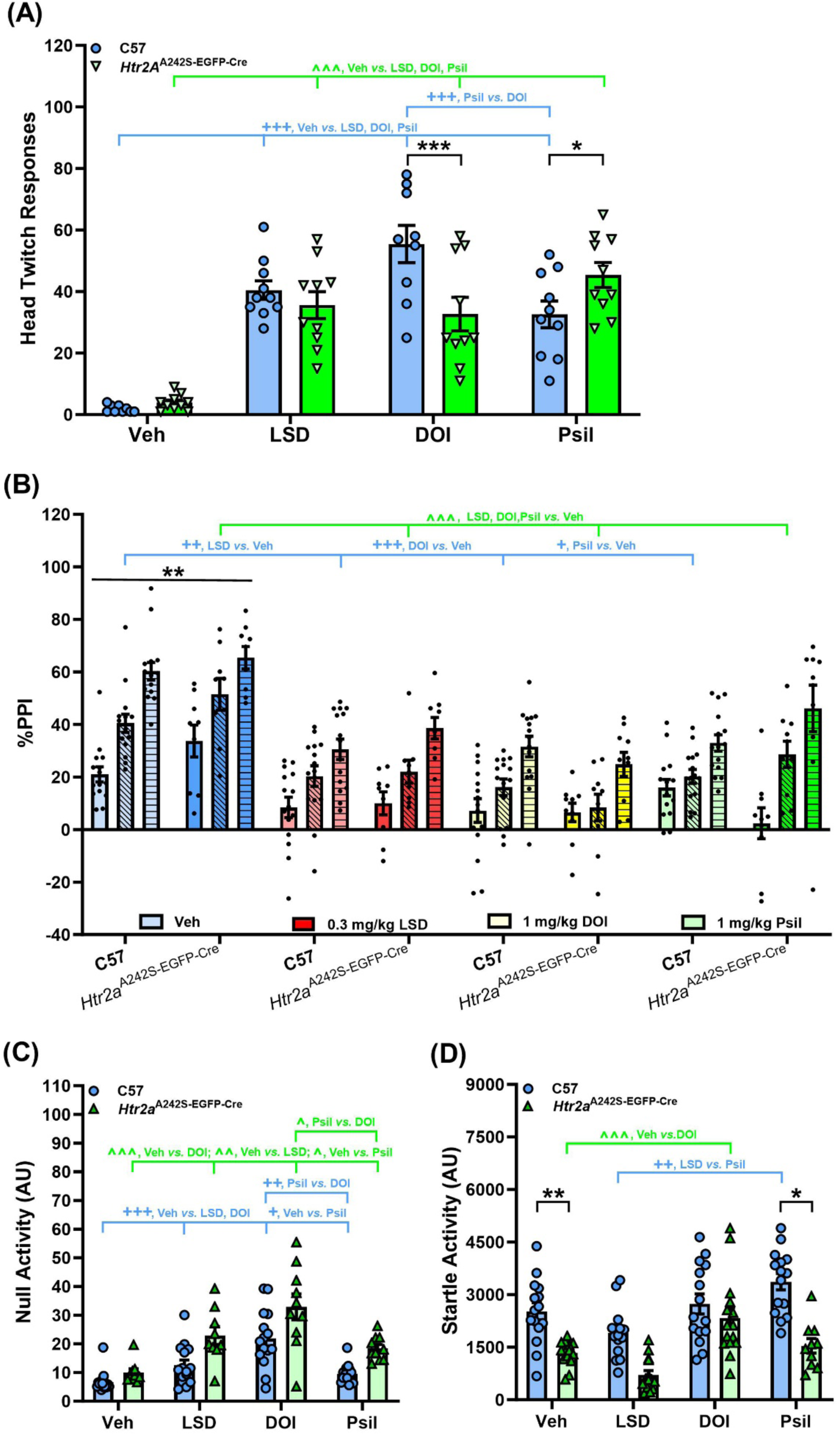
Pharmacological effects of psychedelics (LSD, DOI, and Psilocin) on head twitch responses in C57 and *Htr2a*^A242S-EGFP-CreE/A242-EGFP-Cre^ mice. **(A-D) *Head twitch responses and prepulse inhibition (PPI)*** Head twitch responses were scored over 30 min beginning after injection of the vehicle (Veh), LSD (0.3 mg/kg), DOI (1 mg/kg), or Psil (1 mg/kg). For PPI, mice received the same regimen of treatment and were tested in PPI 10 min later as described in the Methods section. Note, the _C57BL/6J and *Htr2a*_^A242S-EGFP-Cre/A242S-EGFP-Cre^ _mice are termed C57 and *Htr2a*_^A242S-EGFP-Cre^ mice, respectively, below and in the figure panels. All data are represented as means ± SEMs. (A) Head twitch responses in C57 and *Htr2a*^A242S-EGFP-Cre^ animals immediately following administration of the Veh, LSD, DOI, or Psil over the 30 min following administration; n=9-10 mice/genotype/treatment. Two-way ANOVA: treatment [F(3,70)=43.033, *p*<0.001], and genotype by treatment interaction [F(3,70)=6.784, *p*<0.001]; Bonferroni *post-hoc* tests: for C57 *vs. Htr2a*^A242S-EGFP-Cre^ mice (black): \**p*<0.05, Psil; \*\*\**p*<0.001, DOI; for C57 mice (blue): ^+++^*p*<0.05, Veh *vs.* all psychedelics or Psil *vs.* DOI; for *Htr2a*^A242S-EGFP-Cre^ mice (green): ^^^*p*<0.05, Veh *vs*. all psychedelics. (B) Prepulse inhibition: C57 and *Htr2a*^A242S-EGFP-Cre^ mice were injected with the Veh, LSD, DOI, or Psil and tested 30 min later; n=9-10 mice/genotype/treatment. RMANOVA: PPI [F(2,140)=113.340, *p*<0.001], genotype [F(1,70)=4.989, *p*=0.029], and treatment [F(3,70)=23.203, *p*<0.001]; Bonferroni *post-hoc* tests: for C57 vs *Htr2a*^A242S-EGFP-Cre^ mice (black): \*\**p*<0.01, Veh; for C57 mice (blue): ^+^*p*<0.05, Psil *vs.* Veh; ^++^*p*≤0.001, LSD *vs.*Veh; ^+++^*p*≤0.001, DOI *vs.* Veh; for *Htr2a*^A242S-EGFP-Cre^ mice (green): ^^^*p*<0.001, all psychedelics *vs.* Veh. (C) Null activities in C57 and *Htr2a*^A242S-EGFP-Cre^ mice. In the two-way ANOVA Levene’s test of homogeneity was violated and it was still violated when null activity was standardized to genotype or vehicle. Neither the Mann-Whitney U test nor the Wilcoxin W test for genotype (two-tailed) were significant for any treatment. Kruskal-Wallis one-way ANOVA for treatment (two-tailed): [h(3)=27.582, *p*<0.001]; Bonferroni *post-hoc* tests for C57 mice (blue); ^+^*p*<0.05; Veh *vs.* Psil; ^++^*p*<0.01, Psil *vs.* DOI; ^+++^*p*<0.001, Veh *vs.* LSD and DOI; for *Htr2a*^A242S-EGFP-Cre^ mice (green): ^^^*p*<0.05; Veh *vs.* Psil or Psil *vs.* DOI; ^^^^*p*<0.01, Veh *vs.* LSD; ^^^^^*p*<0.001, Veh *vs.* DOI. (D) Startle activities in C57 and *Htr2a*^A242S-EGFP-Cre^ mice. Two-way ANOVA: genotype [F(1,70)=4.295, *p*=0.042], treatment [F(3,70)=7.547, *p*<0.001], and genotype by treatment interaction [F(3,70)=3.740, *p*=0.015]; ANOVA followed by Bonferroni *post-hoc* tests: for C57 *vs*. *Htr2a*^A242S-EGFP-Cre^ mice (black): \**p*<0.05, Psil; \*\**p*<0.01, Veh; for C57 (blue): ^++^*p*<0.05, LSD *vs.* Psil; for *Htr2a*^A242S-EGFP^ mice (green): ^^^*p*<0.001, Veh *vs.* DOI.

**Fig S9.**
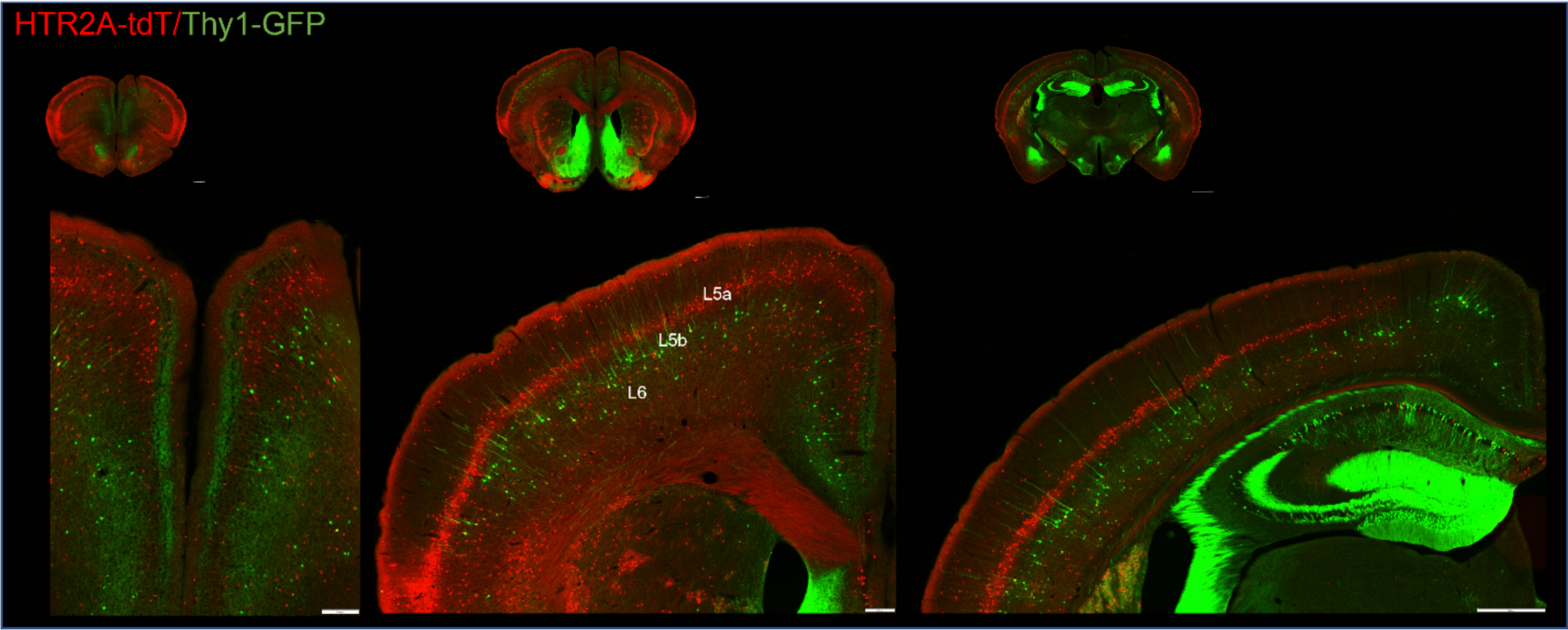
The distribution of HTR2A-containing neurons and Thy1-expressing neurons. *Htr2a*^EGFP-CreERT2/+^ x Ai9 x *Thy1*^GFP/+^ mice were used and were treated with tamoxifen (100 mg/kg for 4 days) to induce tdTomato expression. Since HTR2A-EGFP-CT is very dim without antibody amplification, it was difficult to directly detect. Thus, in this experiment we used tdTomato to label HTR2A-containing neurons and Thy1-GFP to observe their distribution in the cortex. Images were captured under an Olympus slide-scanner with 10X objective. Experiments were conducted on 3 brains with similar results.

**Fig S10.**
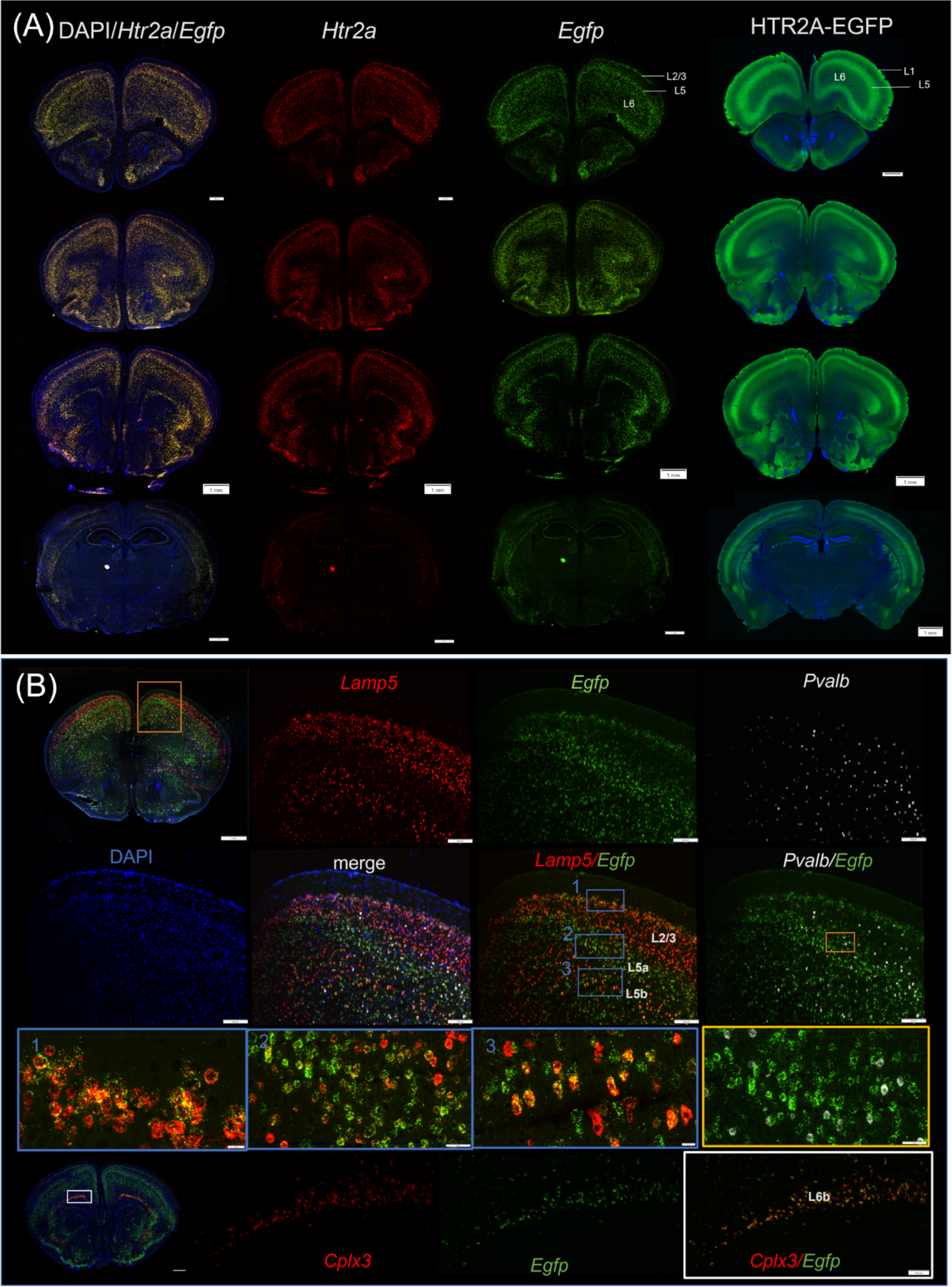
*Htr2a* mRNA distribution. (A) RNAscope was used to detect *Htr2a* and *Egfp* transcripts in the *Htr2a*^EGFP-CreERT2/EGFP-CreERT2^ mice. Brain sections were also stained with anti-GFP antibody to detect protein signals of the HTR2A-EGFP-CT fusion protein. Experiments were conducted with 4 animals containing similar results. Images were taken under a 10X objective with an Olympus VS120 slide-scanner (scale bar 1mm). (B) In the motor cortex area, *Egfp* transcripts were co-localized with *Lamp5* in Layer2/3 and Layer5b, and with *parvalbumin* positive interneurons. *Cplx3* as a layer6b marker showed its co-localization with *Egfp* transcripts. Images were taken under 10X objective with an Olympus VS120 slide-scanner and under a 20X objective with an Olympus FV3000RS confocal microscope. Experiments were conducted with 3 animals achieving similar results.

## Abbreviations

ACC: anterior cingulate cortex
AHA: anterior hypothalamic area
AON: anterior olfactory nucleus
APT: thalamic anterior pretectal nucleus
ARC: arcuate nucleus
AUD: primary auditory cortex
AV: anteroventral thalamic nucleus
BLN: basolateral amydalar nuclei
BMN: basomedial anydalar nuclei
BPN: basilar pontine nuclei
CA1d: dorsal hippocampus, CA1
CA1v: ventral (temporal) hippocampus, CA1
CA2d: dorsal hippocampus, CA2
CA2v: ventral (temporal) hippocampus, CA2
CA3d: dorsal hippocampus, CA3
CA3v: ventral (temporal) hippocampus, CA3
CA4d: dorsal hippocampus, CA4
CA4v: ventral (temporal) hippocampus, CA4
CeA: central amygdalar nucleus
CERc: cerebellar cortex
CERdn: cerebellar deep nuclei
CL: claustrum
CM: thalamic central medial nucleus
Cpu: striatum (caudatoputamen)
CPu-m: striatum (caudatoputamen), matrix
CPu-s: striatum (caudatoputamen), striosome
DI: dysgranular cortex
DG: dentate gyrus
dHIP: dorsal hippocampus
DMN: dorsomedial nucleus
dlPN: dorsolateral pontine nucleus
DR: dorsal raphe nucleus
DTN: dorsal tegmental nucleus
ECT: ectorhinal cortex
END: endopyriform cortex
FP: frontal pole
GP: globus pallidus
GUS: gustatory cortex
hDBB: diagonal band of Broca, horizontal limb
IC: inferior colliculus
ICdg: inferior colliculus, deep gray layer
ICig: inferior colliculus, intermediate gray layer
ICsg: inferior colliculus, superficial gray layer
IL: infralimbic cortex
IO: inferior olivary nuclei
IPN: interpeduncular nucleus
K-F: Kolliker-Fuse nucleus
*l*ENT: entorhinal cortex, lateral
LAN: lateral amygdala nucleus
LDN: thalamic laterodorsal nucleus
LGN: thalamic lateral geniculate nucleus
LH: lateral hypothalamic area
LHb: lateral habenula nucleus
LMN: lateral mammillary nucleus
*l*NAc: nucleus accumbens, lateral
LO: lateral orbital cortex
LOT: lateral olfactory tract
LSN: lateral septal nucleus
*l*PBN: parabrachial nucleus, lateral
*l*POA: preoptic area, lateral
*l*TUO: olfactory tubercle, lateral
M1: primary motor cortex
M2: secondary motor cortex
MeA: medium anygdalar nucleus
MD: thalamic mediodorsal nucleus
mENT: medial entorhinal cortex
mHB: medial habenula nucleus
MGN: thalamic medial geniculate nucleus
MMN: medial mammillary nucleus
mNAc: nucleus accumbens, medial
MO: medial orbital cortex
MOF: motor nucleus of the facial (VII)
MOT: motor nucleus of the trigeminal (V)
mPOA: preoptic area, medal
mPBN: parabracial nucleus, medial
MR: median raphe nucleus
MSN: medial septal nucleus
mTUO: olfactory tubercle, medial
mVES: medial vestibular nucleus
NA: nucleus ambiguus
NAc: nucleus accumbens
ND: nucleus Darkschewitsch
NRTP: nucleus reticularis tegmenti pontis
PAG: periaqueductal gray
pAIC: pregenual anterior insular cortex
pgACC: pregenual anterior cingulate cortex
PF: thalamic parafascicular nucleus
PHA: posterior hypothalamic area
RE: thalamic reuniens nucleus
RHM: thalamic rhomboid nucleus
PFA: perifornical area
PIR: piriform cortex
PL: prelimbic cortex
PVT: thalamic paraventricular nucleus
RM: raphe magnus nucleus
RN: red nucleus
RRF: retrorubral field
RSP: retrospenial cortex
RT: thalamic reticular nucleus
sCIN: supracollosal cingulate cortex
SC: superior colliculus
SCdg: superior colliculus, deep gray layer
SCig: superior colliculus, intermediate gray layer
SCsg: superior colliculus, superficial gray layer
SMC: somatomotor cortex
SN: substantia nigra
SPF: thalamic subparafascicular nucleus
SSC: somatosensory cortex
STN: subthalamic nucleus
SUBd: subiculum, dorsal hippocampus
SUBv: subiculum, vental hippocampus
TMN: tuberomammillary nucleus
TRN: tegmental reticular nucleus of Bechterew
TT: tenia tecta
TUO: olfactory tubercle
VA/VL: thalamic ventral anterior/ventrolateral nuclei
vDBB: diagonal band of Broca, vertical limb
VIS: primary visual cortex
VLO: ventrolateral orbital cortex
VM: ventromedial thalamic nucleus
VMN: ventromedial nucleus
VP: ventral pallidum
VPN: ventral pontine nuclei
VTA: ventral tegmental area
ZI: zona incerta

## Distribution of the HTR2A-EGFP-CT in the whole brain

### Telencephalon

#### Frontal cortices

The HTR2A receptor, as reflected by HTR2A-EGFP-CT, was widely expressed in cortical regions from the frontal pole to the occipital lobe. In most of the neocortex the HTR2A-EGFP-CT was densely expressed in cells distributed in a characteristic laminar pattern, prominently expressed in the superficial aspects of layer 5 (L5).

HTR2A-EGFP-CT fusion proteins were present in all cortical regions of the frontal pole (rostral to infralimbic cortex, as defined by Van de Werd and colleagues (Van De Werd and Uylings, 2008) (Suppl. Figure 1A and 1B) but were most dense in the rostral orbitofrontal (OFC) and the somatomotor cortices. Within the various cytoarchitectonic regions comprising the rostral OFC, the dorsal agranular insular cortex was the most densely invested with HTR2A-EGFP-CT, closely followed by lateral and ventrolateral orbital cortices.

HTR2A-EGFP-CT in the frontal pole formed a continuous band that extended from the anterior cingulate medially to sweep laterally across the motor and somatomotor cortices before penetrating the dysgranular and agranular insular cortices. This band of HTR2A-EGFP-CT continued parallel to the pial surface of the OFC, through the lateral and ventral orbital cortices before turning dorsally to traverse the medial orbital cortex and finally the prelimbic area before returning to the anterior cingulate cortex and encircling the frontal pole.

In contrast to the frontal pole, the general distribution of HTR2A-EGFP-CT in the medial prefrontal cortex--which contains a defined infralimbic cortex (IL, areas 25)--was similar to that seen in the frontal pole with one exception. Here, the L5a band was not present in the IL or in the ventral prelimbic cortex (PL, area 32) (Suppl. #1 Figure C). Ventral to the IL, the medial orbital cortex also contained HTR2A-EGFP-CT.

There are no consistent cytoarchitectonic features that distinguish dorsal and ventral prelimbic cortex. However, the dorsal PL was more densely invested than its ventral counterpart with a greater density of acetylcholinesterase (AChE) -positive fibers (Van De Werd and Uylings, 2008). We therefore examined adjacent tissue sections stained to reveal AChE and HTR2A-EGFP-CT. The ventral (AChE-weak) zone of the PL did not express HTR2A-EGFP-CT but the dorsal PL did (Suppl.#1 Figure C).

#### Other cortical areas

As in the frontal cortices, HTR2A-EGFP-CT in areas posterior to the genu of the callosum formed an arching band in L5a. HTR2A-EGFP-CT extended from the cingulate cortex medially to the rhinal sulcus laterally. The HTR2A-EGFP-CT in the postgenual cortex were most dense in L5a but were present also at lower density in L6, with fewer still in L2/3. HTR2A-EGFP-CT labeling of apical dendritic arborizations was seen in L1.

Ventral to the rhinal sulcus there was a modest density of HTR2A-EGFP-CT that marked the piriform cortex. Caudal to the piriform cortex, the density of HTR2A-EGFP-CT increased in the perirhinal cortex and was very dense in the lateral, but not the medial entorhinal cortex. At this level the intensity of HTR2A-EGFP-CT staining was low-to-moderate in the primary auditory and visual cortices and it was present also in secondary visual and auditory cortical areas.

#### Anterior olfactory areas

There was a moderate plexus of HTR2A-EGFP-CT-positive elements in the anterior olfactory nucleus, particularly in the ventral aspects. The anterior-most piriform cortex was labeled. Interestingly, HTR2A-EGFP-CT was present in low-to-moderate densities in the lateral olfactory tract embedded in the ventrolateral aspect of the olfactory bulb, but it was very dense in the lateral olfactory tract axons that course to and enter the olfactory bulb.

#### Claustrum and endopiriform cortex

The claustrum exhibited moderate HTR2A-EGFP-CT signal through most of its anteroposterior extent, although labeling was more pronounced in the central core of the claustrum. Ventral to the claustrum, HTR2A-EGFP-CT was present in the ventrally contiguous endopiriform nucleus and in the cortex dorsally adjacent to the claustrum.

#### Striatal complex

Both the dorsal striatum (caudatoputamen) and the ventral striatum (nucleus accumbens and olfactory tubercle) exhibited areas of dense HTR2A-EGFP-CT signal. In the dorsal striatum, small clusters of intense HTR2A-EGFP-CT staining were embedded in a much less dense background area. This pattern was similar to the striatal patch-matrix system, in which small islands of weak acetylcholinesterase (AChE) termed striosomes were intermixed with a surrounding matrix compartment enriched in AChE. We therefore examined sections stained with a µ-opioid receptor (MOR) antibody that selectively stains striosomes (Herkenham and Pert, 1981). MOR and HTR2A-EGFP-CT were co-localized in very densely filled striosome-like compartments (Figure 1H). In addition, the subcallosal stria, a thin (∼50 µm) band of striatal tissue that is an elongated striosome running along the ventral surface of the callosum, was both MOR- and HTR2A-EGFP-CT-positive (Figure 1H).

The density of HTR2A-EGFP-CT expression in the matrix (MOR-negative) compartment was observed to differ across the rostrocaudal extent of the dorsal striatum. In the rostral striatum, HTR2A-EGFP-CT-enriched striosomes were present over a diffuse light background of HTR2A-EGFP-CT matrix staining, while more posteriorly in the mid-striatum densely-stained HTR2A-EGFP-CT-positive striosomes were seen in a matrix almost devoid of HTR2A-EGFP-CT staining. However, still more caudal, in the tail of the striatum, it was difficult to differentiate GFP in striosomal patches from the background HTR2A-EGFP-CT matrix staining.

In the rostral half of the nucleus accumbens, HTR2A-EGFP-CT densely distributed the ventrolateral shell and core regions but was much less dense in the medial shell and septal pole areas. More posteriorly, the medial shell was almost devoid of HTR2A-EGFP-CT, while the lateral shell and core were very densely filled with fibers. These distinctions in intensity of HTR2A-EGFP-CT labeling did not correspond to labeling in different accumbal compartments--such as the core and shell--but rather adopted a simply a lateral-to-medial gradient across the accumbens.

The intensity of HTR2A-EGFP-CT in the olfactory tubercle (TUO) paralleled that seen in the accumbens, with the marker of HTR2A-EGFP-CT expression being less dense in the medial than lateral TUO. The lateral TUO was among the most densely-stained areas in the mouse brain.

#### Pallidal areas

There was no significant HTR2A-EGFP-CT of either the globus pallidum or the ventral pallidum.

#### Septum, amygdala, and hippocampus

The septum and diagonal band complex lacked significant HTR2A-EGFP-CT. This led to a forebrain staining pattern in which dense GFP staining in the dorsal and ventral striatum flanked the septum and diagonal band, which lacked significant GFP staining.

The amygdala displayed a heterogenous pattern of HTR2A-EGFP-CT, with most of the amygdala exhibiting very low or no HTR2A-EGFP-CT while the extreme lateral and medial nuclei were moderately-to-densely distribution. HTR2A-EGFP-CT was dense in the lateral nucleus, but the ventromedially-adjacent basolateral nucleus was devoid of HTR2A-EGFP-CT. There was almost no HTR2A-EGFP-CT in the central nuclei, with weak staining of the posterior basomedial and cortical nuclei. The medial amygdalar nucleus was densely filled with HTR2A-EGFP-CT.

In the dorsal hippocampus there was trace expression of HTR2A-EGFP-CT in the pyramidal cell areas and the dentate gyrus. In the ventral (temporal) hippocampus there was a very different pattern, with almost no HTR2A-EGFP-CT in the CA1 and CA2 fields but very intense staining of the ventral aspects of the CA3 field with GFP-labeled pyramidal cell dendrites visible in the stratum oriens. The dentate gyrus did not express significant HTR2A-EGFP-CT.

### Di- and mes-encephalon

The HTR2A-EGFP-CT was widely distributed in the forebrain, with particular enrichment in cortical areas and the striatal complex. In contrast, HTR2A-EGFP-CT expression appeared to be progressively less as one moved from the forebrain to diencephalic, midbrain and pontomedullary areas.

#### Thalamus and epithalamus

Most of the thalamus lacked significant HTR2A-EGFP-CT.

There were two exceptions to the paucity of the HTR2A-EGFP-CT in the thalamus. The subparafascicular nucleus was filled with a moderate density of GFP fibers, while a light plexus of GFP staining filled the anterior pretectal nuclei.

In the epithalamus, the lateral habenula was invested with a light-to-moderate density of HTR2A-EGFP-CT-positive fibers, while in the medial habenula significant HTR2A-EGFP-CT expression was restricted to the dorsomedial tip of the structure. There was no appreciable HTR2A-EGFP-CT in the paraventricular nucleus.

#### Hypothalamus

HTR2A-EGFP-CT was found in only a few hypothalamic areas. The ventromedial nucleus displayed light-to-moderate GFP staining, with trace levels in the dorsomedial nucleus. The zona incerta was lightly filled with HTR2A-EGFP-CT.

In the posterior hypothalamus, HTR2A-EGFP-CT was seen in the mammillary bodies. Little HTR2A-EGFP-CT was present in the lateral mammillary nucleus and the ventral premammillary nucleus, but very dense staining was present in the medial and lateral divisions of the mammillary area, as well as, in the dorsal but not ventral tubero-mammillary nuclei.

#### Mesencephalon

Trace or low levels of diffuse HTR2A-EGFP-CT signal were seen in a numbers of midbrain areas, including the mesencephalic periaqueductal gray, the red nucleus, and the nucleus of Darkschewitz. The density of the HTR2A-EGFP-CT, as reflected by GFP, was low in the interpeduncular nucleus (IPN). There was slightly greater expression in the lateral and dorsal IPN subnuclei than the central and intermediate IPN nuclei (as defined by (Lenn and Hamill, 1984).

Dorsally in the midbrain, low HTR2A-EGFP-CT labeling was present in the superior colliculus, with the greatest density in the intermediate gray layer. In the intermediate gray staining inhomogeneities contributed to a patchy appearance. GFP staining density was less intense in the inferior colliculus.

Weak GFP expression was observed starting in the mesencephalic periaqueductal gray and extending caudally into the rostral pontine periaqueductal gray.

### Rhombencephalon

#### Pons

Across the basilar pons, areas of dense or very dense GFP staining were interdigitated with weak or absent HTR2A-EGFP-CT. In the dorsomedial and dorsolateral nuclei (Mihailoff et al., 1981) staining was dense, while the ventral pontine cells displayed much weaker staining. In addition, there was moderate staining in the tegmental reticular nucleus (of Bechterew) dorsally.

HTR2A-EGFP-CT was moderately dense in the dorsal tegmental nucleus; the surrounding central gray showing significantly less intense staining. In the parabrachial complex, the dorsal parabrachial and Kolliker-Fuse nuclei were weakly positive for HTR2A-EGFP-CT, with trace amounts of the fluorophore observed in the ventral parabrachial nuclei. The paralemniscal nucleus showed light-to-moderate GFP, and the nucleus ambiguus was lightly stained.

Prominent by virtue of absence was HTR2A-EGFP-CT expression in the raphe nuclei. Thus, there was no consistent staining of the dorsal or median raphe nuclei or of the raphe obscurus, while trace HTR2A-EGFP-CT was seen around the lateral aspects of the raphe magnus.

Cranial nerve nuclei exhibited a moderate density HTR2A-EGFP-CT, prominently including the fifth (V, trigeminal) and seventh (VII, facial) cranial nerve nuclei.

#### Medulla

A few medullary areas expressed HTR2A-EGFP-CT. Most prominent among these was the inferior olivary complex (IO). Although staining throughout the IO was moderate-to-dense, there were subnuclear variations in GFP density, including a (relatively) high density of staining in subnucleus A of the medial nucleus, moderate staining in the principal nucleus, and less intense staining in the cap of Kooy of the medial nucleus [following the nomenclature of (Yu et al., 2014)].

#### Cerebellum

No specific HTR2A-EGFP-CT was observed in either the cerebellar cortex or the deep nuclei.

#### White matter

Despite the dense expression of HTR2A-EGFP-CT in L5 cortical neurons, which gave rise to extensive descending myelinated projections as well as some interhemispheric connections, there was no significant GFP labeling of the corpus callosum. Similarly, neither the anterior commissure, nor the internal capsule were labeled. However, in the forebrain the lateral olfactory tract was lightly labeled. By contrast, the lateral olfactory tract outside of the corpus of the brain was very intensely labeled.

More posteriorly in the brain several myelinated fiber areas expressed some HTR2A-EGFP-CT, although in no case was such staining dense. In the posterior thalamus, the fasciculus retroflexus exhibited light-to-moderate HTR2A-EGFP-CT and could be traced from the habenula to the interpeduncular nucleus.

In addition, low levels of HTR2A-EGFP-CT were observed in the medial accessory oculomotor tract, and a small number of moderately-stained HTR2A-EGFP-CT-positive axonal clusters were visible in the lateral lemniscus. Finally, moderate HTR2A-EGFP-CT was present in the axons of the facial nerves.

**Supplementary#1-Figure.**
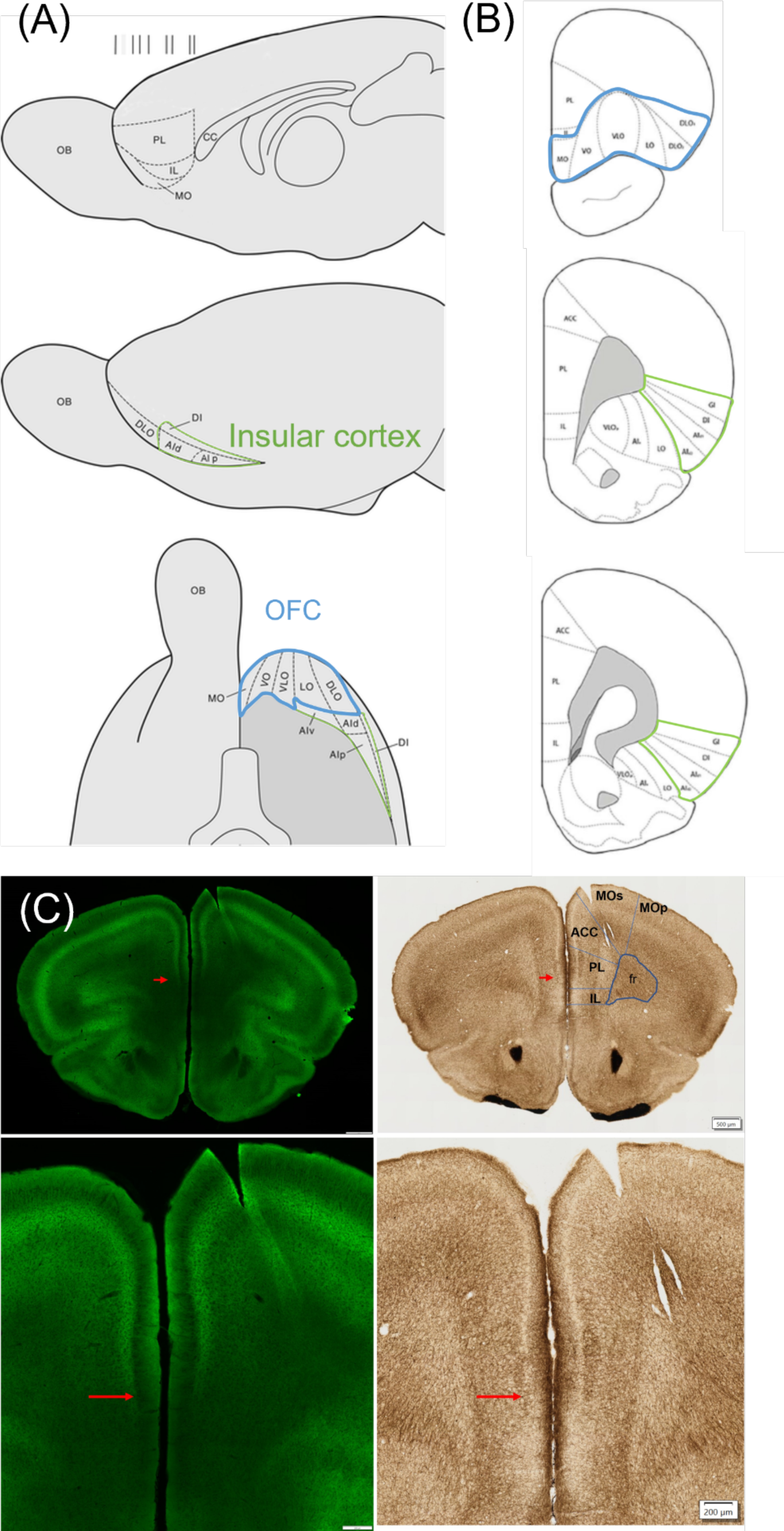
Characterization of HTR2A-EGFP-CT expression patterns in different brain regions of mice. **(A/B)** Schematic representation of the cytoarchitectonic areas of the frontal lobe of the brain. **(A)** The sagittal view of medial and lateral brain region, as well as horizontal view of the ventral regions on the frontal lobe show mPFC (medial prefrontal cortex), OFC (orbitofrontal cortex, as ventral PFC), and insular cortex (dorsal and ventral agranular insular cortex, AId and AIv; posterior agranular insular cortex, AIp; dysgranular insular cortex, DI). The schema was adapted from (Van De Werd and Uylings, 2008) **(B)** Coronal sections from rostral to caudal view for mPFC, OFC, and insular cortex. The schema was adapted from (Murphy and Deutch, 2018) **(C)** AchE staining: sequential brain sections through the frontal lobe area were used to visualize the distribution of HTR2A-EGFP-CT in the mPFC. The cytoarchitecture features of AchE staining showed strongest staining in the dorsal prelimbic cortex that was absent in the ventral prelimbic cortex. The blue arrow indicates the boundary between the dorsal and ventral prelimbic cortex. HTR2A-EGFP-CT showed its distributions in the dorsal prelimic cortex but was not present in the ventral prelimbic cortex and infralimbic cortex. Images scale bar: 500 µm. Similar results were obtained with 3 animals. MOs: secondary motor cortex, MOp: primary motor cortex, AAC: anterior cingulate area, PL: prelimbic cortex, IL: infralimbic cortex, and fr: anterior forceps.

